# Mapping adipocyte interactome networks by Halotag-enrichment-mass spectrometry

**DOI:** 10.1101/2023.12.24.573280

**Authors:** Junshi Yazaki, Takashi Yamanashi, Shino Nemoto, Atsuo Kobayashi, Yong-Woon Han, Tomoko Hasegawa, Akira Iwase, Masaki Ishikawa, Ryo Konno, Koshi Imami, Yusuke Kawashima, Jun Seita

## Abstract

Mapping protein interaction complexes in their natural state *in vivo* represents the holy grail of protein network analysis. Detection of protein interaction stoichiometry has been an important technical challenge, as few studies have focused this, yet this may be solved by artificial intelligence and proteomics. Here, we describe the development of HaloMS, a high-throughput HaloTag-based affinity purification–mass spectrometry assay for protein interaction discovery. The approach enables the rapid capture of newly expressed proteins, eliminating tedious conventional one-by-one assay. As a proof-of-principle, we used HaloMS to evaluate protein complex interactions of 17 regulatory proteins in human adipocytes. The adipocyte interactome network was validated using an *in vitro* pull-down assay and artificial intelligence-based prediction tools. The application of HaloMS to probe adipocyte differentiation facilitated the identification of previously unknown transcription factor–protein complexes, revealing proteome-wide human adipocyte transcription factor networks, and shedding light on how different pathways are integrated.

## INTRODUCTION

Lifestyle-related diseases, such as diabetes, are estimated to affect hundreds of millions of people worldwide and are a major obstacle to extending healthy life expectancy (https://www.who.int/news-room/fact-sheets/detail/diabetes) (*1*). Many of these diseases are caused by metabolic disorders. Numerous studies have evaluated the molecular regulators of metabolism. For example, studies on transcription factors (TFs) and their interacting partners (i.e., the transcriptional network) have identified various relevant TFs, including CCAAT/enhancer-binding protein and peroxisome proliferator-activated receptor (CEBPs, PPARs) (*2*). To date, most systematic transcriptional network studies have focused on nucleic acids and metabolites (*3, 4*), while few have focused on proteins (*5, 6*). Nevertheless, elucidating the protein networks that are directly related to disease is essential to addressing the central question of which molecular factors regulate metabolism, including those associated with lifestyle-related diseases. Unfortunately, considering the existing one-to-one protein–protein interaction detection technology, at least 400 million experiments would be required to elucidate the protein–protein interactions in an organism with approximately 20,000 genes, such as humans (*7*). Therefore, the development of a nonbiased, high-throughput technology that would allow the comprehensive detection of protein complex interactions is an important technical challenge. Further, with the increasing focus on targeted drug discovery, researchers around the world are aiming to integrate artificial intelligence (AI) in proteomics research (*8–14*).

While the search for biomarkers related to diseases is progressing rapidly, a systematic interaction network of proteins involved in lifestyle-related diseases has not yet been elucidated. Here, we report on the development of HaloMS, a high-affinity capture method that combines the fabrication of *in situ* synthesized proteins for affinity purification with mass spectrometry (APMS). We applied this improved APMS to map protein complex interaction networks in adipocytes, which are closely related to lifestyle-related diseases, using 17 human regulatory proteins as bait. An in-depth analysis of the adipocyte interactome network using the newly developed technology revealed heretofore unknown specific and interacting networks of adipocyte signaling pathways regulated by multiple human proteins.

## EXPERIMENTAL PROCEDURES

### Experimental Design

#### Arabidopsis thaliana

*Arabidopsis thaliana* plants used in this study were in the ecotype Col-0 background. Plants were grown on 0.6% (w/v) gelzan (Sigma-Aldrich, cat. no. G1910) plates containing Murashige and Skoog (MS) salt (Fujifilm Wako, cat. no. 392-00591) and 1% sucrose (Fujifilm Wako, cat. no. 196-00015) medium at 22°C with a photoperiod of 16 h white light (40–50 µmol m^-2^ s^-1^) and 8 h darkness.

### HEK 293-F cells

Commercially available HEK 293-F cells (Thermo Fisher Scientific, cat. no. R79007) were cultured in a 125-mL flask in 30 mL of FreeStyle 293 Expression Medium, according to the manufacturer’s recommendations. After cells were cultured (1 × 10^6^ cell/mL), 1 × 10^10^ cells were harvested 48 h after seeding when cultures were in the exponential growth phase. After discarding the cell medium, cells were detached mechanically by scrapping with 10 mL of cold phosphate-buffered saline (PBS; Nacalai Tesque, cat. no. 05150-45), and the cell solution was washed twice by centrifugations at 1200 × *g* and 4 °C for 3 min. The cell pellet was immediately placed in liquid nitrogen and stored at −80 °C until use.

### HeLa cells

HeLa cells (JCRB Cell Bank, cat. no. JCRB9004) were cultured in a 15-cm dish to 90% confluence in Dulbecco’s Modified Eagle Medium (DMEM, high glucose, Nacalai Tesque, cat. no. 08459-64) containing 10% (v/v) fetal bovine serum (FBS; GIBCO, cat. no. 10437-028) and 1% penicillin/streptomycin (Nacalai Tesque, cat. no. 26253-84) at 37 °C in a 5% CO2 incubator. The cultured cells were detached using trypsin-EDTA (Nacalai Tesque, cat. no. 32777-44) at 37 °C for 5 min, and 1 mL of PBS (Nacalai Tesque, cat. no. 05150-45) was added. Cells (1 × 10^6^ cells/mL) were collected by centrifugation at 1200 × *g* and 4 °C for 3 min; the cell pellet was stored at –80 °C until use.

### Primary human adipose-derived stem cells

Human adipose-derived stem cells (hADSCs) obtained from healthy donors were purchased from Lonza Inc. (Lonza Inc., cat. no. PT5006). Cells at passage two were expanded three times in culture (i.e., to passage 5) in DMEM/F-12 medium (Wako Pure Chemical Industries, Ltd., cat. no. 042-30795) supplemented with 10%(v/v) FBS, 100 U/mL penicillin, and 100 μg/mL streptomycin (basal medium). When the cells reached 80%(v/v) confluence (two days after seeding), adipogenic differentiation was induced with insulin (10 μg/mL, Sigma, cat. no. I9278), dexamethasone (1 μM, Nacalai Tesque, cat. no. 11107-64), 3-isobutyl-1-methylxanthine (50 μM, Nacalai Tesque, cat. no. 19624-44), and rosiglitazone (5 μM, Wako, cat. no. 180-02653; Differentiation medium). hADSCs were cultured for 21 days in basal medium (described as “pre-adipocytes”) or differentiation medium (described as “adipocytes”) with twice-a-week medium change. A total of 1 × 10^8^ cells were harvested by scraping, rinsed three times with ice-cold PBS, and centrifuged at 1200 × *g* and 4 °C for 3 min. The cell pellets were stored at -80 °C until use. All cells were incubated at 37 °C in a 5% CO2 humidified incubator.

### ORF clones

The initial HaloMS testing and comparison with protein array system data were performed using five pIX-HALO expression clones (AT1G32640, AT1G71930, AT3G62420, AT5G28770, and AT5G65210), as described previously (*15–17*). To test human cultured cells, human pENTR-ORFs clones (Dnaform, National Institute of Technology and Evaluation) in the Gateway-compatible entry vector were recombined using Gateway LR clonase II (Thermo Fisher Scientific) into the pIX-Halo:ccdb destination vector (*16, 18*).

### Preparation of plant proteins

Protein isolation from mature leaves of *A. thaliana* wild type Columbia was performed as previously described (*19*), with slight modification. Briefly, 15-d-old mature leaves were frozen in liquid nitrogen, ground using MB1200 Multi-beads shocker (Yasui Kikai), and homogenized in an extraction buffer [50 mM Tris-HCl pH 7.5, 150 mM NaCl, 10% glycerol, 2 mM EDTA, 5 mM dithiothreitol, 2% (v/v) IGEPAL CA-630, 2 mM sodium molybdate, 2.5 mM NaF, 1 mM phenylmethylsulfonyl fluoride, 1 mM sodium orthovanadate, and 1 Roche cOmplete ULTRA tablet/50 mL solution]. After incubation at 4 °C for 30 min with gentle rolling, samples were sonicated using a Bioruptor (UCD-250, Cosmo Bio) with High mode for 10 s sonication plus 10 s without sonication (interval) as one set. After 10 sets of sonication, the crude protein solution was incubated at 4 °C for an additional 30 min. Phenylmethylsulfonyl fluoride was added for a final concentration of 2.44 mM. The samples were then centrifuged at 15,310 × *g* at 4 °C for 10 min. The supernatant was transferred to a new tube, and the centrifugation step was repeated twice more. The protein in the final sample was quantified using the Qubit Protein Assay Kit with the Qubit 2.0 Fluorometer (Thermo Fisher Scientific).

### Protein extraction from human cells

Proteins from human cell lines were extracted in IP lysis buffer (Thermo Fisher Scientific, Pierce IP Lysis Buffer, cat. no. 87788) containing protease inhibitors (cOmplete ULTRA tablet, Sigma-Aldrich, cat. no. 5892791001) and phosphatase inhibitors (PhosSTOP tablet, Sigma-Aldrich, cat. no. 4906837001). The buffer was added to cell pellets obtained from 1 × 10^10^ cells, and the samples were agitated at 4 °C for 30 min. The cell lysate was then centrifuged at 18,000 × *g* at 4 °C for 30 min, and the supernatant was collected. Protein concentration in the extract was determined using a BCA protein assay kit (Thermo Fisher Scientific, cat. no. 23225) and adjusted to 2 μg/μL using the IP lysis buffer.

### First-generation HaloTag ligand plate assay

The initial HaloTag-based APMS (HaloMS) was performed using a commercially available HaloTag ligand plate (Promega, HaloLink 96 Well Plate, cat. no. CS180802). For HaloMS plate preparation, 500 ng of HaloTag fusion ORF plasmid were translated, and the proteins expressed in a HaloLigand plate at 30 °C for 2 h, using the TNT wheat germ system (Promega, cat. no. L4140). The reaction mixture contained 9.5 μL of nuclease-free water, 12.5 μL (50%) of wheat germ extract, 1.0 μL of 10× TNT buffer, 0.5 μL of amino acid mix without methionine, 0.5 μL of amino acid mix without lysine and cysteine, 0.5 μL (50 unit) of T7 RNA polymerase, and 0.5 μL of RNase inhibitor (Promega). The wells were then washed with PBS with 0.1% (v/v) Tween 20 (PBST) and subsequently incubated with a human cell lysate containing 50 mg proteins per 25 μL proteins at 4 °C for 2 h to allow for protein interaction. The wells were washed in PBST and MS-grade water and used for on-plate protein digestion.

### On-plate protein digestion

One-hundred microliters of 50 mM Tris-HCl (pH 8.0) and 500 ng of trypsin/Lys-C mix (Promega, cat. no. V5072) were added to the washed HaloTag ligand plate following the first-generation assay and mixed gently at 37 °C overnight to digest the proteins. The digested sample (supernatant) was collected into a new 96-well PCR plate by multi-channel pipetting, treated with 20 mM tris (2-carboxyethyl) phosphine at 80 °C for 10 min, and alkylated using 30 mM iodoacetamide at 25 °C for 30 min in the dark. Subsequently, the alkylated sample was acidified with 20 μL of 5% (v/v) trifluoroacetic acid and desalted using a STAGE tip (GL Sciences Inc., cat. no. 7820-11200) according to the manufacturer’s protocol. This was followed by drying in a centrifugal evaporator (miVac Duo concentrator, Genevac Ltd.). The dried sample was resuspended in 10 μL of 2% (v/v) acetonitrile containing 0.1% trifluoroacetic acid. Finally, 4 μL of samples were used for LC-MS/MS analysis.

### LC-MS/MS

Peptides were directly injected onto a 75 μm × 12 cm nanoLC nano-capillary column (Nikkyo Technos Co., Ltd., Tokyo, Japan) at 40 °C and separated over a 60 min gradient at a flow rate of 200 nL/min using an UltiMate 3000 RSLCnano LC system (Thermo Fisher Scientific). Peptides eluted from the column were analyzed using a Q Exactive HF-X or Orbitrap Exploris 480 (Thermo Fisher Scientific) for overlapping window DIA-MS (*20, 21*). MS1 spectra were collected in the range of 495–745 m/*z* at 30,000 resolution to set an automatic gain control target of 3e6 and maximum injection time of 55 ms. MS2 spectra were collected in the range of > 200 m/*z* at 30,000 resolutions to set an automatic gain control target of 3e6 with an “auto” maximum injection time, and stepped normalized collision energy of 22, 26, and 30%. The isolation width for MS2 was set to 4 m/*z* and overlapping window patterns in 500–740 m/*z* were used as previously reported (*22*). MS files were searched against a spectral library of human proteins and HaloTag using Scaffold DIA (Proteome Software, Inc.). The spectral library of human proteins and HaloTag was generated from the amino acid sequence of the human protein database (UniProt, Proteoem ID UP000005640, reviewed, canonical) by Prosit (*23, 24*). The Scaffold DIA search parameters were as follows: experimental data search enzyme, trypsin; maximum missed cleavage sites, 1; precursor mass tolerance, 10 ppm; fragment mass tolerance, 10 ppm; static modification, cysteine carbamidomethylation. The protein identification threshold was set at both peptide and protein false-discovery (FDR) rates < 1%. Peptide quantification was performed using the EncyclopeDIA algorithm in Scaffold DIA (*25*). For each peptide, the four highest-quality fragment ions were selected for quantitation. MS data generated in the current study were deposited in JPOST under the accession number PXD041085 (https://repository.jpostdb.org/).

### Pull-down assay

The assay was performed using HaloLink magnetic beads, according to the manufacturer’s recommendations (Promega). The corresponding human ORF clones were transferred by Gateway LR recombination into pIX-Halo:ccdB and pIX-3×HA:ccdB destination vectors (*16*). Competent bacteria (*Escherichia coli*, strain TOP10, Thermo Fisher Scientific, cat. no. C404010) were transformed with the resulting recombination products. The transformants were selected in liquid Plusgrow II medium (Nacalai Tesque, cat. no. 08202-75) containing 50 mg/mL ampicillin, and plasmid DNA was extracted and purified using the NucleoSpinPlasmid kit (Macherey-Nagel). Bait proteins encoded by pIX-Halo were expressed using the TNT T7-coupled wheat germ extract system (Promega, cat. no. L4140) according to the manufacturer’s recommendations. Twenty microliters of bait proteins (pIX-Halo-ORFs) were mixed and agitated with rotation using 5 μL of Halo magnetic beads in a total volume of 50 μL of PBST at 25 °C for 1 h. Subsequently, beads with the HaloTag fusion protein baits were washed with PBST, mixed with 20 μL of prey protein fused with a triple hemagglutinin (3×HA), and agitated with rotation at 25 °C for 2 h. Next, the mixture was washed three times with 200 μL of PBST. The washed beads were heated in 20 μL of SDS sampling buffer (Thermo Fisher Scientific, cat. no. NP0007) at 90 °C for 5 min. The bait and prey proteins (2.5 μL; 10% input) were used as a loading control for the original protein amount.

### Second-generation HaloTag ligand plate assay

HaloTag-PEG-biotin ligand (Promega, cat. no. G859A; 500 pmol in 100 μL of PBS buffer) was added to the wells of avidin-coated plates (Thermo Fisher Scientific, cat. no. 15507), and incubated with moderate agitation for 1 h at 25 °C. The plates were then washed three times with nanopure water and incubated with 200 μL of PBSB for 1 h at 25 °C with moderate agitation. Finally, the plates were placed in a clean environment (Labconco, purifier class II biosafety cabinet) for 60 min to dry, and the coated plate was stored at –80 °C until use.

### Protein–protein interaction AI analysis

AlphaFold (v2.1.1) was downloaded from https://github.com/deepmind/alphafold onto Ubuntu 20.04 comprising 28 cores, 128 GB memory, two NVIDIA RTX A5000, and three 4TB SSD (*14*). The list of human and *Arabidopsis* protein sequences was obtained from the UniProt database (*26*) and The Arabidopsis Information Resources (on July 11, 2022 at https://www.uniprot.org/, https://www.arabidopsis.org/), respectively. AFM was performed as the default implementation condition. AFM provides five model confidence scores, the interface-predicted template modeling scores (ipTM). A weighted combination of pTM and ipTM was used to compute the interaction confidence, with reference to previous reports, where model confidence = 0.8 × ipTM + 0.2 × pTM (*14*). The mean and SD values were calculated based on these five scores. *P*-values were obtained by comparing pairs of test sets. HaloTag pairs that differed at *P* < 0.05 were considered positive. LocalColabFold (v1.4.0) was downloaded from https://github.com/YoshitakaMo/localcolabfold into Ubuntu 20.04 comprising eight cores, 64 GB memory, NVIDIA GeForce RTX 3090, 8TB SSD, and 12TB HDD. ColabFold (v1.3.0) was downloaded by running the local ColabFold script (*11*). In addition, AlphaFold2-multimer-v2, which is a trained model parameter of AFM, was downloaded from https://github.com/deepmind/alphafold. LocalColabFold were performed using the following settings:--amber, --use-gpu-relax, --templates, --num-recycle 3, --random-seed 0, --num-models 1, --model-order 1, --model-type AlphaFold2-multimer-v2.

### UMAP and hierarchical clustering

Protein homology determined in CFM was visualized based on the metagenome visualization method (*27*). First, each sequence was embedded as a 256-dimensional vector using localcolabfold (*11*) and visualized using the umap v0.5.3 and scanpy v1.9.3 Python packages (*27, 28*), where dimensionality reduction was applied directly to the embedding vectors (use_rep=‘X’ in scanpy.tl.umap) with default parameters (3 or 4-nearest-neighbor graph via approximate Euclidean distance, UMAP min_dist=0.5) (*29*). Hierarchical clustering and dendrogram generation were performed using embedding vectors obtained from ColabFold using the hierarchical clustering algorithm (“single” method, “euclidean” metric) from the scipy v1.9.3 package in Python (*30, 31*).

### Statistical Rationale

We conducted *t*-tests for most statistical analyses if no specification was indicated. Statistical analysis of data was performed using BellCurve for Excel (Social Survey Research Information Co., Ltd.) and https://www.socscistatistics.com/tests/. The employed quantification methods and web-based tools are described in each relevant figure/Supplementary Table legend.

## RESULTS

### Development of HaloMS, a HaloTag-based affinity purification–mass spectrometry (MS) system

In conventional APMS, tagged protein-encoding sequences are first transformed into a cell and then retrieved by binding to ligand-capturing beads in a reaction tube. The interacting protein complex is then pulled down from a biological sample, and the proteins are identified one-by-one via MS. We modified this classical system by incorporating a high-affinity capture tag, the HaloTag (Promega), to immobilize nascent proteins expressed with the cell-free system in a 96-well plate (Figure 1) (*15*). In the assay, *in vitro* expressed and translated HaloTag-labeled proteins are covalently bound to a small chemical ligand chloroalkane that coats the plate surface. This approach allows for rapid capture of newly expressed fusion proteins and eliminates the cumbersome process of purifying individual proteins on ligand beads, avoiding the possibility of compromising protein functional integrity.

**Figure 1.**
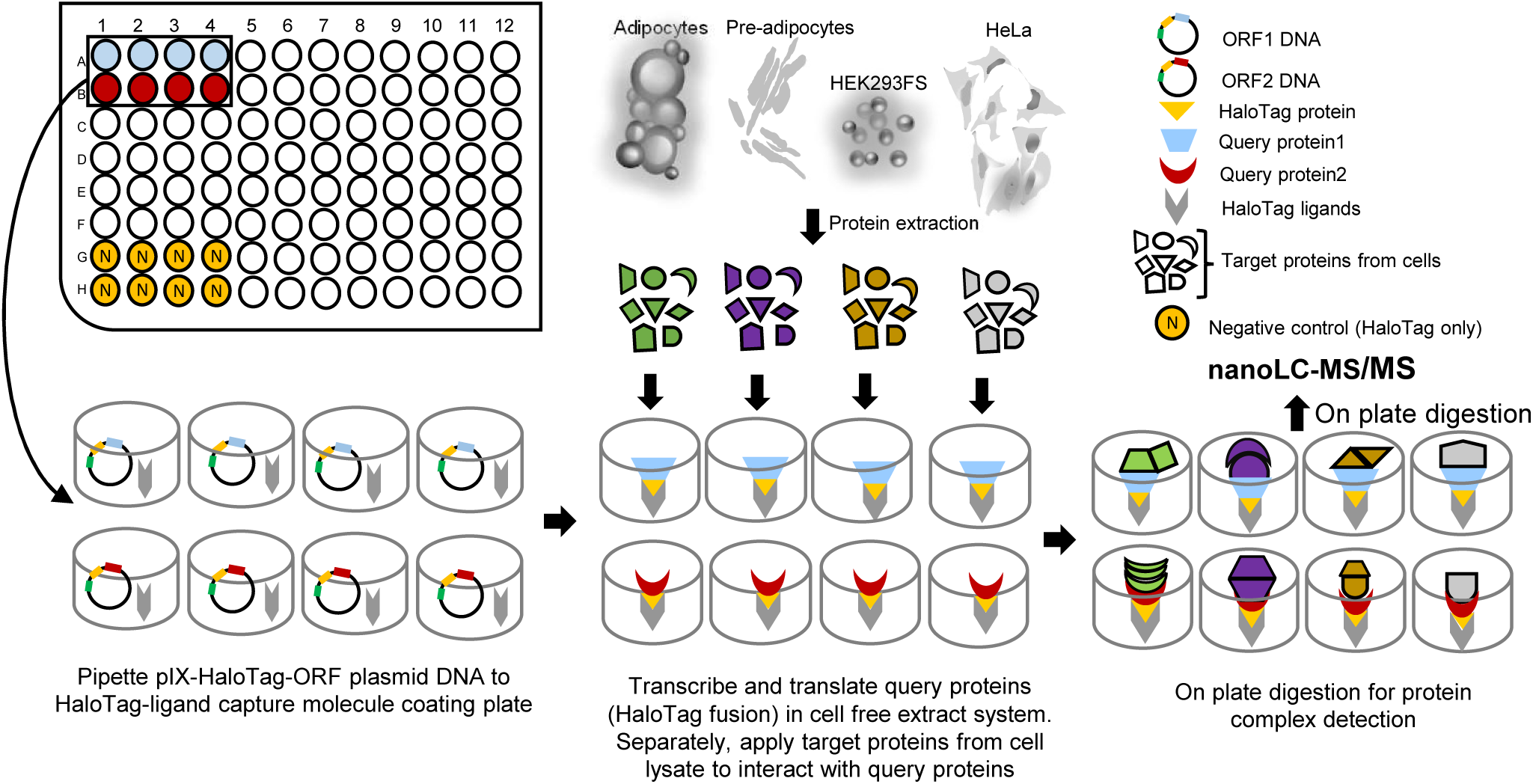
Overview of the HaloMS assay. HaloTag-ORF plasmid DNA is placed in a well of a HaloTag-ligand-coated plate. The addition of coupled transcription– translation reagent results in protein expression and probe protein capture, accordingly. Target protein lysates from various cells can be added to the wells, and protein complex interactions are detected by liquid chromatography-tandem mass spectrometry (LC-MS/MS).

To test the activity of the HaloTag-fused protein bound to the chloroalkane ligand-coated 96-well plate, we detected protein–protein interactions in the model organism *A. thaliana* using a query protein set previously used in a similar protein immobilization method with HaloTag (*15*). After HaloTag-based affinity purification MS (hereafter, HaloMS), we assessed the expression of five query TFs, namely, AT1G32640, AT1G71930, AT3G62420, AT5G28770, and AT5G65210, based on the MS intensity. The MS intensities of the query proteins were between 8.41 and 9.52 (log 10 scale), indicating that the levels of protein produced from the five ORFs were consistent (Supplementary Table S1). We then probed the lysate of *Arabidopsis* leaves containing 9999 identified proteins, including splice variants, for complex interactions with the five TFs, which were translated and expressed using the wheat germ expression system and bound to HaloTag ligand plates (Supplementary Table S2). We set the protein identification threshold for the peptide and protein at a FDR of < 1%. An interaction was scored as positive when the log ratio of the signal intensity obtained from the negative control (i.e., 33 kDa HaloTag protein) was > 1 standard deviation (SD) above the median. By using this criterion, HaloMS for the five query proteins produced a dataset of 740 interactions and 306 proteins (Supplementary Table S3). We then calculated the overlap between the interactions revealed by HaloMS and those in a published protein array dataset containing 1544 interactions and 1234 proteins, which uses the same HaloTag protein immobilization method (*15*). HaloMS recapitulated a statistically significant 10 out of 709 protein array interactions in the shared protein space (*P* < 0.00001, χ^2^ test with Yates’ correction; hereafter “χ^2^”). This significant association between the two approaches suggested that these interactions are not affected by the test method (HaloMS vs. protein array).

We next investigated the degree to which the identified protein–protein interactions could be classified using gene ontology (GO) functional annotation terms to assign a higher probability of biological relevance. The resultant enriched terms for all target proteins interacting with the five query proteins, as identified by HaloMS, are provided in Supplementary Table S4. Except for “mRNA binding,” “RNA binding,” and “binding,” a significant proportion (*P*-value > 1.64E-14) of HaloMS targets of the five TFs tested were related to the enzyme and carbon binding activity.

By contrast, according to the enrichment analysis based on the protein array data (*15*), most targets were related to transcription, “binding,” and “protein binding” (Supplementary Table S5). The considerable differences in GO enrichment of the protein–protein interactions detected using the two methods could be explained by a wider dynamic range of the expressed proteins in the cell lysate used in HaloMS compared to that in the input used for protein arrays (Supplementary Figure S1 A and B).

We next determined the overlap of interactions between the HaloMS dataset and the literature-curated interactions in the BioGrid database (https://thebiogrid.org/). HaloMS recapitulated a statistically significant 1 of 79 BioGrid interactions in the shared protein space (*P* = 0.019044, χ^2^ test). This moderate overlap may be associated with differences in the biochemical experiments, fusion tags, etc., leading to the detection of different subsets of true interactions in the specific assay (*32*). Alternatively, this could be due to the relatively small number of *Arabidopsis* interactions available in the dataset (81,804 in BioGrid on August 27, 2021).

Collectively, the low overlap suggested that HaloMS is highly complementary to the public dataset for *Arabidopsis*, which was generated primarily using a yeast two-hybrid system (Y2H). Further, HaloMS has several advantages. For example, it does not require the preparation of a large number of clones and allows for *in vivo* screening of protein complexes in a variety of physiological states. It also avoids the cumbersome stages of individual transgenic cell line preparation and carrier bead purification of conventional APMS, as the query proteins are synthesized cell-free and captured on plates on demand.

### HaloMS reliably detects protein complex interactions in cultured human cells

We then tested HaloMS for the detection of protein–protein interactions in human cells. To this end, we transferred a set of 16 Gateway pENTR ORF clones, including ATF3, GATA2, and FOS, which encode well-studied human protein-coding genes into a Gateway Convertible HaloTag expression vector, and *in vitro* expressed and captured the proteins on a HaloLink 96-well plate (see Materials and Methods). The intensity of the expressed 16 query proteins was 7.45–9.67 (log 10 scale), indicating consistent levels of produced and captured proteins (Supplementary Figure S1C). We then used these 16 proteins to probe the lysate of a mixture of HeLa and HEK293F cells containing 7130 proteins to detect their protein complex interactions (Supplementary Table S6). The same criterion for FDRs of peptide, protein, and interaction scoring as positive was applied as for the initial *Arabidopsis* experiment. Thus, the generated HaloMS dataset contained 3302 interactions and 913 proteins (Supplementary Table S7). We determined its interaction overlap with the 1,057,434 interactions documented in BioGrid (on June 29, 2022, at https://thebiogrid.org/). HaloMS recapitulated a statistically significant number of 183 of 4220 literature-curated interactions in the shared protein space (*P* < 0.00001, χ^2^ test). This indicated that the *in vitro* expressed proteins immobilized on the 96-well plate were capable of specific interactions with the thousands of proteins in the lysate. Thus, the interaction overlap analysis indicated that HaloMS yields reliable protein–protein interaction data that are statistically indistinguishable from the literature-curate interaction dataset.

### *In vitro* pull-down assay validates the HaloMS dataset

A critical concern for any new technology regards the quality of the obtained data. To address this, we followed an approach that we have previously described, in which a subset of interactions from a new dataset was systematically validated in a second interaction assay, in our case, a pull-down assay (*32–35*). As a caveat, no assay can detect all protein–protein interactions, and each has a different interaction detection profile (*34, 36*). To validate the HaloMS dataset generated for the human cell lines, we selected 32 out of 3302 pairs of the detected interactions, corresponding to approximately 1% of all interactions in the dataset, and subsequently tested them using a pull-down assay (Figure 2A, Supplementary Figure S2, Supplementary Table S8) (*15*). We compared the obtained data with benchmarking data generated previously (*15, 37*). In brief, we benchmarked the new data against (i) a set of known positive interactions (PRS: 49 pairs) and (ii) a set of randomized pairs of interactions that, to date, have not been supported by experimental evidence (RRS: 69 pairs). Of the 32 pairs, 7 (21.8%) scored positive (Figure 2B, Supplementary Table S8). This proportion was statistically different from that for PRS and RRS (*P* = 0.0003 and 0.0041, respectively, Fisher’s exact test; Figure 2B).

**Figure 2.**
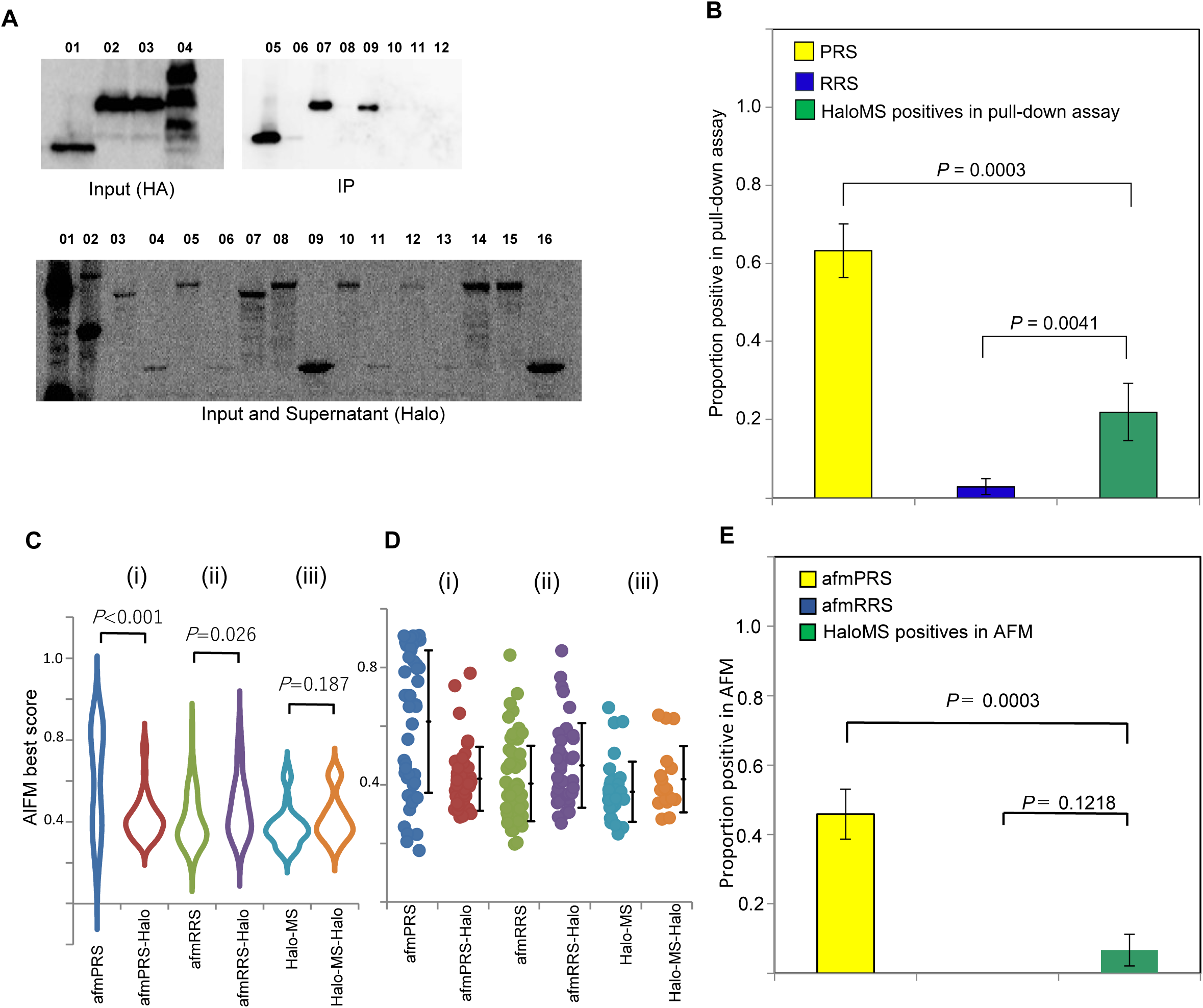
Quality of the HaloMS assay. (A) HaloMS protein–protein interactions are recaptured by pull-down assay. Representative results of the *in vitro* pull-down experiments for Halo alone, Halo-tagged, and 3×HA-tagged proteins are shown. Five percent of input is detected using an anti-HA antibody (labeled “Input (HA),” lane 01-04); co-purified Halo-tagged and 3×HA-tagged proteins are detected using an anti-HA antibody (labeled as “IP,” lane 05–12), and 5% of supernatant after binding to HaloLink magnetic beads and input for HaloTag-labeled proteins (labeled “Input and Supernatant (Halo),” lane 01–16) are tested to determine the relative amounts of Halo-tagged and HA-tagged proteins, and the binding efficiency to HaloLink magnetic beads in the experiments. The results for all protein pairs evaluated in the pull-down assay are shown in Supplementary Figure S3 and Supplementary Table S8. (B) The proportion of positive scoring pairs by pull-down assay. The histogram shows the proportion of positive scoring pairs from the PRS dataset, the set of randomized sample pairs from reference 15, and the set of sample pairs positive in HaloMS. Error bars represent the SE of the proportion. The proportion of positive PRS pairs (yellow bar) is significantly higher than that of RRS pairs (blue bar; *P* < 0.0001, Fisher’s exact test). (C) Artificial intelligence-based prediction of protein– protein interactions is detected by HaloMS and pull-down assay. The violin plot distribution of the highest scores is obtained by AlphaFold-Multimer (AFM) across six datasets (Supplementary Table S9). (D) Dot plot distribution showing the average, first-quartile, and third-quartile AFM best scores for six datasets: afmPRS, afmPRS-Halo, afmRRS, afmRRS-Halo, HaloMS, and HaloMS-Halo. (E) Artificial intelligence-based prediction of protein–protein interactions. The proportion of positive scoring pairs from the afmPRS dataset, the set of randomized sample pairs from reference 15, and the set of sample pairs positive in HaloMS are shown. Error bars represent the SE of the proportion.

The validation rate for the HaloMS positive interactions suggested that the obtained dataset reflects a different aspect of the well-documented interactions reported in the literature and detected by direct protein–protein interaction analysis. In contrast, HaloMS yielded protein interaction data of a reliable quality that were statistically different from the random reference set. As anticipated, the reproducibility (21.8%) of HaloMS, which includes indirect interactions in a protein complex, vs. PRS, which is composed of binary protein–protein interactions, was low. Taken together, the analysis of the pull-down assay data against the benchmark dataset indicated that HaloMS can be used to reveal interactions that are statistically independent of binary interaction data, literature-curated interactions, and random protein pairs.

### AI analysis supports protein–protein interactions detected by HaloMS

Although we were able to validate the novel HaloMS protein–protein interactions using a pull-down assay, we next asked whether these interactions could be supported by another type of analysis, e.g., AI predictions based on the amino acid sequence. Accordingly, we used AlphaFold-Multimer (hereafter AFM) to predict and validate protein–protein interactions detected by HaloMS and the pull-down assay. To determine the FDR of the human cultured cell HaloMS dataset, we measured the AFM sensitivity and background by benchmarking against a set of controls supported by the pull-down assay, PRS, and RRS. This allowed us a subsequent interpretation of the retest rate of the new dataset by HaloMS, which was also validated by the pull-down assay, in light of these benchmarking data. To measure the sensitivity and background of the AFM assay, we benchmarked it against (i) a set of 48 well-documented interactions from the literature (afmPRS) (*15*) and (ii) a set of 71 random interactions from the published dataset (afmRRS) (*15*). As a new dataset, we tested 32 pairs from the HaloMS dataset from human cultured cells, which were supported by the pull-down assay (Supplementary Table S8). For each dataset, a negative control set with HaloTag-only proteins was also analyzed by AFM to obtain a validity score (Supplementary Table S9). The distribution of AFM best scores from the three datasets validates the characteristics of each dataset (Figure 2C and D). In the benchmark experiments for the afmPRS (i) and afmRRS (ii) sets, the reproducibility of afmPRS differed significantly (t-test, *P* < 0.001) from that of the negative-control pairs; those of afmPRS-Halo and afmRRS were not significantly different, as expected for a negative control; afmRRS-Halo scored higher than afmRRS (*P* = 0.026). However, the distribution of recaptured HaloMS interactions was statistically indistinguishable from HaloMS-Halo (Figure 2C, iii) by AFM (*P* = 0.187) despite 21.8% reproducibility in the pull-down assay.

We used an average of the five scores to determine the model confidence obtained by AFM to compute the significance of differences between each protein pair and the HaloTag-only control pairs, as in the pull-down assay (for a “digital” pull-down assay; Supplementary Table S10). In benchmark experiments using (i) and (ii) reference sets, 22 of 48 afmPRS pairs (46%) and 0 of 55 afmRRS pairs (0%) scored significantly higher than the HaloTag-only controls, defining the sensitivity and background of the digital pull-down assay. Of the 30 pairs of HaloMS, the scores for 2 (6%) were significantly positive in the digital pull-down assay, statistically different from the afmPRS result (*P* = 0.0003, Fisher’s exact test; Figure 2E). This reproduction rate was similar to the obtained pull-down result, including all positive pairs (6/30 pairs, 20%, Supplementary Table S10), and was not significantly different from the afmRRS set (*P* = 0.1218, Fisher’s exact test; Figure 2E). The afmPRS, afmRRS, and HaloMS scores by AFM ranged from 0.500 to 0.652, 0.3061 to 0.3473, and 0.3465 to 0.42369, respectively, estimated using a 95% confidence interval. These results support the findings of the pull-down evaluation, i.e., the HaloMS data represent complex interactions, not just binary interactions, with few directly interacting pairs. Although the results by AFM were similar to those of the pull-down assay, the number of 30 pairs with pull-down information obtained by HaloMS may not have been as sufficient as the number of pairs evaluated by AFM.

### HaloMS TF interactome network for adipocyte differentiation response

As a proof-of-principle, we used HaloMS to generate an adipocyte interactome dataset. An adipocyte cell stores triglyceride droplets and helps to convert lipids into energy. In addition, an adipocyte can secrete hormones and other effector chemicals that play important roles in metabolism control, including cell turnover (*38*), adaptation (*39–41*), and endocrine roles (*42, 43*). These adipocytes, which are associated with metabolic diseases through different pathways, can be used for mapping protein interaction networks by applying HaloMS technology; it may be possible to construct a centralized signaling pathway. We set 17 human proteins as query proteins, including nine TFs (ATF1, ATF2, ATF3, ATF5, GATA2, CEBPA, CEBPD, CREB1, and PPARA) with known roles in adipocytes that mediate differentiation from pre-adipocytes. We additionally developed an alternative capturing technique to produce query proteins on the 96-well plate. In this alternative protocol, HaloTag ligand concentrations were lower than those used in the first-generation HaloMS to lower the cost of the HaloTag capturing plate; this also addressed the problem of the original HaloTag plate ceasing to be commercially available (see Materials and Methods for details). We assessed the protein capturing efficiency of four query proteins (GATA2, ATF3, CTNNA1, and HaloTag) based on MS intensity. The intensities of the query proteins were 7.78–9.79 (log10 scale), indicating that consistent protein levels were being captured (Supplementary Figure S3 A, red), with the capture levels similar to those of first-generation HaloMS (8.14–9.62, log10 scale MS intensity, Supplementary Figure S3 A, light blue). Considering that most TFs function as part of a complex (29, *44–46*), we expected that the identification of additional proteins that interact with TFs in adipocytes would reveal novel adipocyte differentiation signaling components and connections among the already-known signaling pathways. Further, for several of the proteins in a set of 17 human proteins, including nine TFs with known roles in the adipocyte, some interactions were already identified in the BioGrid database (thebiogrid.org). To systematically address the adipocyte-specific interaction network, we probed four different cell types (adipocyte, pre-adipocyte, HeLa, and Human Embryonic Kidney cell as HEK) with the 17 query proteins using HaloMS. Using these cell types, the technique was applicable to heavily used cancerous, multi-generation, passaging cultures, such as HEK and HeLa, as well as primary cells that presumably reflect physiological conditions, such as adipocytes. To confirm protein expression levels in the input sample (the four types of cells), we assessed global protein expression across the cell types using liquid chromatography-tandem MS (LC-MS/MS) after proteins were digested, alkylated, and acidified. Among the 9405 proteins with similar expression ranges, we identified 7258 in adipocytes, 7552 in pre-adipocytes, 8149 in HeLa cells, and 8225 in HEK cells (Supplementary Figure S3B, C, Supplementary Table S11). The expression level of over 65% of proteins was similar in the different cells (Supplementary Figure S3D). Six percent of proteins fell outside the commonly expressed protein set. The intensities of query proteins immobilized on the 96-well plate and used for the screen were 7.64–9.34 (log10 scale), indicating that consistent protein levels were produced from the 17 ORFs in the assay (Supplementary Figure S3E, Supplementary Table S12). We scored a positive interaction when the average intensity from multiple datasets was > 1 SD above the median of the ratio, a 3.07-fold difference on average, obtained with only the 33-kDa HaloTag proteins as the negative control. Pairs with negative control of 0 and no ratio available were considered positive at 1 SD from the median of the intensity distribution. According to these criteria, the HaloMS screen of the 17 human proteins in the four different cell types produced a TF-HaloMS dataset containing 6512 interactions and 1690 proteins (Figure 3A, Supplementary Table S13). Over 70% of the interactions were cell type-specific, unlike the expression of input proteins. Only 96 out of 6512 interactions (1.5%) were commonly detected in all examined cell types. Comparison of the interaction number and protein expression levels revealed that for the adipocyte-specific interactions (1201 pairs), low-expression proteins were involved in the highest number of interactions (488 pairs, 40%; Figure 3B). Other cell-specific protein–protein interactions tended to be detected in proteins with relatively high expression levels, e.g., in HeLa and HEK cells (Figure 3B). We then determined the overlap of interactions between the TF-HaloMS dataset and literature-curated interactions in BioGrid (thebiogrid.org). The HaloMS screen of the four different cell types recapitulated statistically significant 443 of 5974 BioGrid interactions in the shared protein space (*P* < 0.00001, χ^2^ test; Supplementary Table S13). The significant overlap suggested that HaloMS using first- and second-generation 96-well plates was an approach that is statistically reliable not only to interactions described in the literature but also to novel protein–protein interactions.

**Figure 3.**
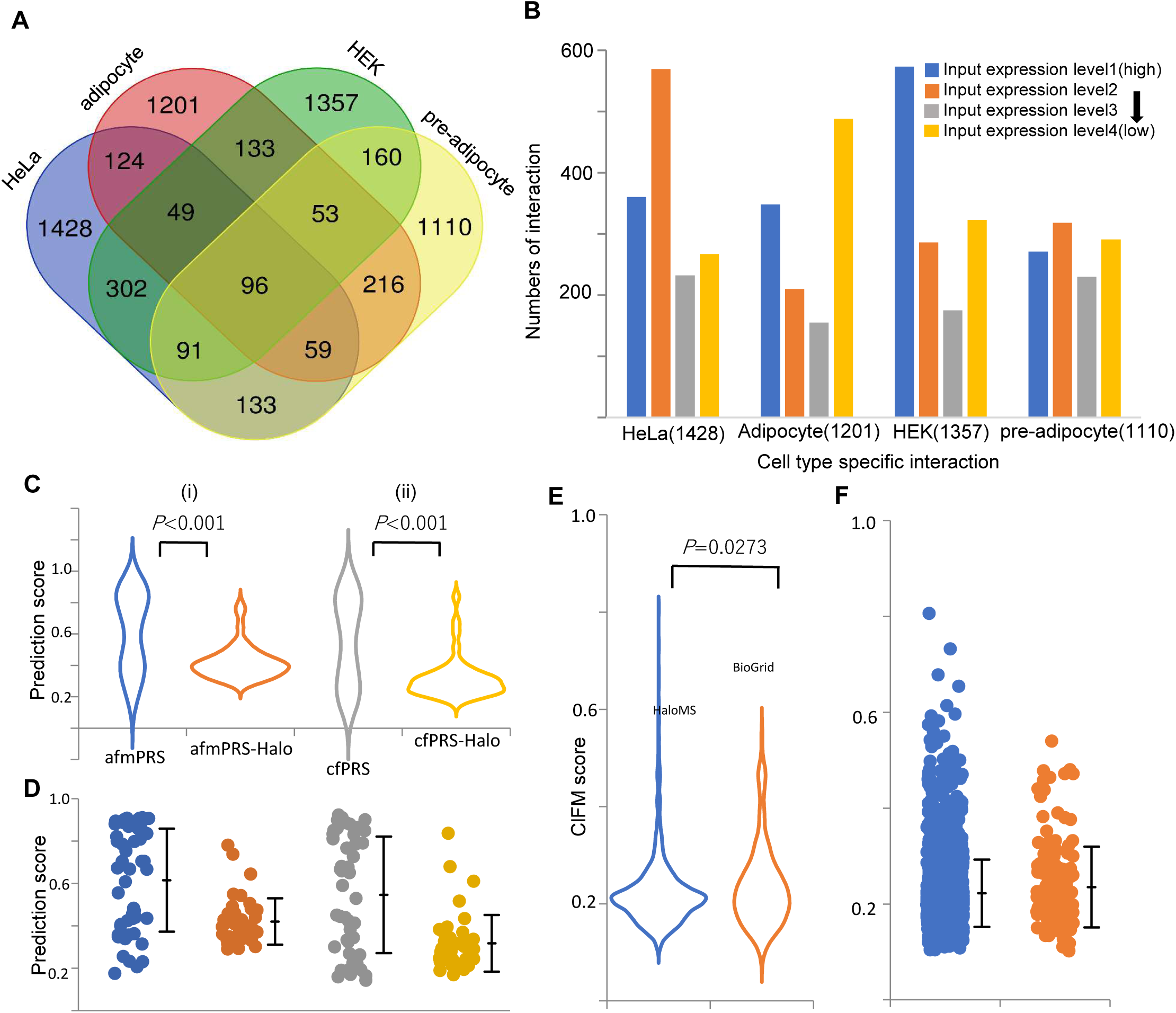
Interactome profiling in four different cell types. (A) Venn diagram of the interactome status of four cell types. Over 70% of the interactions are cell-type specific, unlike the input protein expression. Venn diagram is generated using software available at https://bioinformatics.psb.ugent.be/webtools/Venn/. See also Supplementary Table-S13. (B) Comparison of the number of interactions and protein expression levels. The highest number of interactions for low-expression proteins is detected among adipocyte-specific interactions. (C) Violin plot distribution of AFM and CF scores in the PRS dataset showing the validity of the interaction assessment (Supplementary Table S14). (D) Dot plot distribution of the average, first-quartile, and third-quartile AFM and CF scores for the datasets. (E) Violin plot distribution of CF scores for the HaloMS dataset validating the interaction assessment of each dataset of adipocyte- and preadipocyte-specific interactions. (F) Dot plot distribution of the average, first-quartile, and third-quartile CF scores for each dataset. In the HaloMS experiments for (iii) novel (blue) and known (orange) interactions of the adipocyte–preadipocyte-specific interaction dataset, the distribution of the CF score overlaps in the novel vs. known-pair comparison. See also Supplementary Table S14.

### Biological validity of the TF-HaloMS dataset

We next obtained evidence of the biological validity of the TF-HaloMS dataset and new insights into signal transduction during adipocyte differentiation. To this end, we validated several of the biologically interesting novel interactions by AI. Specifically, we predicted protein–protein interactions using ColabFold Multimer (hereafter CFM), which offers an accelerated prediction of protein structures and complexes with a 40−60-fold faster search than AFM (*11*). To estimate the FDR of the HaloMS adipocyte dataset and pre-adipocyte-specific interactions, we determined the sensitivity and background of CFM using the PRS dataset and compared the findings to the AFM results (Figure 3C and D). Due to the limited machine resources, we used only one CFM score to analyze significant differences for 49 well-documented interactions from the literature (cfPRS) (*15*), HaloTag-only negative controls (cfPRS-Halo), and the adipocyte- and pre-adipocyte-specific interaction dataset from HaloMS (Figure 3E and F, Supplementary Table S14). The distribution of the dataset CFM scores validated the traits of each dataset. In the benchmark experiments for the datasets for: (i) afmPRS and the negative control (afmPRS-Halo), and (ii) cfPRS and the negative control (cfPRS-Halo), the reproducibility of cfPRS, as well as the afmPRS, was significantly different (*t*-test, *P* < 0.001) than that of negative-control pairs (*t*-test, *P* < 0.001; Figure 3C). Of the 1201 and 1110 interactions selected based on all adipocyte- or all pre-adipocyte-specific interactions in the TF-HaloMS dataset, 1093 and 1028, respectively, were validated using CFM (Supplementary Table S14). The TF-HaloMS dataset evaluated by CFM included 86 known pairs of adipocyte-specific pairs and 71 known pairs of pre-adipocyte-specific pairs, and the distribution of the CFM scores for these and novel interactions were not significantly different (*P* = 0.027; Figure 3E, Supplementary Table S14). This overlap between known and novel interactions suggested that the TF-HaloMS dataset of adipocyte- and pre-adipocyte-specific interactions is a reliable protein–protein interaction dataset.

## DISCUSSION

### HaloMS-based exploration of network communities in adipocyte

During protein network analysis, it is important to identify densely interacting components that function together, i.e., the intra-network community (*47*). Accordingly, we analyzed the pathways activated in the four different cell types. More specifically, we used average-linkage clustering to identify intra-network communities in adipocytes and investigated their biological relevance.

Pathway analysis for the cell type-specific pairs of proteins (Supplementary Figure S4, Supplementary Table S13) identified 83 communities containing more than four proteins in the HaloMS dataset as significantly cell type-associated proteins (Supplementary Figure S4, Supplementary Tables S15 and S16). The adipocyte-specific interactions included mitochondria-associated neurodegenerative disease pathways (including Alzheimer’s, Parkinson’s, and Huntington’s diseases; Supplementary Figure S4, S5A, Supplementary Table S17) (*48*). The activation of these pathways implied that the thermogenesis subnetwork in adipocytes was activated, indicating the biological relevance of the TF-HaloMS dataset. The specific enrichment of the PPAR signaling pathway (*49*), non-alcoholic fatty liver disease (*48*), and fluid shear stress and atherosclerosis (*50, 51*) in adipocytes further supported the validity of the interactions detected in the current study (Supplementary Figure S5B-D, Supplementary Tables S18–S20). The pre-adipocyte-specific interactions, including a pathway for adrenergic signaling in cardiomyocytes (Supplementary Figure S6A, Supplementary Table S21) (*52*), thyroid hormone synthesis (Supplementary Figure S6B, Supplementary Table S22) (*53, 54*), AMPK signaling (Supplementary Figure S6C, Supplementary Table S23) (*55*), platelet activation (Supplementary Figure S6D, Supplementary Table S24) (*56*), peroxisome (Supplementary Figure S6E, Supplementary Table S25) (*57, 58*), and N-glycan biosynthesis (Supplementary Figure S6F, Supplementary Table S26) (*59*), supported the correct profiling of interactions by HaloMS.

The relationship between the interacting proteins is a critical question in HaloMS, which was devised to specifically detect protein complexes. Accordingly, we clustered groups of proteins interacting with the 17 protein baits obtained specifically from adipocytes by HaloMS using uniform manifold approximation and projection (UMAP), dendrograms, and tree view clustering according to structure and prediction scores by CFM (Supplementary Figure S7). In the UMAP dataset of GATA2, a key regulator of adipocyte biogenesis, several same-family proteins (e.g., G3BP1, G3BP2), same pathway-proteins (e.g., SF3B3, POLR3A, and IARS1 for transcription and translation, and DHX15, DHX9, RTCB, SNRPA1, and UPF1 for RNA splicing), and subunit proteins of the same complexes (e.g., SSRP1 and SUPT16H for the FACT complex) were relatively closely clustered, as anticipated (Supplementary Figure S8). The GTP-binding protein regulator DRG1, which had the highest CFM score (0.596 out of the maximum score of 1.0) among adipocyte-specific protein–protein interactions with GATA2, was located close to DPYSL2, which plays a role in neuronal development and polarity, in the dendrogram as well as UMAP analysis (Supplementary Figure S7–9, Supplementary Figure S8, Supplementary Table S27). RNA-binding proteins LUC7L2 and RALY were also located in close proximity in UMAP (Supplementary Figure S8). Furthermore, CUL3, a member of the proteasomal degradation pathway, was positioned close to DRG1, also known as NEDD3, in both dendrogram and UMAP analysis and has been reported to interact with the cullin-associated protein, CAND2 (Supplementary Figure S8, Supplementary Figure S7–9, Supplementary Table S27) (*60–62*). They may be involved in the cullin-RING ubiquitin ligase network as a complex (*63*).

Hence, the clustering analyses based on AI inference of HaloMS data may contribute to the knowledge of how multiple TF complexes form to control adipocyte biogenesis and help to illuminate additional hidden features of the protein network that are regulated by previously unknown pathways.

#### HaloMS reveals novel interactions of heterogeneous nuclear ribonucleoproteins (hnRNPs)

The TF-HaloMS dataset includes protein interactions among TFs from several signaling pathways that are specific to the differentiated adipocyte (Supplementary Figure S4, S9A, Supplementary Table S28). Interconnections between the long-noncoding RNA (hereafter “lnRNA”) and hnRNPs in adipocyte biology have been documented (*64*). More than 100 brown-adipocyte lnRNAs, including lncBATE1, were identified as requirements for the establishment and maintenance of brown-adipocyte identity, and that interact with hnRNPU, which is also required for brown adipogenesis (*65, 66*). In the TF-HaloMS dataset generated in the current study, several key regulators of adipocyte biogenesis, namely, ATF2, ATF5, CREB1, PPARA, GATA2, CEBPA, and CEBPD, interact with HNRNP family proteins, HNRNPL and HNRNPU (Supplementary Figure S4, Supplementary Table S28). These interactions were independently detected specifically in the adipocyte, except for the PPARA–HNRNPL interaction, for which interaction was inferred in prostate cancer cells by the affinity capture-RNA method (*67*).

HNRNPU interacts with lnRNA, Blnc1, which has been proposed to up-regulate brown and beige fat thermogenesis and white fat expansion by interacting with the transcription regulator EBF2 and Zbtb7b and thus inhibit adipose tissue inflammation (*64, 66, 68, 69*). Our findings suggest a new network and role for the HNRNP family, especially HNRNPU, interacting with key proteins as a direct regulator of adipocyte biogenesis and maintenance, permitting crosstalk between lnRNA and cell differentiation. HNRNPU interactions with key proteins, such as TFs, might decide the cell fate by controlling access to the genome to specific lnRNA (*65, 68, 70*). This interaction sharing by multiple hnRNPs with a key regulator may provide a route to facilitate the processing of mRNA for cell differentiation signaling.

#### HaloMS reveals novel interactions of ATP-dependent RNA helicase DEAD box proteins (DDXs)

In the obtained TF-HaloMS network, we identified a subnetwork of ATP-dependent RNA helicases, the DDXs. These proteins are involved in RNA metabolism, from transcription to degradation, and are important players in gene expression (*71*). HaloMS revealed novel networks involving DDXs, relative to the previously reported cellular process map (*71*), extending the known network by the interactions of at least six key regulators, GATA2, ATF3, CEBPD, CREB1, CEBPA, and PPARA (Supplementary Figure S9B, Supplementary Table S29). As detected by HaloMS, these six key regulators of adipocyte biogenesis interact with DDX17, DDX39B, DDX46, and DDX5 in an adipocyte-specific manner. Consistent with previous observations (*72–78*), the HaloMS network analysis also revealed molecular links between DDXs (DDX1, DDX41, DDX39B, and DDX17) and key regulators of carcinoma biogenesis (CASP3, CTNNA1, and FOS), although the molecular mechanism of these connections remains unclear. Based on the TF-HaloMS analysis, we propose the existence of cross-regulation of adipocyte differentiation and carcinoma cell turnover signaling via protein–protein interactions between the key regulators and DDXs. Indeed, RNA helicases from the DEAD box family, DDX5 and DDX17, cooperate with hnRNPH/F splicing factors to define cell specificity (*79*). The HaloMS interaction analysis also points to a possible connection between DDX17, HNRNPs (H1/H2/U), and key TFs of adipocyte biogenesis (CEBPA, CEBPD and ATF3; Supplementary Figure S9C, Supplementary Table S30). Since DDX17 functions as a coregulator of master transcriptional regulators of cell differentiation (*79, 80*), these members might comprise the core network of adipocyte differentiation cooperating with lnRNA.

#### HaloMS reveals novel interactions of the spliceosome in adipocyte

The TF-HaloMS pathway analysis of adipocyte- and pre-adipocyte-specific pairs of proteins (Supplementary Figure S4, Supplementary Table S13) identified major spliceosome-associated proteins in both cases (Supplementary Table S16). Comparing the respective cell-specific protein networks, the adipocyte-specific network (72 edges) had approximately 7-fold more interactions than the pre-adipocyte-specific network (10 edges; Supplementary Figure S9D, Supplementary Table S31). In the adipocyte-specific network, SRSF3, SNRPD2, SRSF1, DDX46, SRSF7, U2AF1, PUF60, and HNRNPU shared four or more interactions each with adipocyte regulators. SRSF3 protein is a regulator of brown-adipocyte formation (*81*) and is involved in hepatocyte maturation and metabolism (*82*). SRSF1 generates the PPAR gamma isoform and is involved in pre-adipocyte differentiation (*83*). In addition, the early pre-spliceosomal complex members SRSF7, FUS, DDX46, DDX5, HNRNPA3, and HNRNPU (*84*) were detected in the adipocyte-specific network, suggesting that spliceosome plays an important role in adipocyte differentiation in complex with key adipocyte regulatory factors (*85*).

#### HaloMS reveals novel interactions of diabetic cardiomyopathy network

In the TF-HaloMS dataset of adipocyte- and pre-adipocyte-specific interactions (Figure 3A, Supplementary Table S13), we identified a subnetwork of proteins involved in the diabetic cardiomyopathy pathway (Supplementary Figure S10A, Supplementary Table S32). A considerable relationship between adipose tissue and cardiovascular disease has been reported (*86, 87*), and inadequate adipose tissue function is responsible for cardiovascular events and heart failure in individuals with obesity, metabolic syndrome, and type 2 diabetes (*88*). The HaloMS dataset expanded the known diabetic cardiomyopathy network, relative to interactions previously reported in the literature (Supplementary Figure S10A, Supplementary Table S32), by at least 39 protein interactions (*89*). In this subnetwork, COL3A1 interacts with seven adipocyte regulatory molecules, namely, ATF2, ATF3, ATF5, CEBPA, CREB1, GATA2, and PPARA. According to a previous study, COL3A1 (collagen III protein) is required for adipogenesis with the adhesion G-protein–coupled receptor Gpr56 (*90*), and the protein also induces the progression of ischemic heart failure (*91*). Based on the TF-HaloMS network data, we hypothesize that diabetic cardiomyopathy and adipocyte differentiation occur via protein– protein interactions among three collagen family proteins, eight mitochondrial proteins, and nine master regulators of adipocyte (Supplementary Figure S10A, Supplementary Table S32). Thirty percent of this diabetic cardiomyopathy subnetwork overlapped with the thermogenesis subnetwork (Supplementary Figure S10B, and C, Supplementary Table S32) composed of mitochondrial proteins. These findings point to a novel potential connection between diabetes-related signaling pathways, such as the PPAR signaling pathway, and energy homeostasis (*92*).

#### HaloMS reveals links between endoplasmic reticulum (ER)-phagy, neurodegenerative disease, and adipocyte

Among interactions of a group of proteins not mapped by the aforementioned pathway analysis in the TF-HaloMS dataset (*P* > 0.001, Supplementary Figure S4, Supplementary Table S16), we found a novel subnetwork of proteins involved in ER-phagy, neurodegenerative disease, and myelodysplastic and master regulators in adipocyte (Supplementary Figure S10D, Supplementary Table S33). The relationships between adipose tissue, diabetes, and ER-phagy have been previously explored (*93, 94*). UFL1, which ubiquitinates UFM1, interacts with DDRGK1 and is involved in ER-phagy and in the development of diabetes (*95, 96*). This subnetwork includes DNAJC13 (Parkinson’s disease) (*97*), MAP7D1 (autism) (*98*), EIF2B4 (leukoencephalopathy with vanishing white matter) (*99, 100*), TREX1 (Aicardi–Goutières syndrome, systemic lupus erythematosus, familial chilblain lupus, Cree encephalitis, cryofibrinogenemia, and retinal vasculopathy with cerebral leukodystrophy) (*101*), and LUC7L2 (myeloid malignancies) (*102, 103*). The related genes are involved in neurodegenerative and myelodysplastic diseases, providing new insights as novel network members of the adipocyte master regulator.

Overall, the protein–protein interactions described in the case studies above generated a wide range of hypotheses on the mutual modulation of adipocyte-specific signaling pathways. These and more can be derived from the HaloMS dataset, which demonstrates the power of HaloMS to uncover unexpected network complexity even in characterized signaling systems.

## Conclusion

We describe the development of HaloMS, a HaloTag-based APMS methodology, and its application to map interactions for human TFs involved in adipocyte regulation. Validation in a benchmarked interaction assay demonstrated that the quality of the TF-HaloMS dataset is comparable to that of published interaction datasets. The quality of the data was further confirmed by the validation of a subset of interactions, including benchmarked interaction, by AI. Validation in a pull-down experiment supported the notion that 20% of HaloMS positives were due to direct protein–protein interactions rather than indirect protein–protein interactions. Significant overlap with previously described interactions in a public database of human proteins indicated that HaloMS provided a high-quality protein–protein interaction dataset. Despite the statistically reliable overlap of HaloMS with the *Arabidopsis* dataset, the HaloMS overlap was moderate, indicating that the HaloMS dataset reflects the findings of the Y2H assay, which is the main data source in the *Arabidopsis* dataset (*17, 104*) and may suggest that fewer data may be obtained by APMS interaction assays in plants. Further, the moderate finding of reproducibility by AI approaches suggests that the interactions detected by HaloMS are novel and scarce in the AI training dataset.

Protein arrays, which use the same protein capturing method as HaloMS, are based on *in vitro* synthesis of target proteins and do not consider their expression levels in the cell, resulting in a constant copy number of target proteins. This may explain why target proteins involved in “transcription,” such as TFs with a low-expression level in the cell, are more likely to interact with TFs as query proteins. However, HaloMS uses cellular protein extracts as the source of targets, which may reflect the dynamic range of proteins expressed in the cell.

HaloMS offers several advantages over conventional protein interaction assays, such as Y2H, protein arrays, and one-by-one APMS. The high-throughput synthesis and capture of bait proteins *in situ* eliminate the need for individual in-cell protein expression and purification.

Additionally, because native samples are used as targets, rather than proteins expressed *in vitro* or in yeast, complexes composed of cofactors other than proteins, such as TF–RNA-binding protein–lnRNA complexes, can be detected. In addition to cultured cells, HaloMS can be readily applied to other samples, particularly human organs or tissues for which query transformations are difficult and to diseased specimens for which intact conditions are examined. We envision that proteome-scale HaloMS could be adapted for other sample targets, such as brown, white, beige adipocyte, and non-transformable diabetic specimens. Biologically, the TF-HaloMS dataset confirms known interactions, provides biochemical support for genetic and molecular biological observations, and illuminates previously unknown mutual regulation of adipocytes signaling pathway.

### Limitations of the study

HaloMS has several potential limitations. In the current study, we identified a large number of cell type-specific interaction partners for the 17 query proteins. This was somewhat surprising. An important reason for the large number of interactions could be the selection of bait proteins that are critical components of some of the most vital adipocyte regulation processes. By screening TFs of adipocyte central signaling pathways against different cell types, it is likely that we inadvertently identified cell type-specific connected TF-related proteins and family members, such as hnRNPs and DDXs. These differentiated adipocyte-specific TFs and TF-related proteins have intrinsically disordered regions, and their network members may aggregate to transcriptionally regulate adipocyte differentiation (*105–109*). HaloMS functions by immobilizing query proteins to a solid support, which may limit the accessibility of the bait proteins to target proteins from the cell lysate. Thus, once interactions are identified despite the anchoring proteins, these interactions can be validated by the second assay, including AI technology. The new technology described in this study has the potential to yield significant results in mapping transcriptional protein complexes in patients; however, its detection is limited by the minimal amount of protein input necessary to detect protein–protein interactions. There is a sensitivity limitation in this system when studying these transcriptional protein complexes in rare/limited cell populations that may not be abundant in healthy donors or patients. In such cases, ultra-sensitive mass spectrometry methods have been developed that can identify trace samples and should be used for relevant analyses (*110*).

## ACKNOWLEDGEMENTS

Ministry of Education, Science, Sports and Culture, Grant-in-Aid for Scientific Research (C), 2019-2021 [19K0574 to J.Y.]. Takeda Science Foundation, Visionary Research Grant to J.Y. Japan Diabetes Foundation, Novo Nordisk Pharma Ltd., research grant 2022 to JY. We would like to thank Editage (www.editage.jp) for English language editing.

## DATA AVAILABILITY

Plasmids generated in this study are available upon request.

These plasmid vectors below are available from the Arabidopsis Biological Resource Center (ABRC; https://abrc.osu.edu/).

pIX-Halo:ccdb destination vector: https://www.arabidopsis.org/servlets/TairObject?type=vector&id=1001200298 pIX-3×HA:ccdB destination vector: https://www.arabidopsis.org/servlets/TairObject?type=vector&id=1001200524

MS data generated in the current study were deposited in JPOST under the accession number PXD041085 (https://repository.jpostdb.org/; link to preview; https://repository.jpostdb.org/preview/1533709895647ecfc437f55. Access key; 2252). The code used in the current study has been deposited as DOIx and is publicly available as of the date of publication. Additional information needed to reanalyze the data reported in this work is available from the corresponding author upon request.

## SUPPLEMENTARY DATA

Supplementary Data are available at NAR online.

## AUTHOR CONTRIBUTIONS

Junshi Yazaki: Conceptualization, Investigation, Methodology, Data analysis, Writing – original draft. Takashi Yamanashi: Methodology, Data analysis, Writing – original draft. Shino Nemoto: Investigation, Writing – original draft. Atsuo Kobayashi: Investigation. Yong-Woon Han: Tomoko Hasegawa: Investigation. Akira Iwase: Investigation, Writing – original draft. Masaki Ishikawa: Investigation. Ryo Konno: Investigation, Data analysis. Koshi Imami: Data analysis. Yusuke Kawashima: Conceptualization, Investigation, Methodology, Data analysis, Writing – original draft. Jun Seita: Conceptualization, Data analysis.

## Abbreviations

(APMS): affinity purification with mass spectrometry
(hereafter AFM): AlphaFold-Multimer
(AI): artificial intelligence
(BCA assay): Bicinchoninic acid assay
(CEBPs): ColabFold Multimer
(CFM): CCAAT/enhancer-binding protein
(DDXs): DEAD box proteins
(DMEM): Dulbecco’s Modified Eagle Medium
(FDR): false-discovery rates
(FBS): fetal bovine serum
(GO): gene ontology
(HaloMS): HaloTag-based affinity purification Mass Spectrometry
(hnRNPs): heterogeneous nuclear ribonucleoproteins
(hADSCs): Human adipose-derived stem cells
(HEK): Human Embryonic Kidney cell
(ipTM): interface-predicted template modeling scores
(LC-MS/MS): liquid chromatography-tandem MS
(lnRNA): long-noncoding RNA
(PBST): PBS with 0.1% (v/v) Tween 20
(PPARs): peroxisome proliferator-activated receptor
(PBS): phosphate-buffered saline
(pTM): predicted template modeling scores
(SD): standard deviation
(TFs): transcription factors
(UMAP): uniform manifold approximation and projection

---

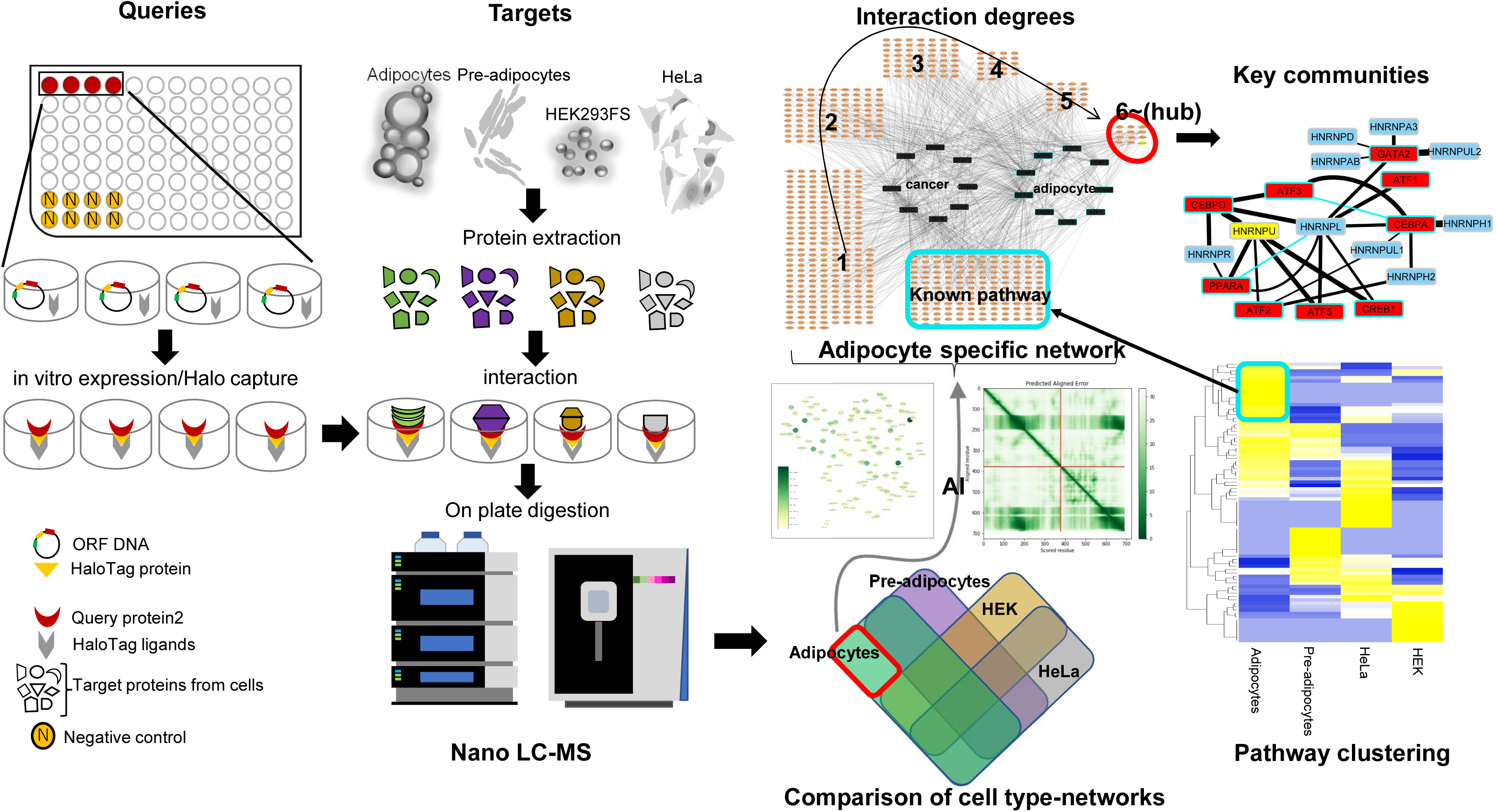

## Supplementary Materials for MAPPING ADIPOCYTE INTERACTOME NETWORKS BY HALOTAG-ENRICHMENT–MASS SPECTROMETRY

**Figure S1.**
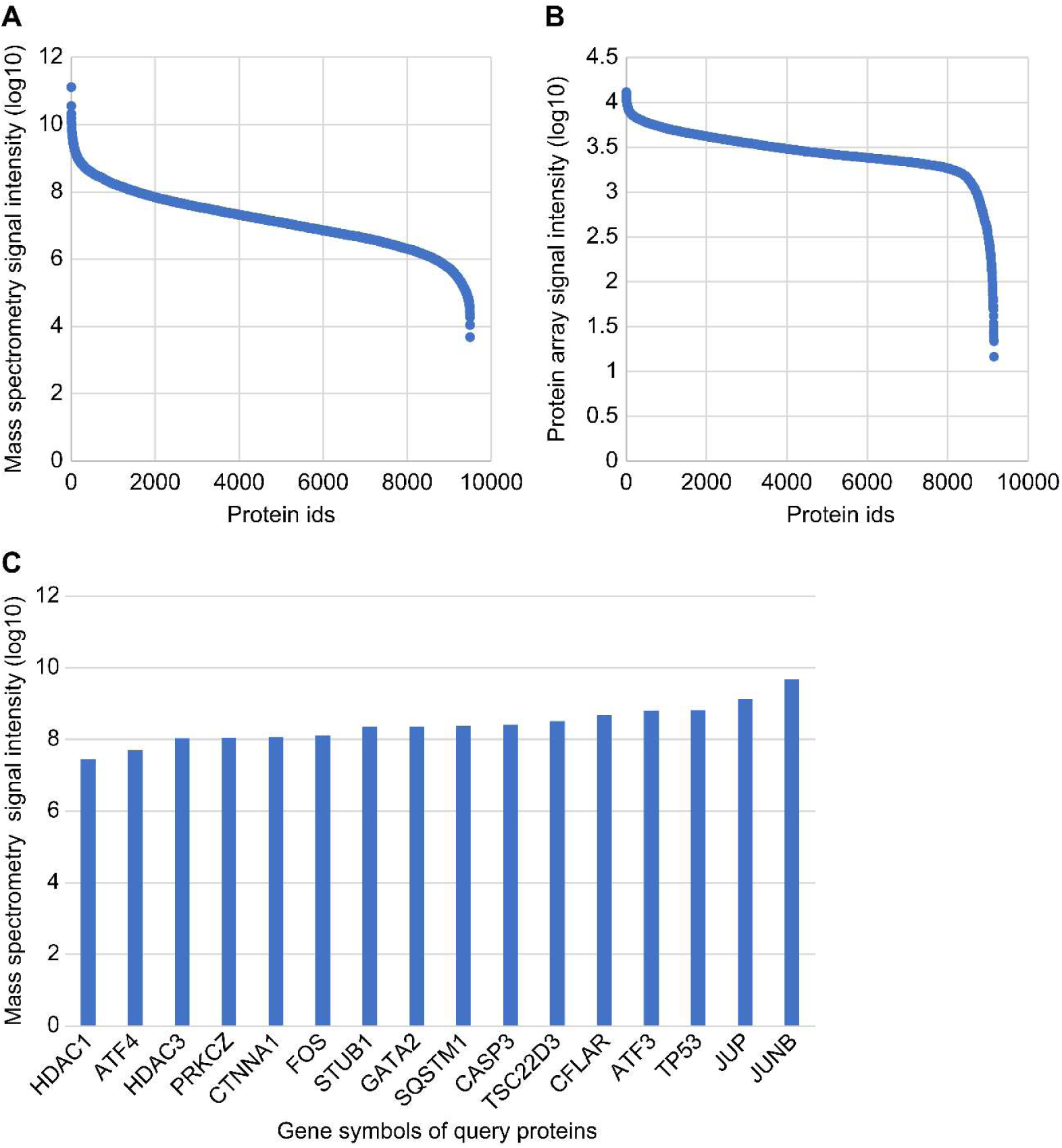
Related to Figure 1. Comparison of two different expression assays. Global protein expression levels are determined by (A) mass spectrometry and (B) protein array (reference *15*) detection with an anti-HaloTag antibody. The sorted histogram (blue data points) indicates log_10_ signal intensity of protein from an extract of mature leaves of *Arabidopsis thaliana* wild type Columbia in (A) and from all ORF clones on the array in (B). Comparison of the 9999 proteins detected by MS to those loaded on the protein array (reference *15*) shows a 63% overlap. (C) Related to Supplementary Table S3. Consistent expression of 16 human HaloTag query proteins immobilized on a ligand-coated 96-well plate. Gene symbols of *in vitro* expressed and captured proteins are shown along the x-axis. Signal intensity from mass spectrometry (log_10_) is shown along the y-axis.

**Figure S2.**
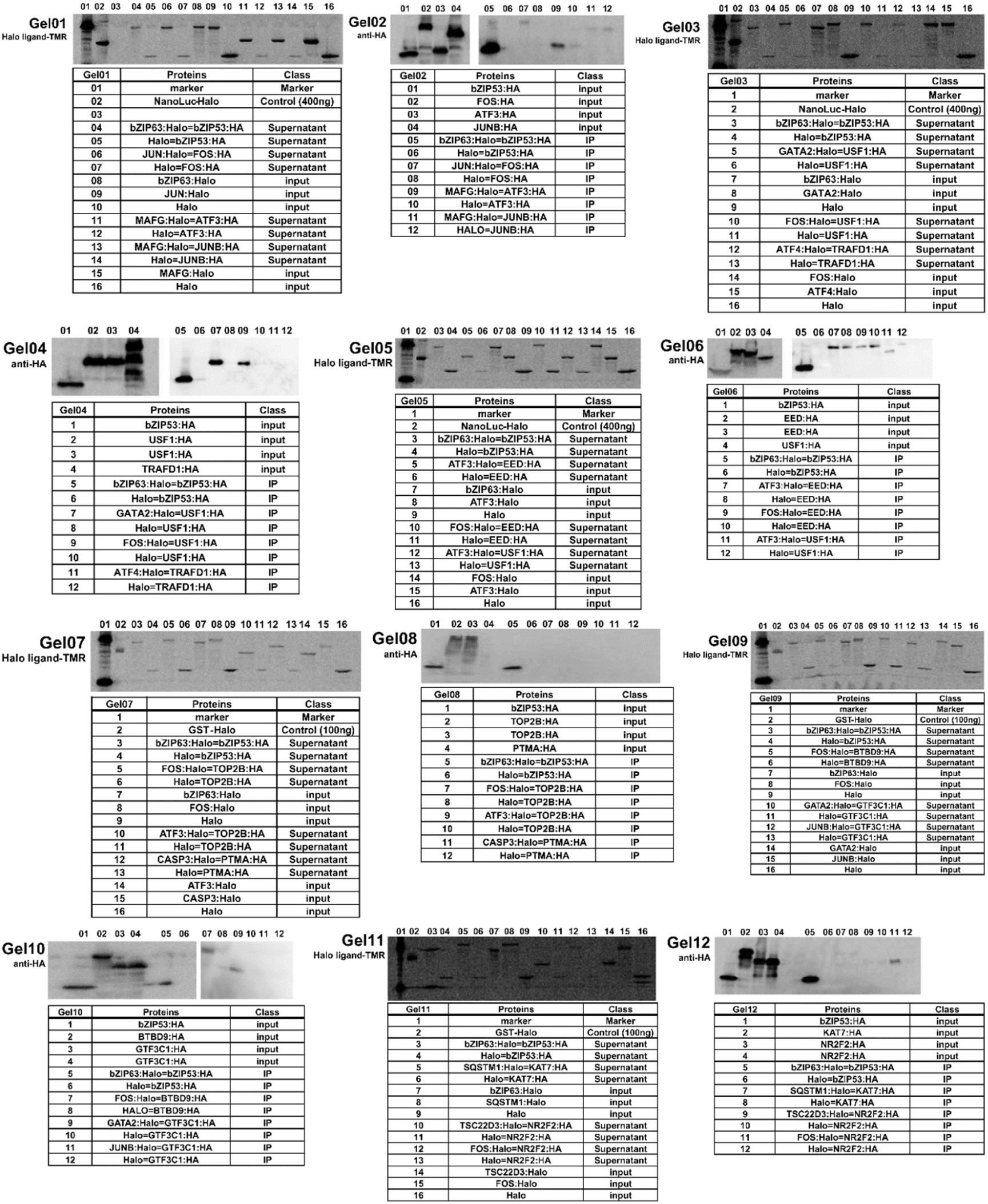
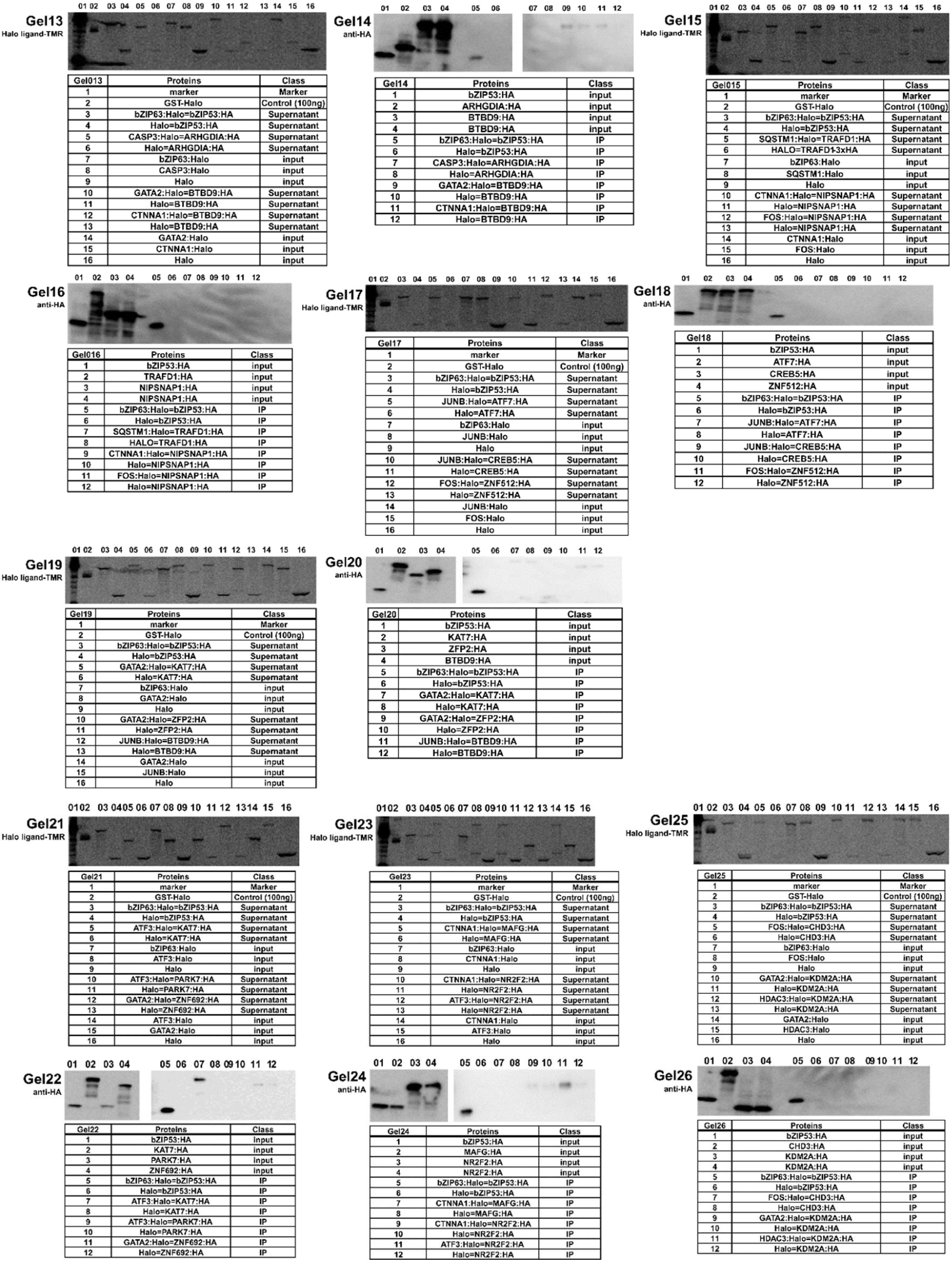
Related to Figure 2A. HaloMS protein–protein interactions are replicated by pull-down assay. Representative results of the *in vitro* pull-down experiments for Halo alone, Halo-tagged, and 3×HA-tagged proteins are shown. Co-purified Halo-tagged and 3×HA-tagged proteins are detected using an anti-HA antibody (labeled as “IP”); 5% of input (labeled “input”) and 5% of supernatant after binding to HaloLink magnetic beads (labeled “Supernatant”) are tested to determine the relative amounts of Halo-tagged and HA-tagged proteins, and the binding efficiency to HaloLink magnetic beads in the experiments. A detailed description of all gel lanes is given in Supplementary Table S8.

**Figure S3.**
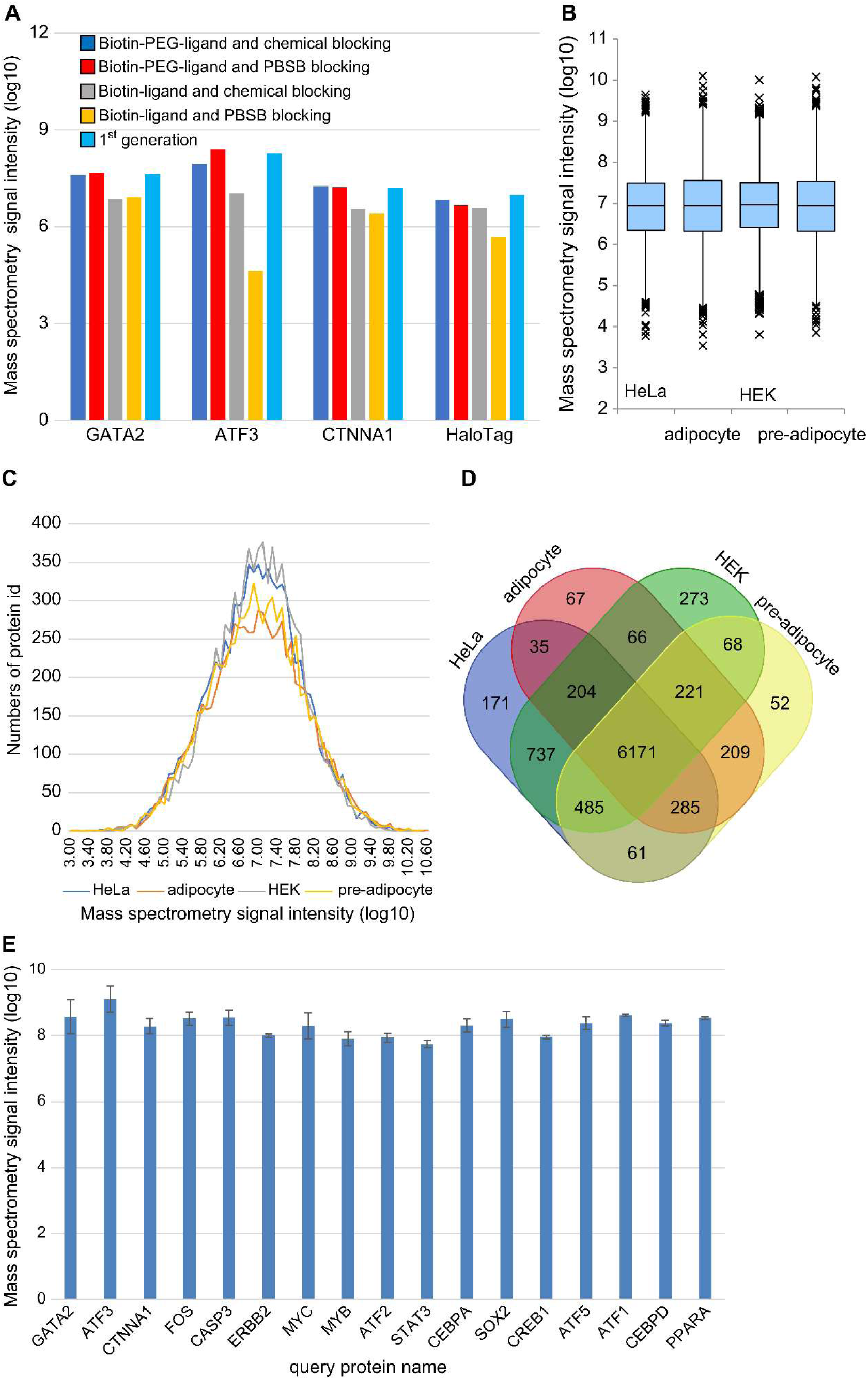
Related to Figure 3A. (A) Inconsistent expression of four HaloTag query proteins immobilized on different ligand-coated 96-well plates. Gene symbols of *in vitro* expressed and captured proteins are shown along the x-axis. Each colored histogram indicates different ligand-coating conditions. Signal intensity from mass spectrometry (log_10_) is shown along the y-axis. The ‘1^st^ generation’ indicates the commercial HaloTag ligand plate (Promega, HaloLink 96 Well Plate, cat no CS180802) used for the initial HaloMS. ‘Biotin-PEG-ligand’ indicates HaloTag-PEG-biotin ligand (Promega, cat. no. G859A) in the method section. The ‘chemical blocking’ indicates Blockmaster reagents (Medical & Biological Laboratories, cat. no. J-PA1080). ‘Biotin-ligand’ indicates HaloTag-biotin ligand (Promega, cat. no. G828A). ‘PBSB’ indicates PBS buffer containing 10 mg/mL of BSA. (B) Proteome profiling of four different cell types. Box plot distribution of peptide signal intensities from four different cell types. The legend for the boxplot includes the following elements: the upper boundary line represents the third quartile, the lower boundary line represents the first quartile, the line within the box represents the median, and any individual dots outside the box represent outliers. All four boxplots show a resemblance, suggesting a uniform trend across the data distributions. (C) Normal distribution of protein expression by each cell type. (D) Venn diagram of the protein expression status of the four cell types. Venn diagram is generated using software available at https://bioinformatics.psb.ugent.be/webtools/Venn/ (E) The intensities of query proteins immobilized on the 96-well plate and used for the screen.

**Figure S4.**
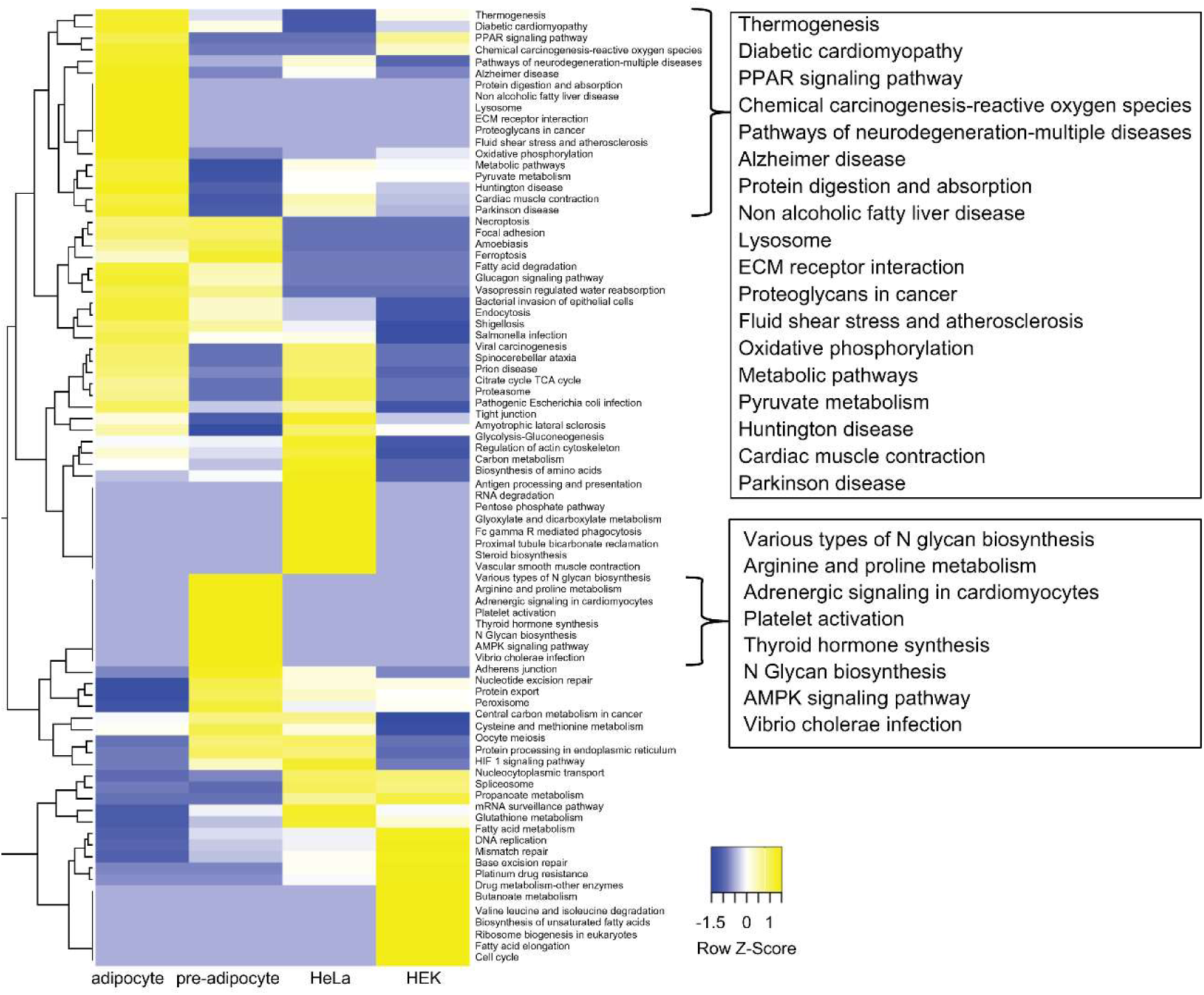
Pathway analysis of four cell type-specific pairs of proteins in the HaloMS dataset. Interacting proteins detected in each cell-specific interaction are enriched and classified by pathway analysis (https://david.ncifcrf.gov/). The heat map is generated by obtaining the z-score from the *P*-values of the enriched classes and reveals the presence of cell-specific subnetworks (http://www.heatmapper.ca/expression/). Clustering is performed by average linkage using the Euclidean distance measure. Samples that are not enriched in a pathway and for which no *P*-value is obtained have an assigned *P*-value (–log_10_) of 0 (Supplementary Tables S15 and S16).

**Figure S5.**
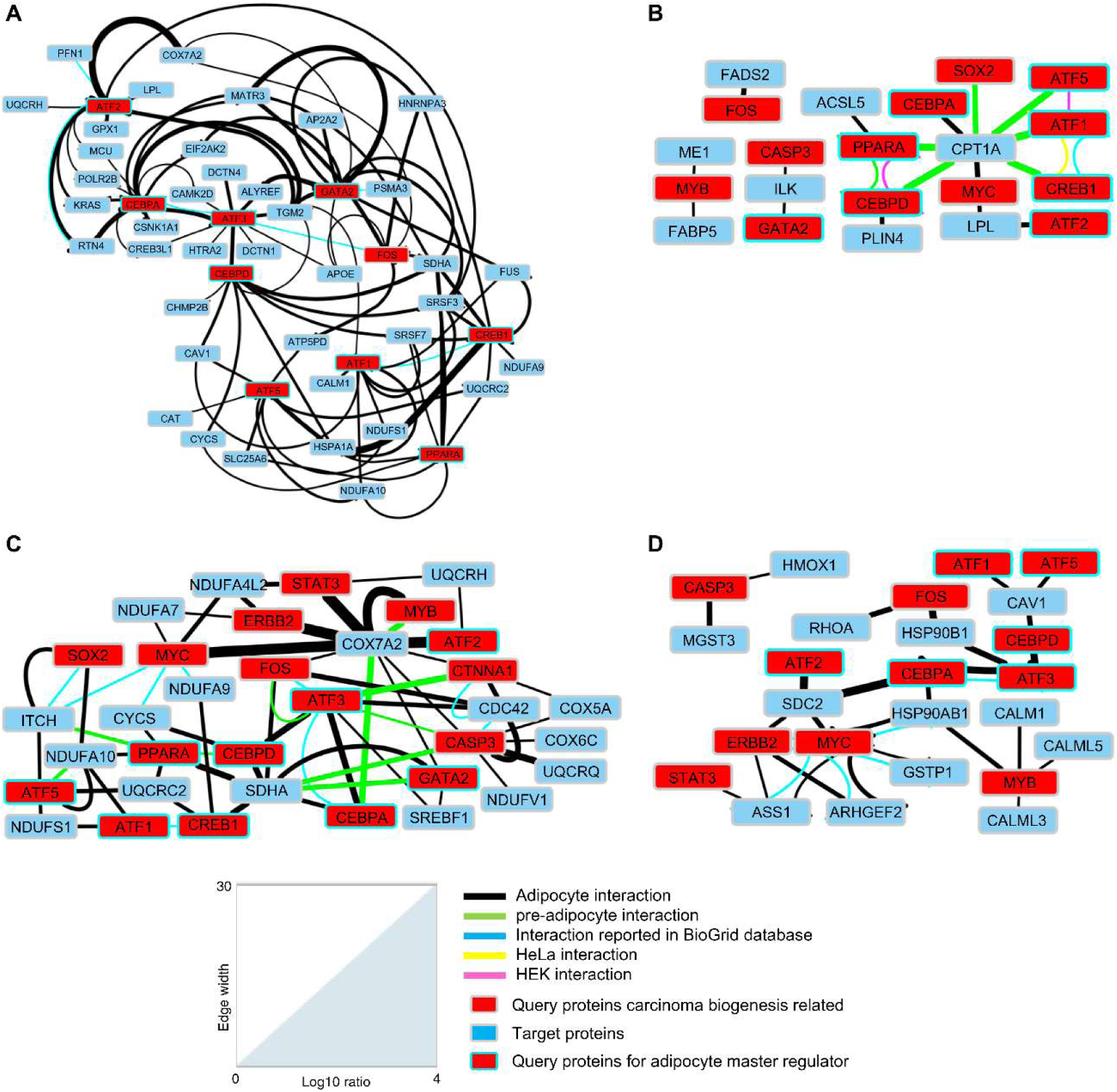
Related to Figure S4. A subnetwork within the adipocyte-specific interaction network supporting the validity of detected interactions in the current study. (A) The adipocyte-specific interaction network contains a subnetwork of mitochondria-associated neurodegenerative disease pathways, including amyotrophic lateral sclerosis, Alzheimer’s, Parkinson’s, Huntington’s, and prion diseases. Activation of these pathways implies activation of the thermogenesis subnetwork in adipocytes. Protein pairs of the pathway are listed in Supplementary Table S17. (B) Detection of a well-known PPAR signaling pathway in adipocytes suggests the validity of the HaloMS dataset. Protein pairs of the pathway are listed in Supplementary Table S18. (C) The adipocyte-specific interactions showing enrichment of terms for non-alcoholic fatty liver disease support the validity of the detected interactions in the current study. Protein pairs of the pathway are listed in Supplementary Table S19. (D) Fluid shear stress and atherosclerosis proteins in adipocytes, suggesting the significance of the HaloMS data set. Protein pairs of the pathway are listed in Supplementary Table S20. The thickness of the edge indicates the log_10_ ratio in the indicator.

**Figure S6.**
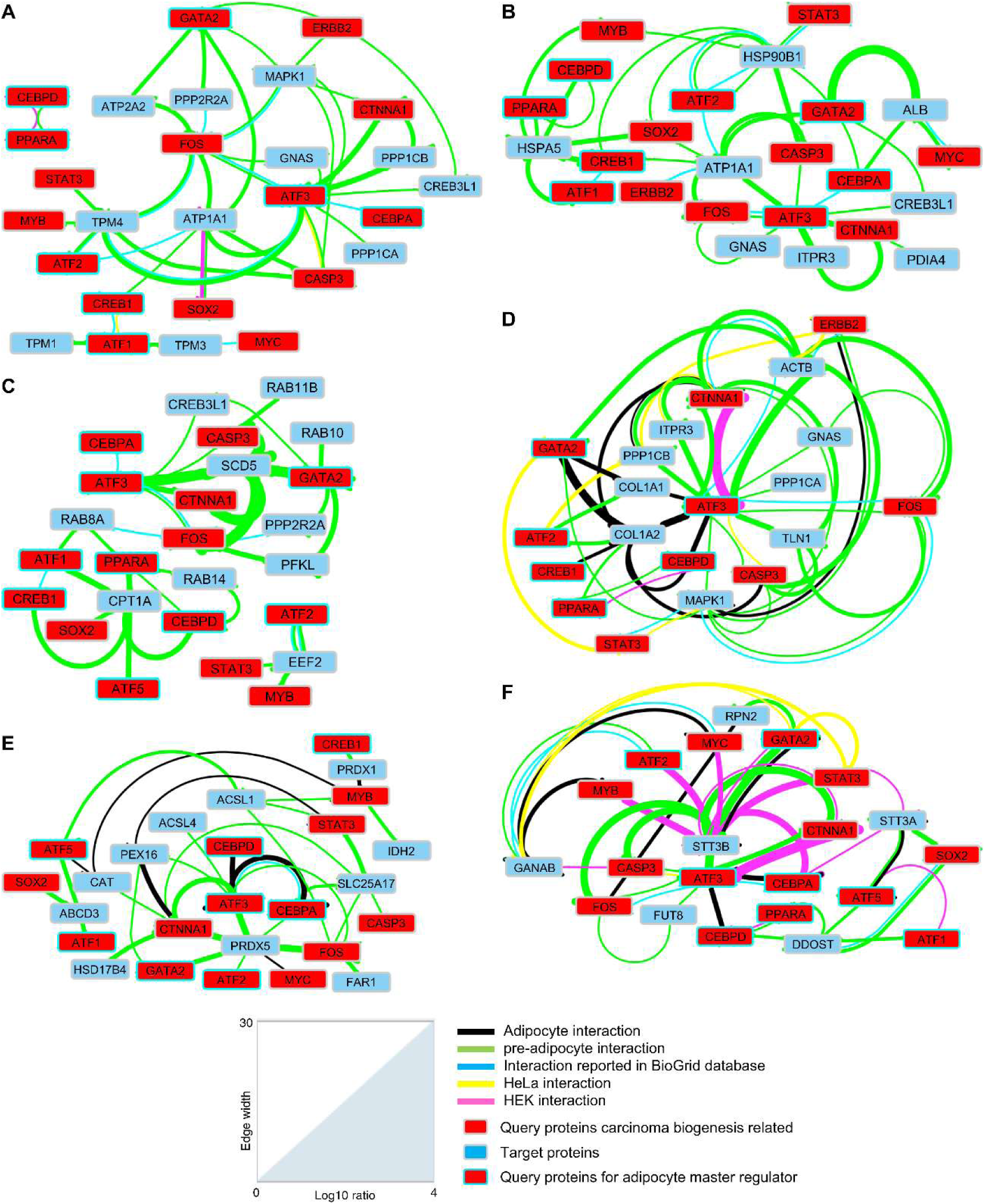
Related to Figure S4. A subnetwork within the pre-adipocyte-specific interaction network supports the validity of interaction detection. (A) Pathway for adrenergic signaling in cardiomyocytes in pre-adipocytes, suggesting the significance of the HaloMS data set. Protein pairs of the pathway are listed in Supplementary Table S21. (B) Pathway for thyroid hormone synthesis in pre-adipocytes, suggesting the significance of the HaloMS data set. Protein pairs of the pathway are listed in Supplementary Table S22. (C) Pathway for AMPK signaling in pre-adipocytes, suggesting the significance of the HaloMS data set. Protein pairs of the pathway are listed in Supplementary Table S23. (D) Pathway for platelet activation in pre-adipocytes, suggesting the significance of the HaloMS data set. Protein pairs of the pathway are listed in Supplementary Table S24. (E) Pathway for peroxisome in pre-adipocytes, suggesting the significance of the HaloMS data set. Protein pairs of the pathway are listed in Supplementary Table S25. (F) Pathway for N-glycan biosynthesis in pre-adipocytes, suggesting the significance of the HaloMS data set. Protein pairs of the pathway are listed in Supplementary Table S26. The thickness of the edge indicates the log_10_ ratio in the indicator.

**Figure S7.**
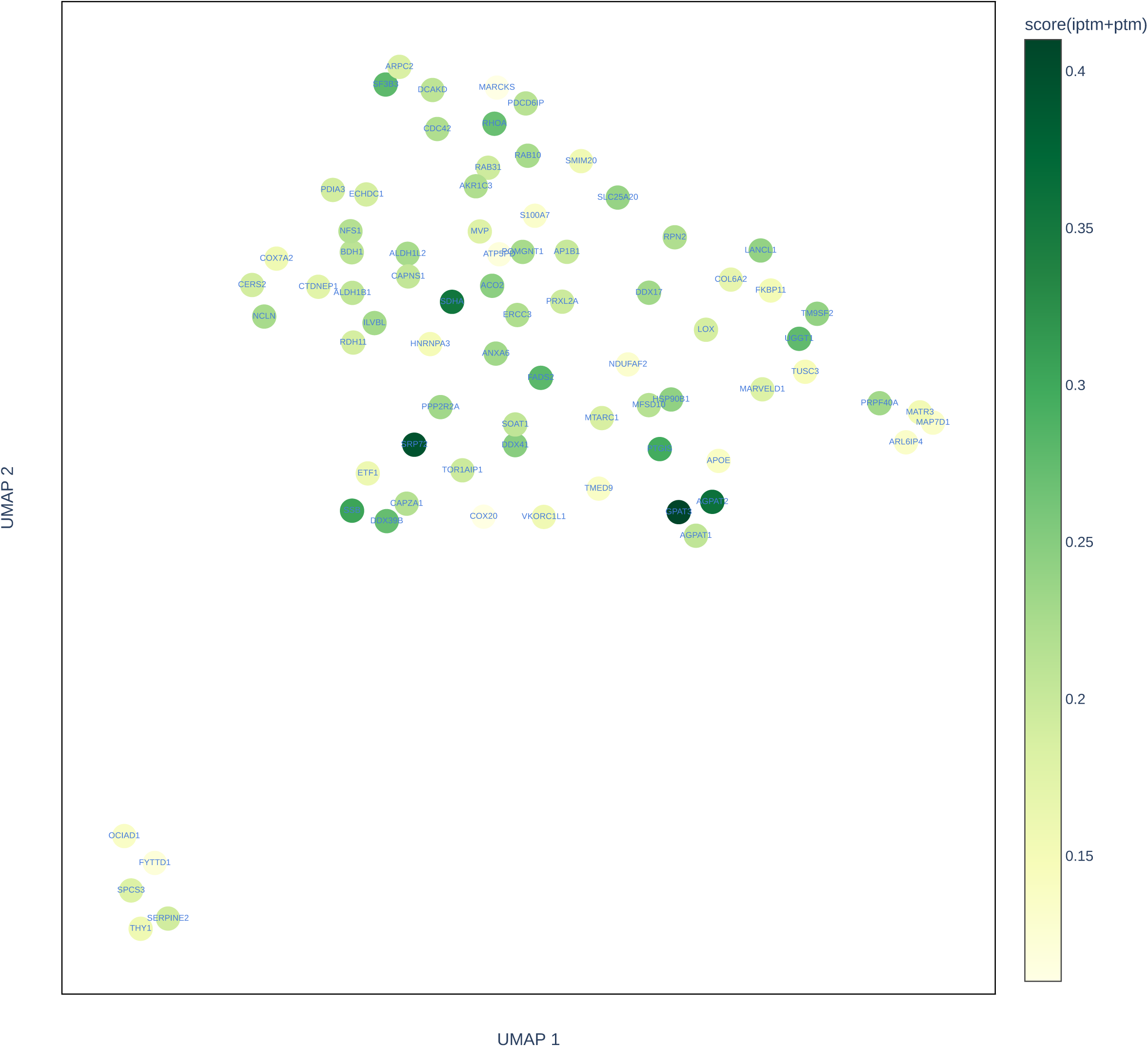
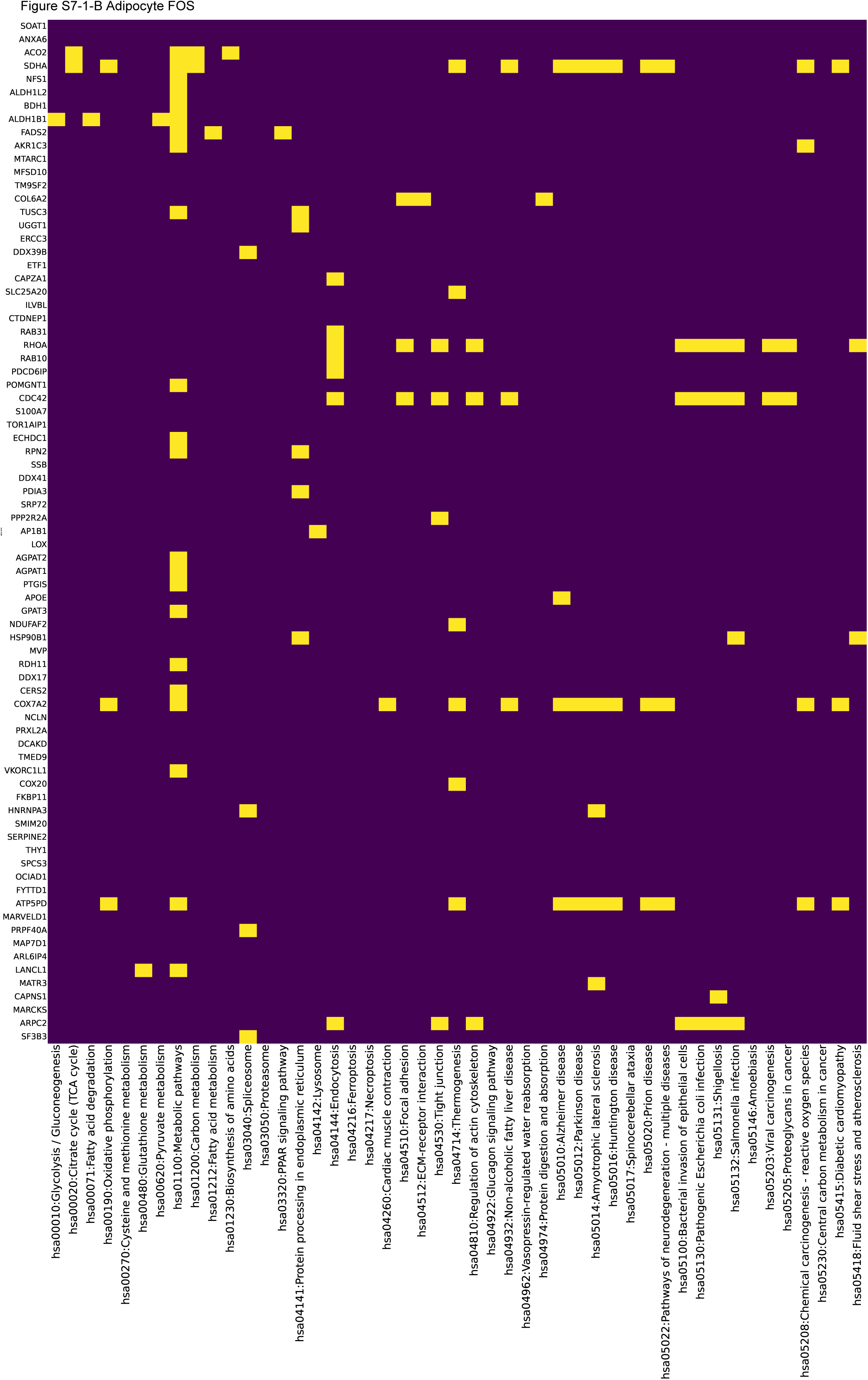
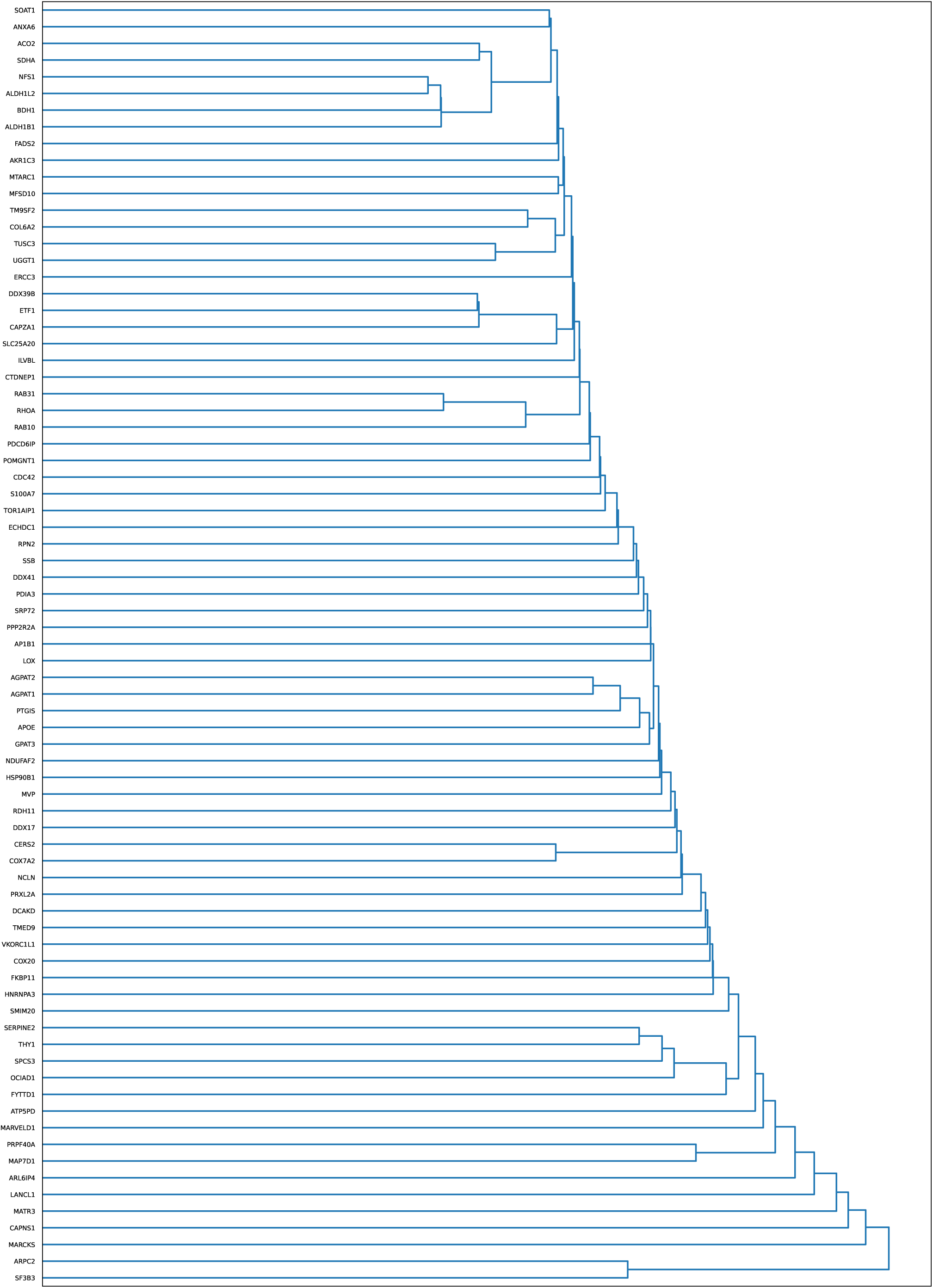
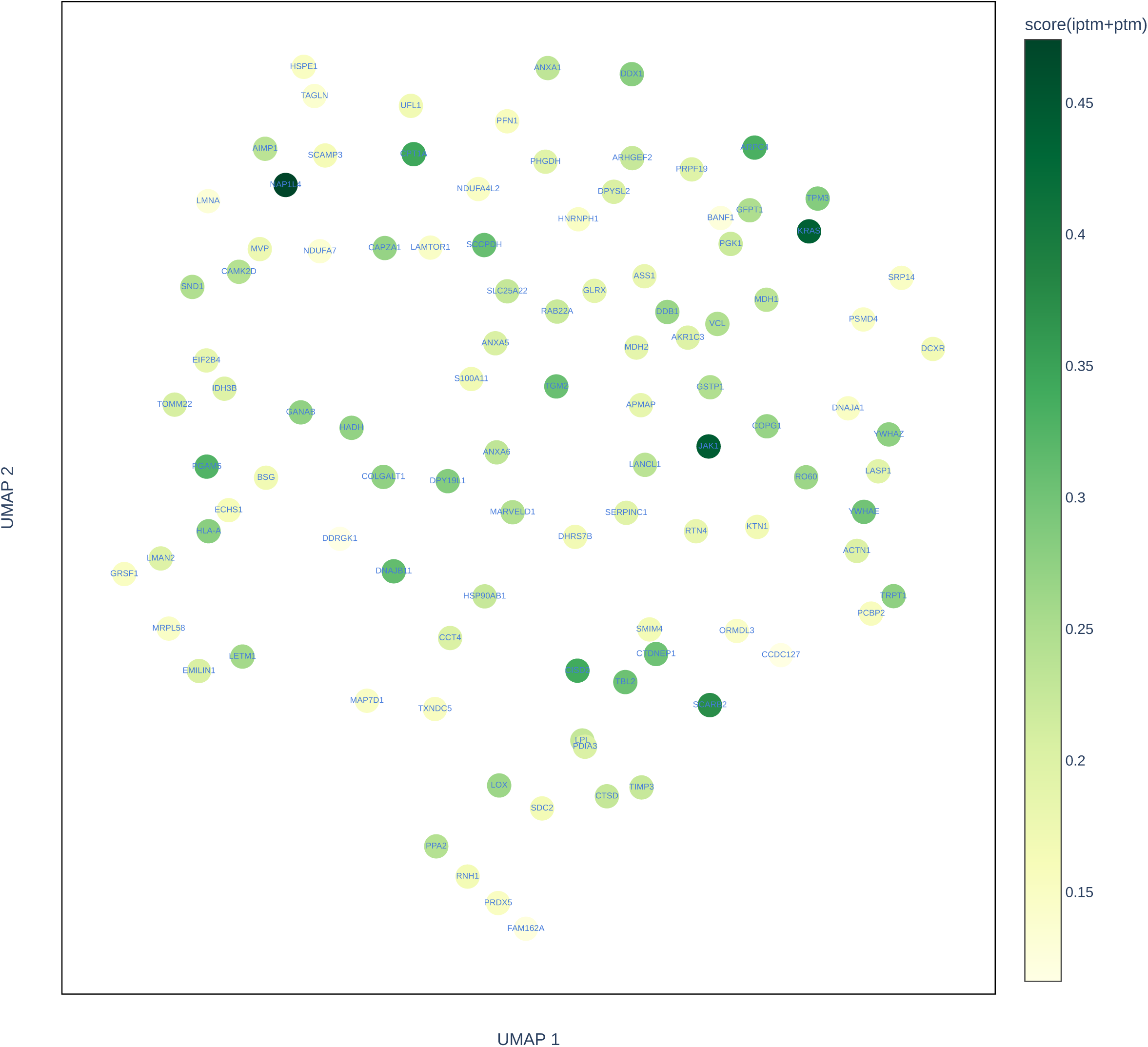
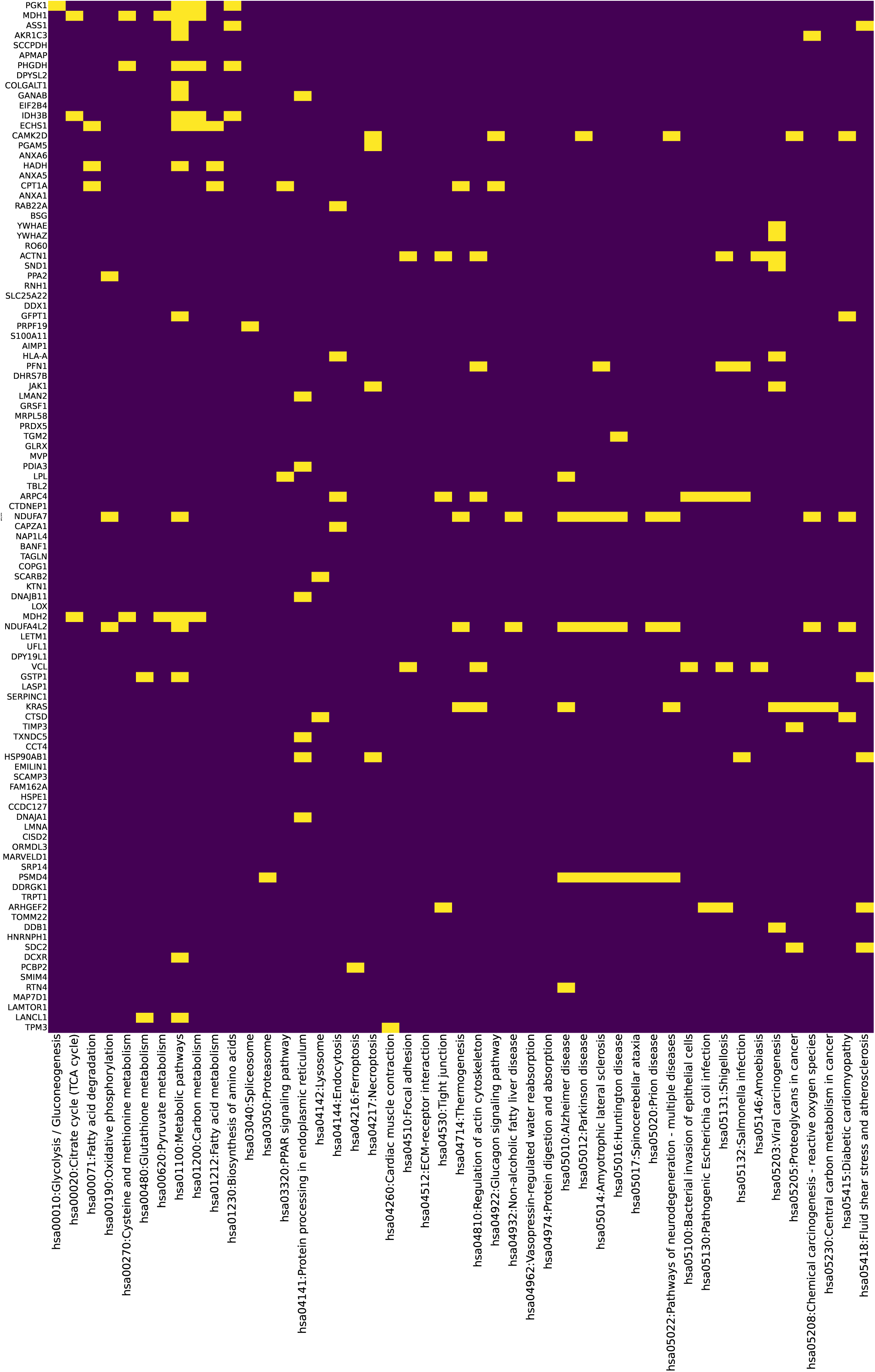
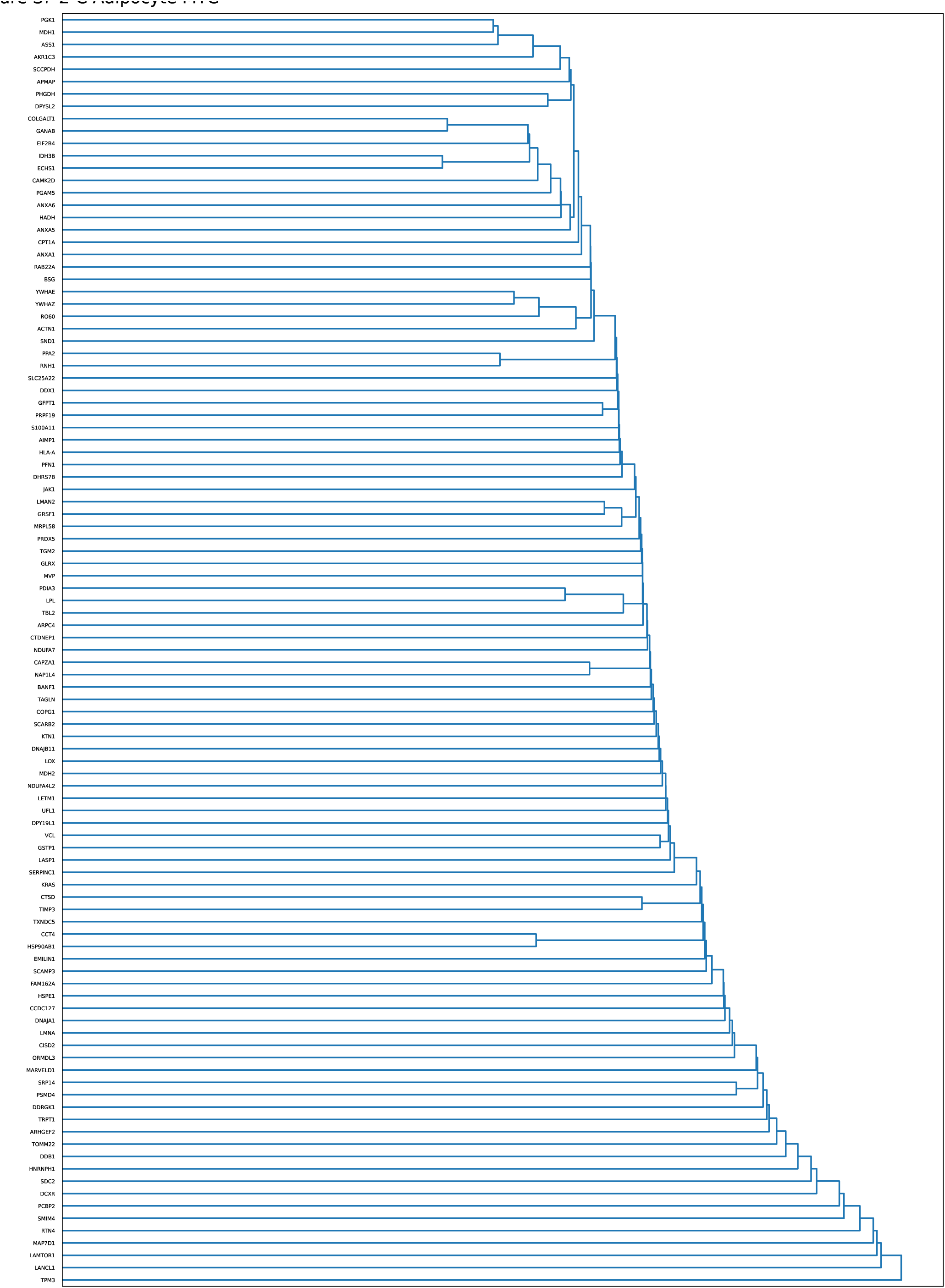
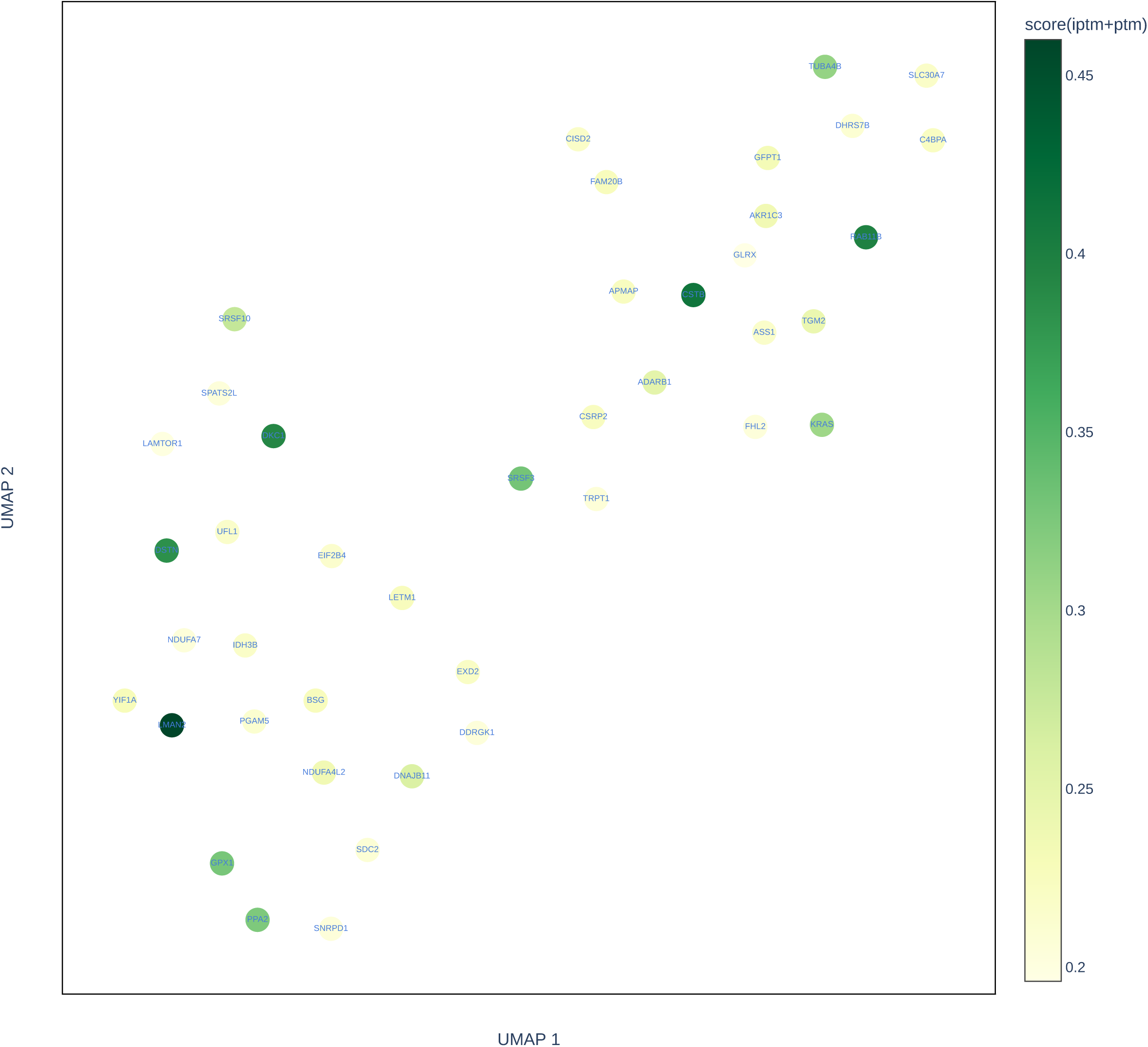
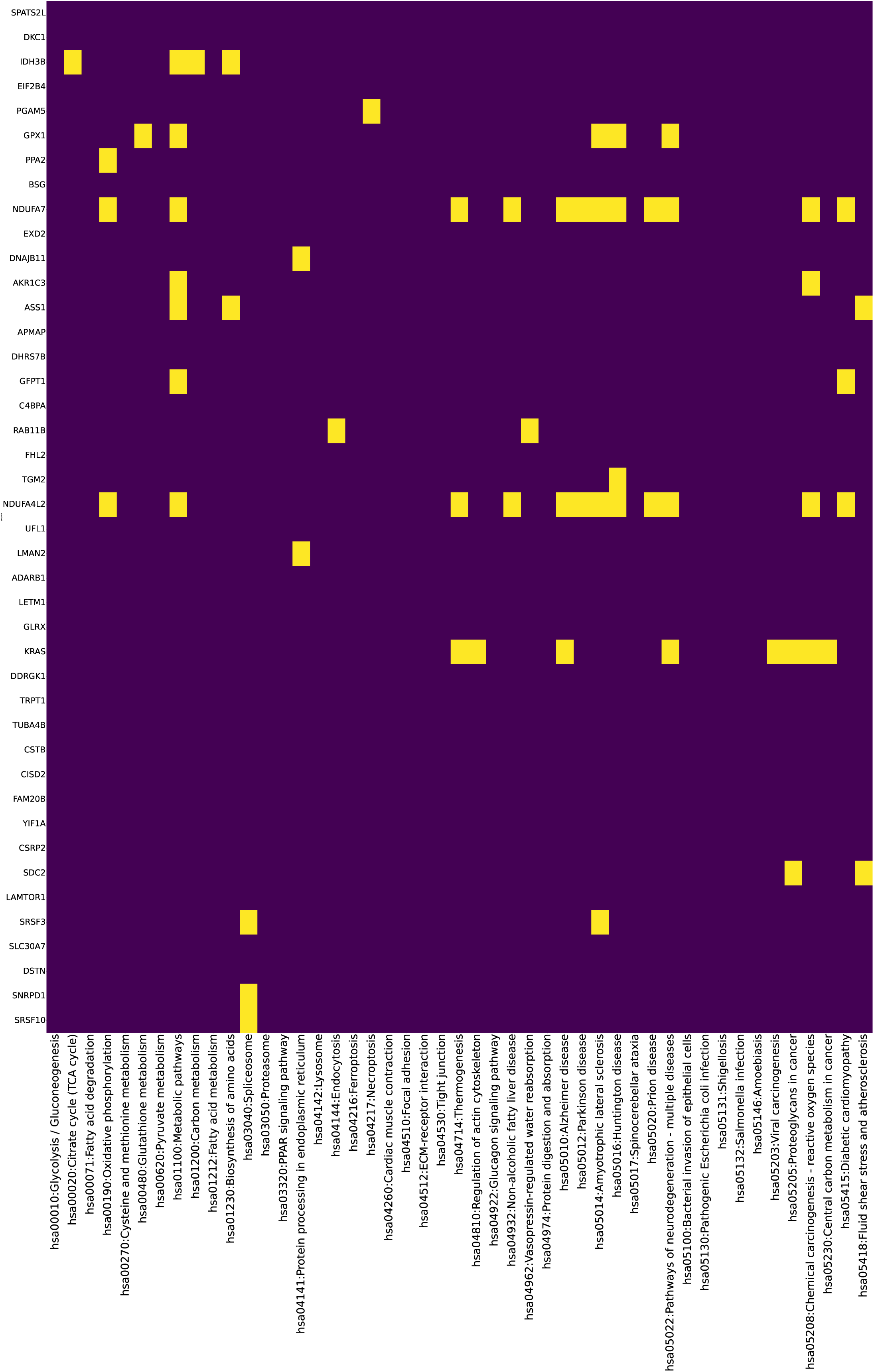
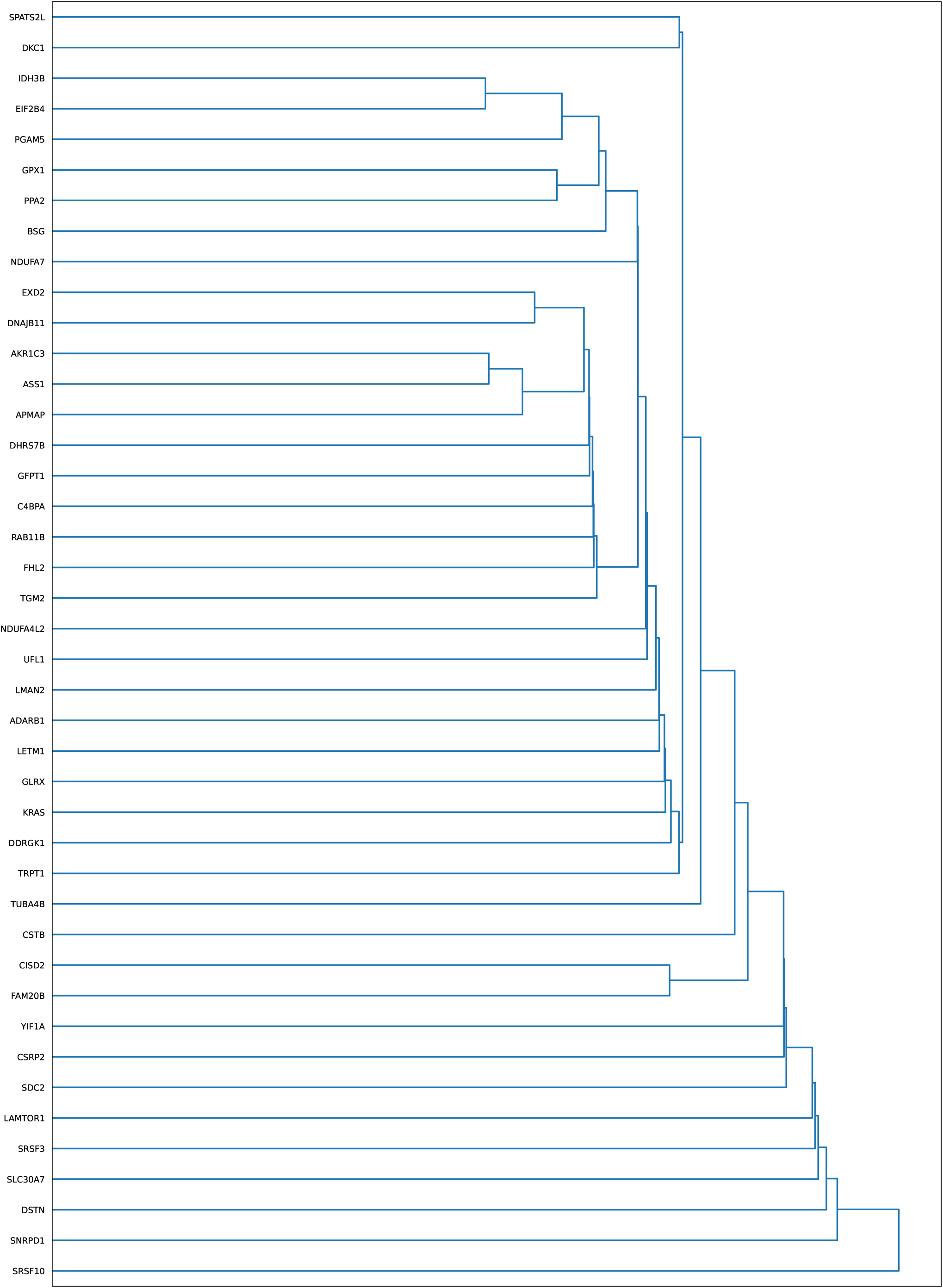
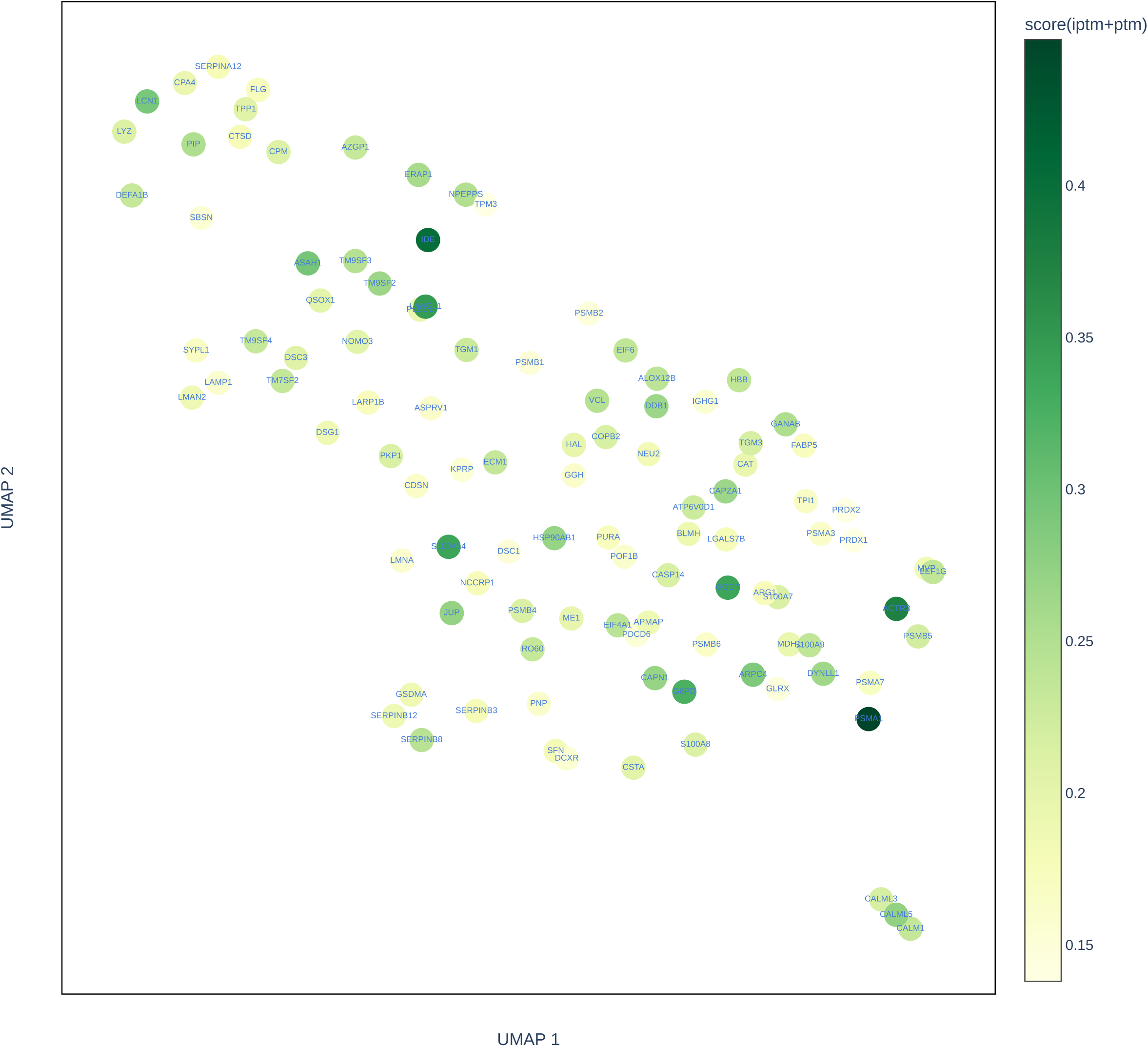
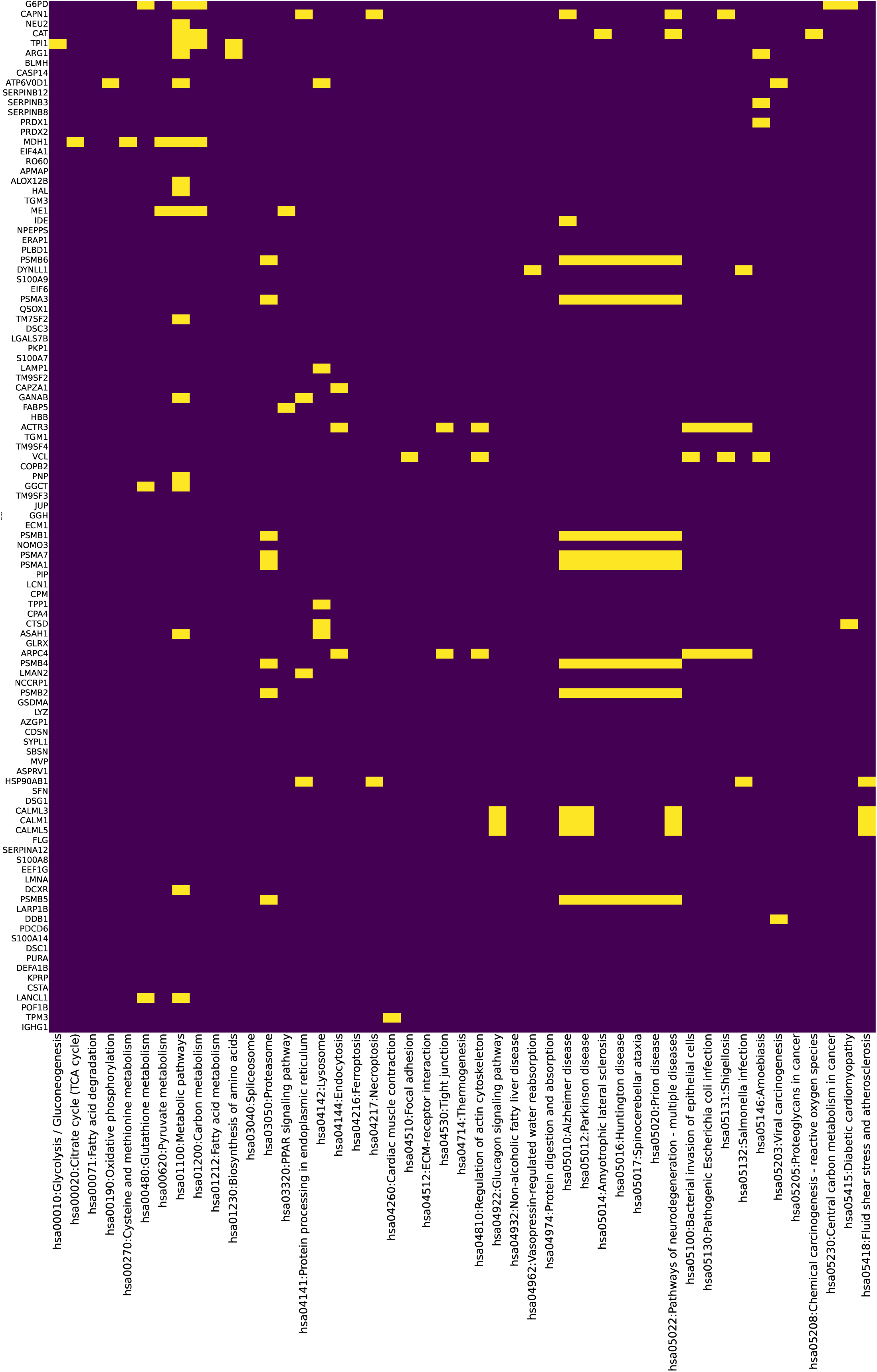
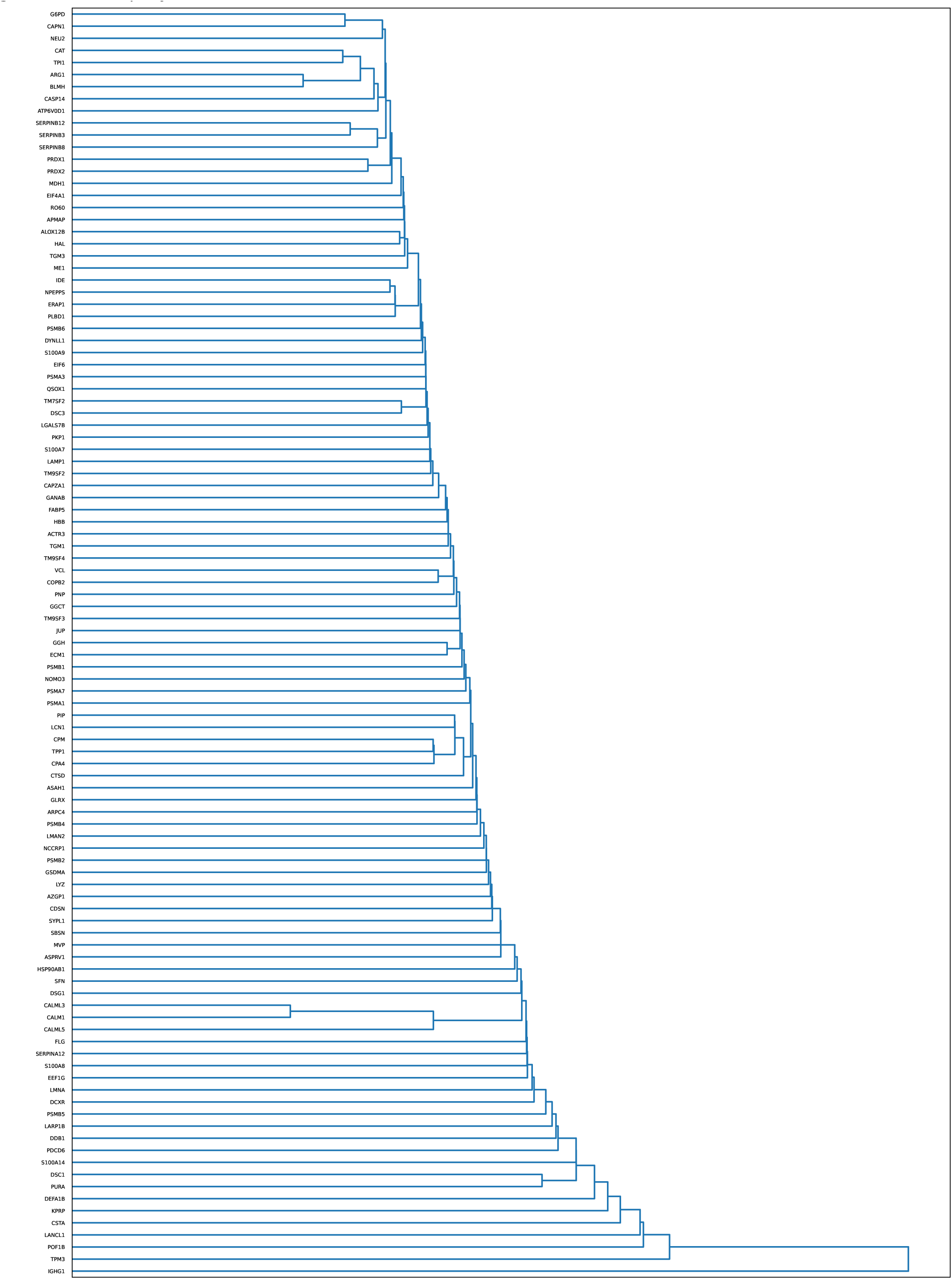
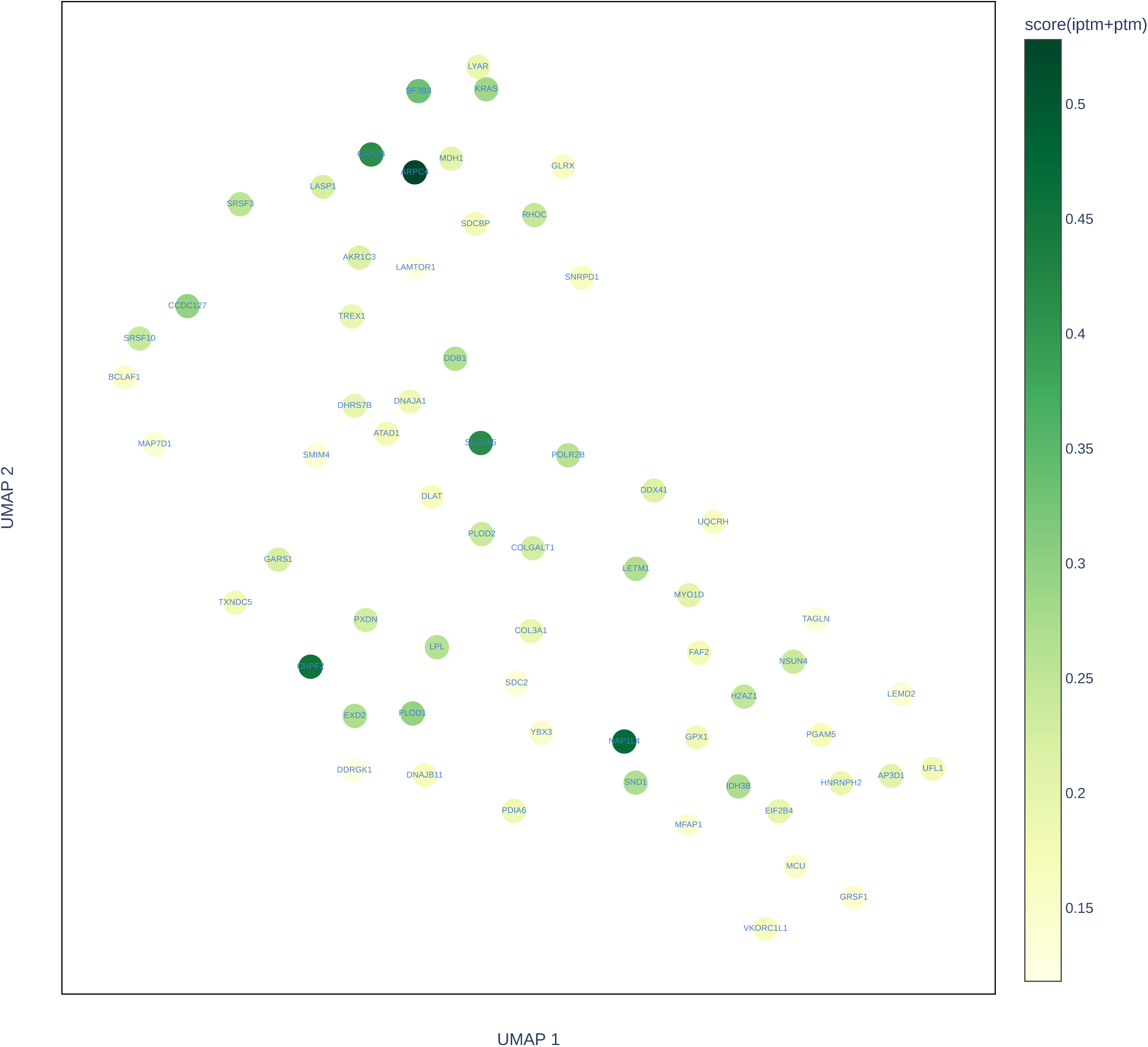
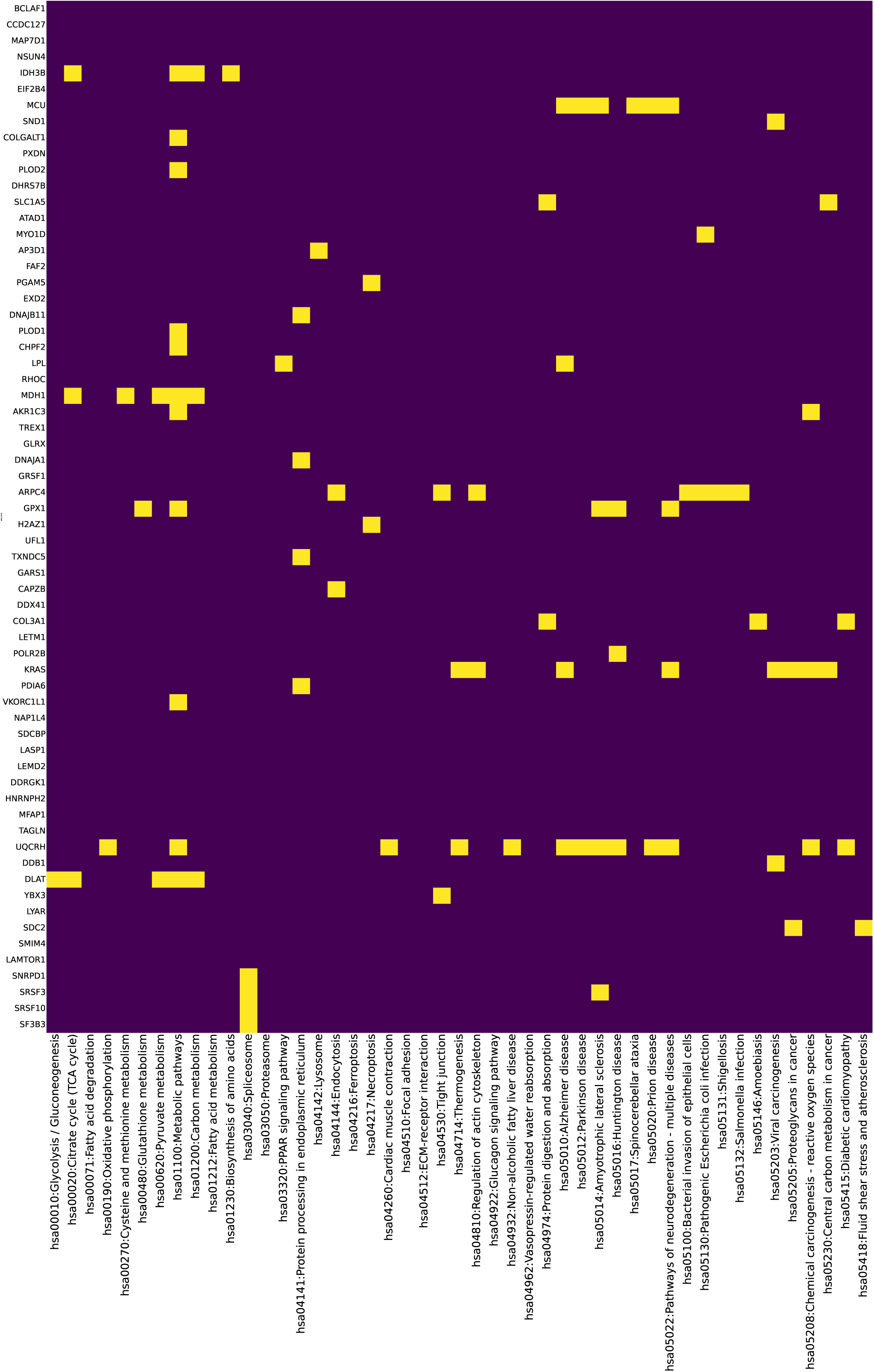
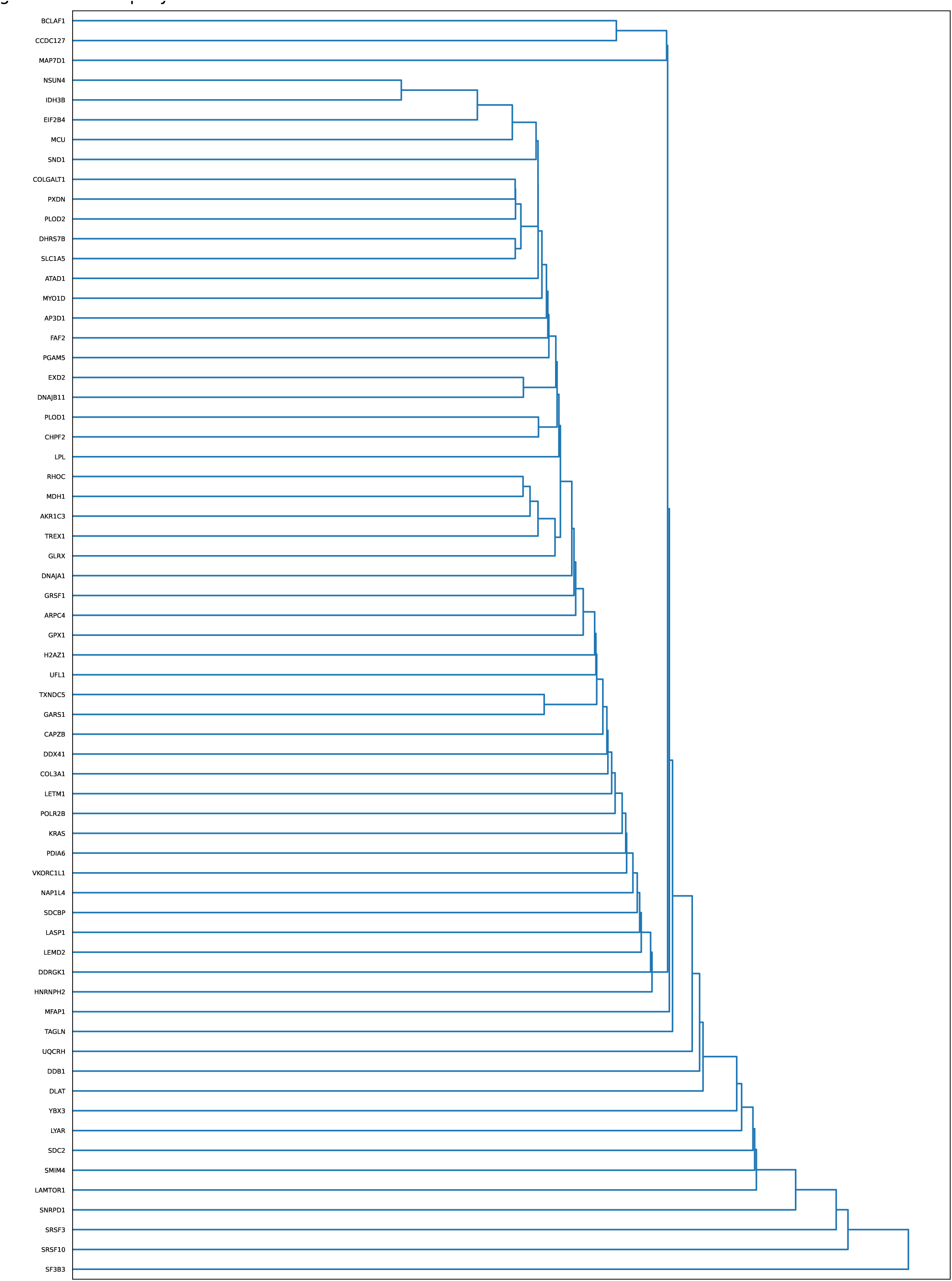
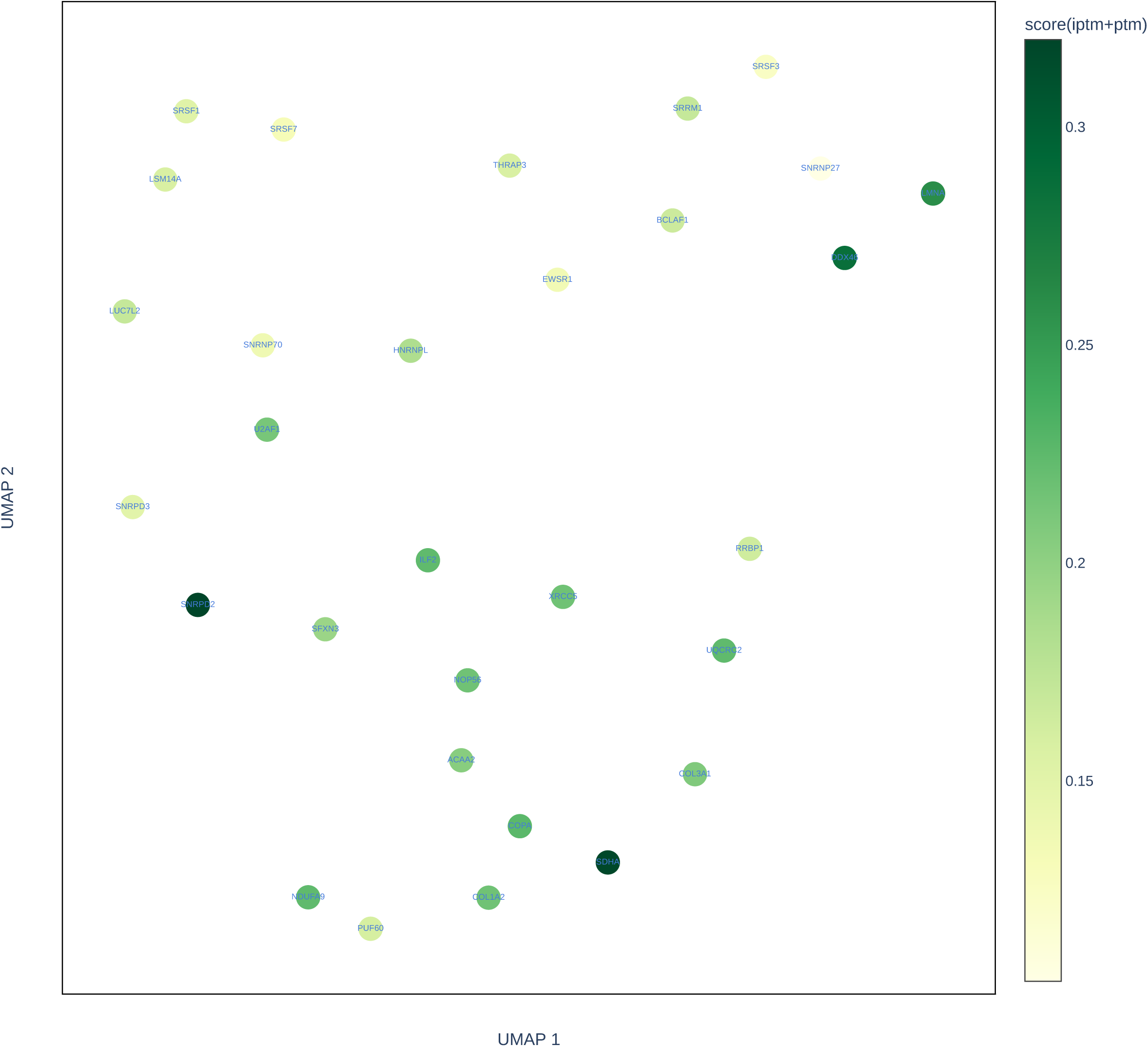
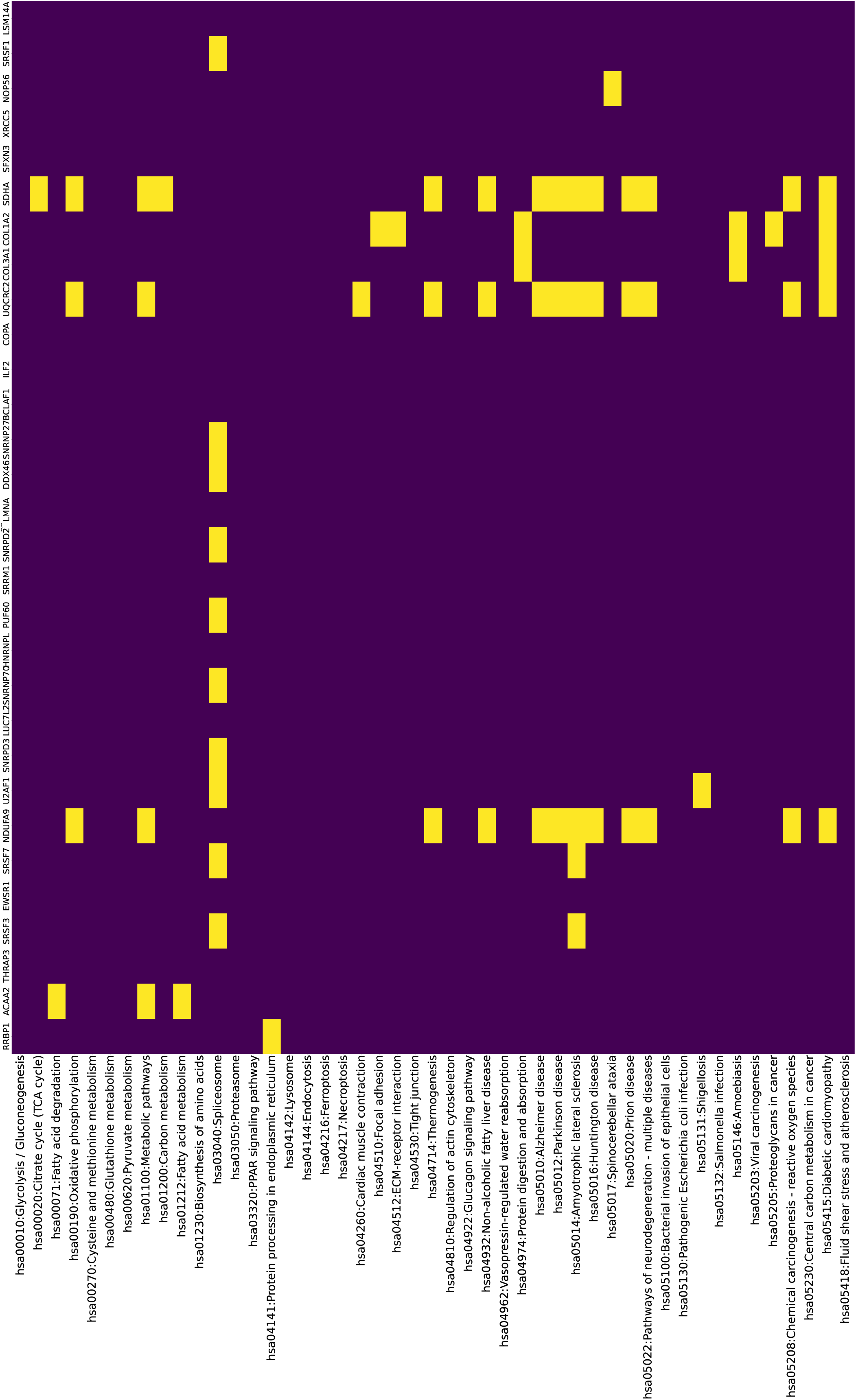
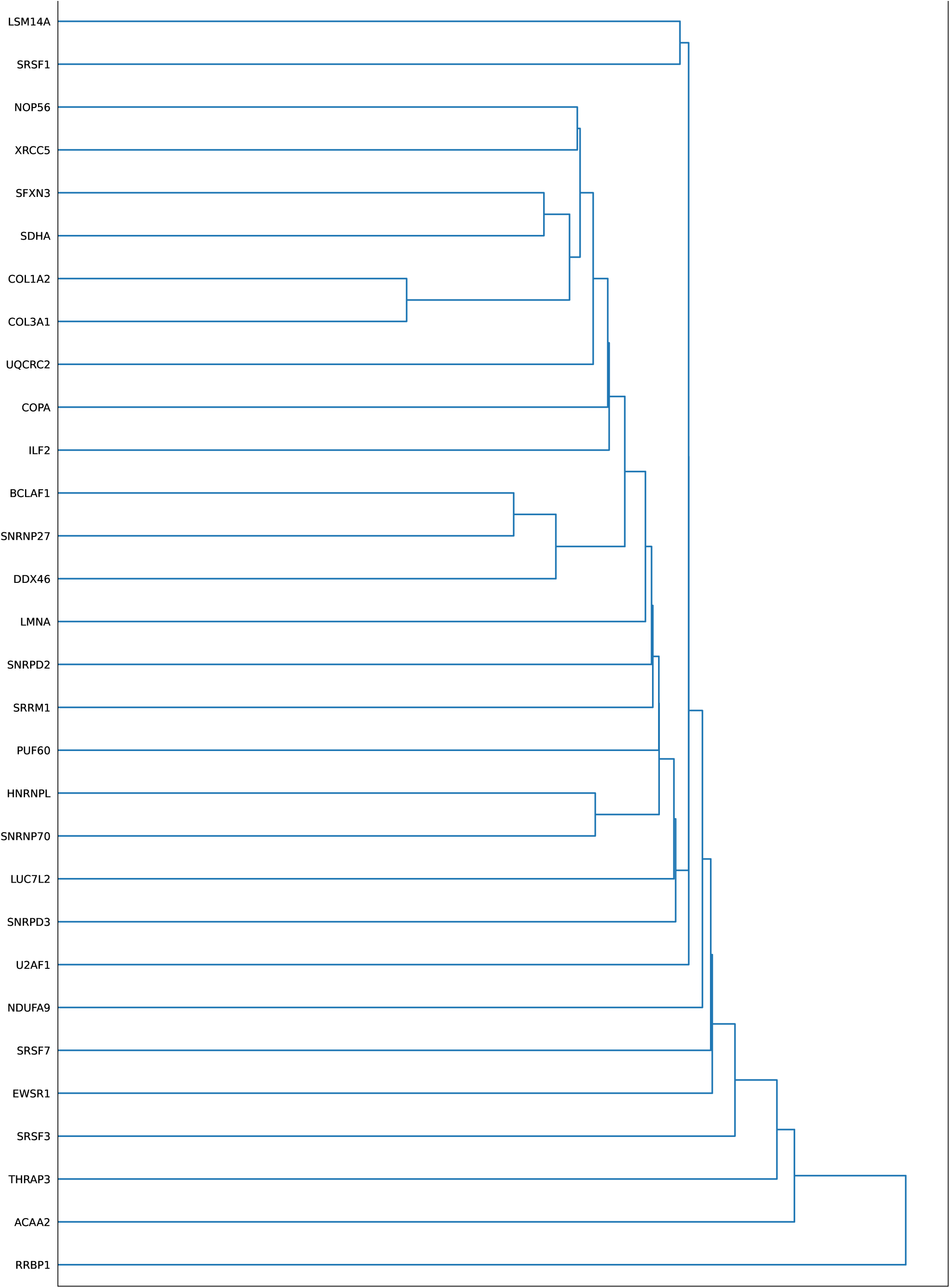
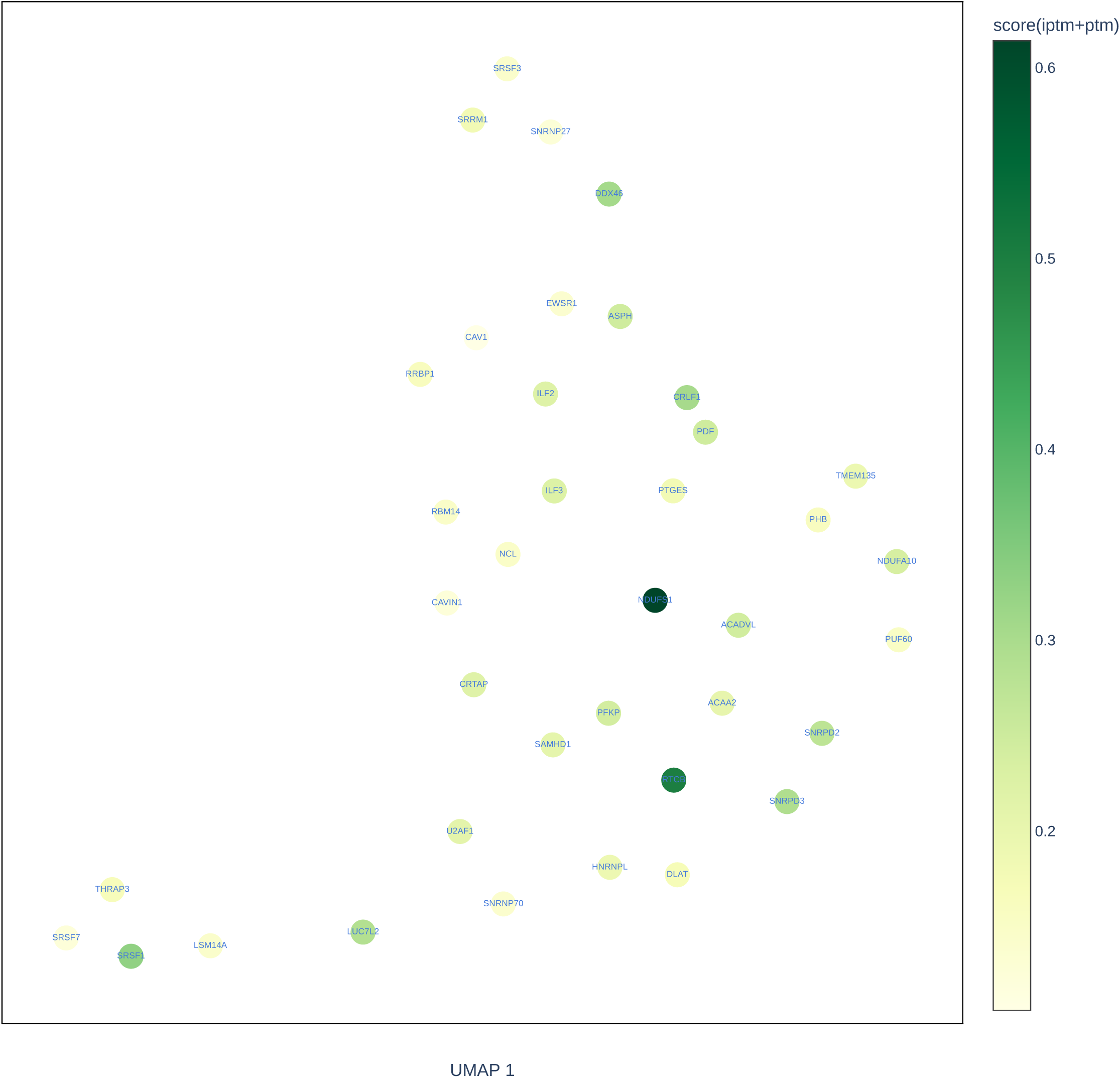
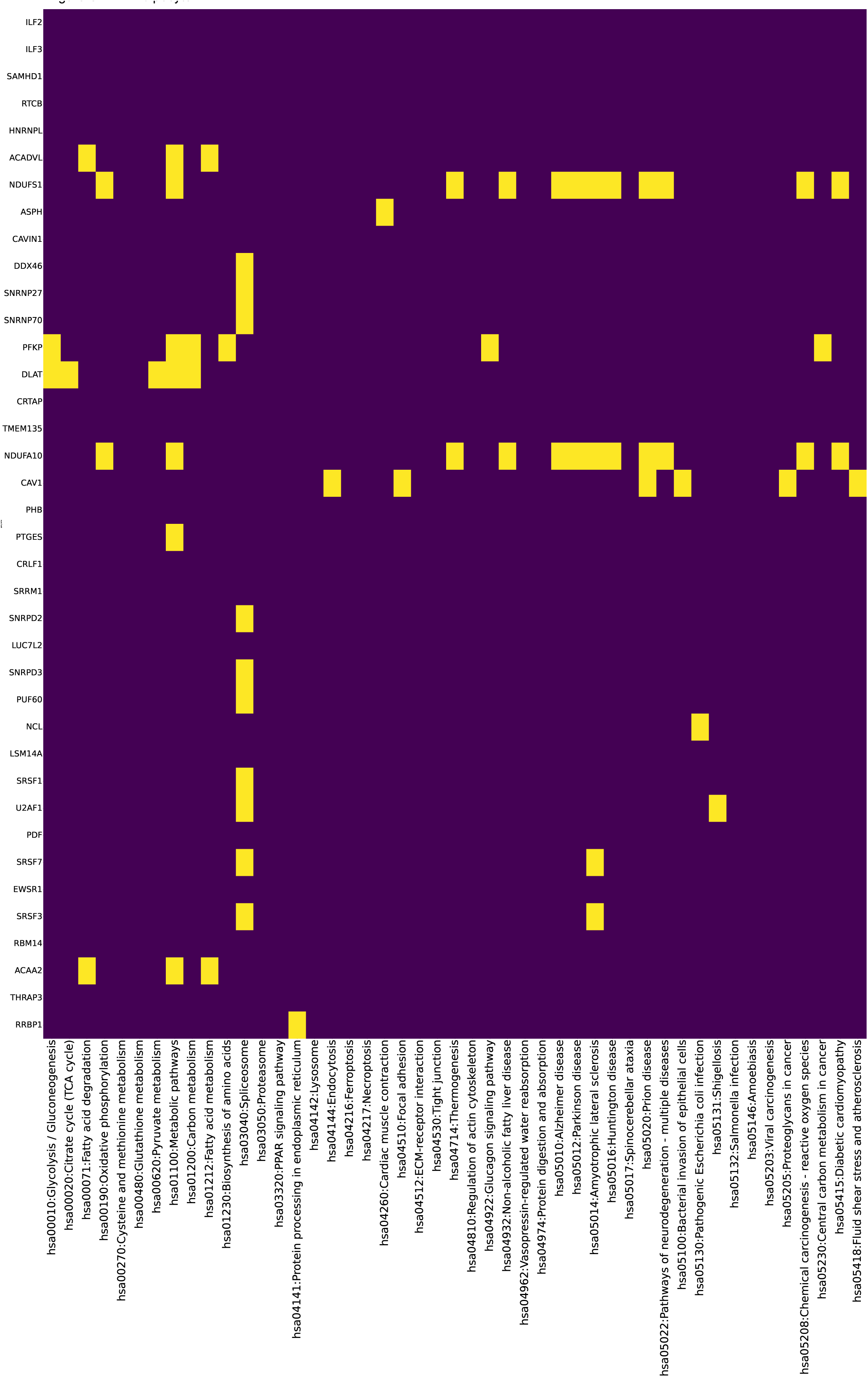
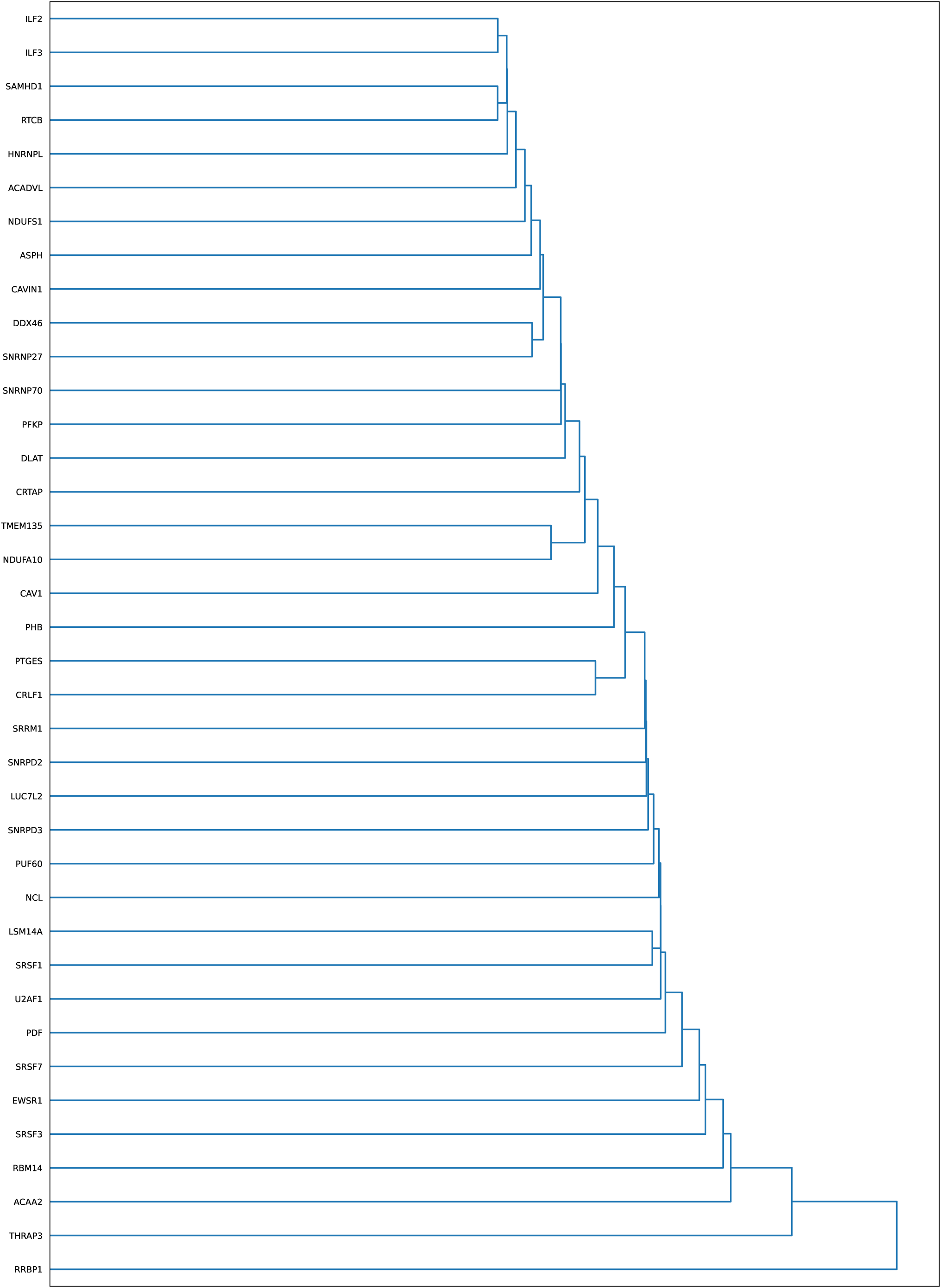
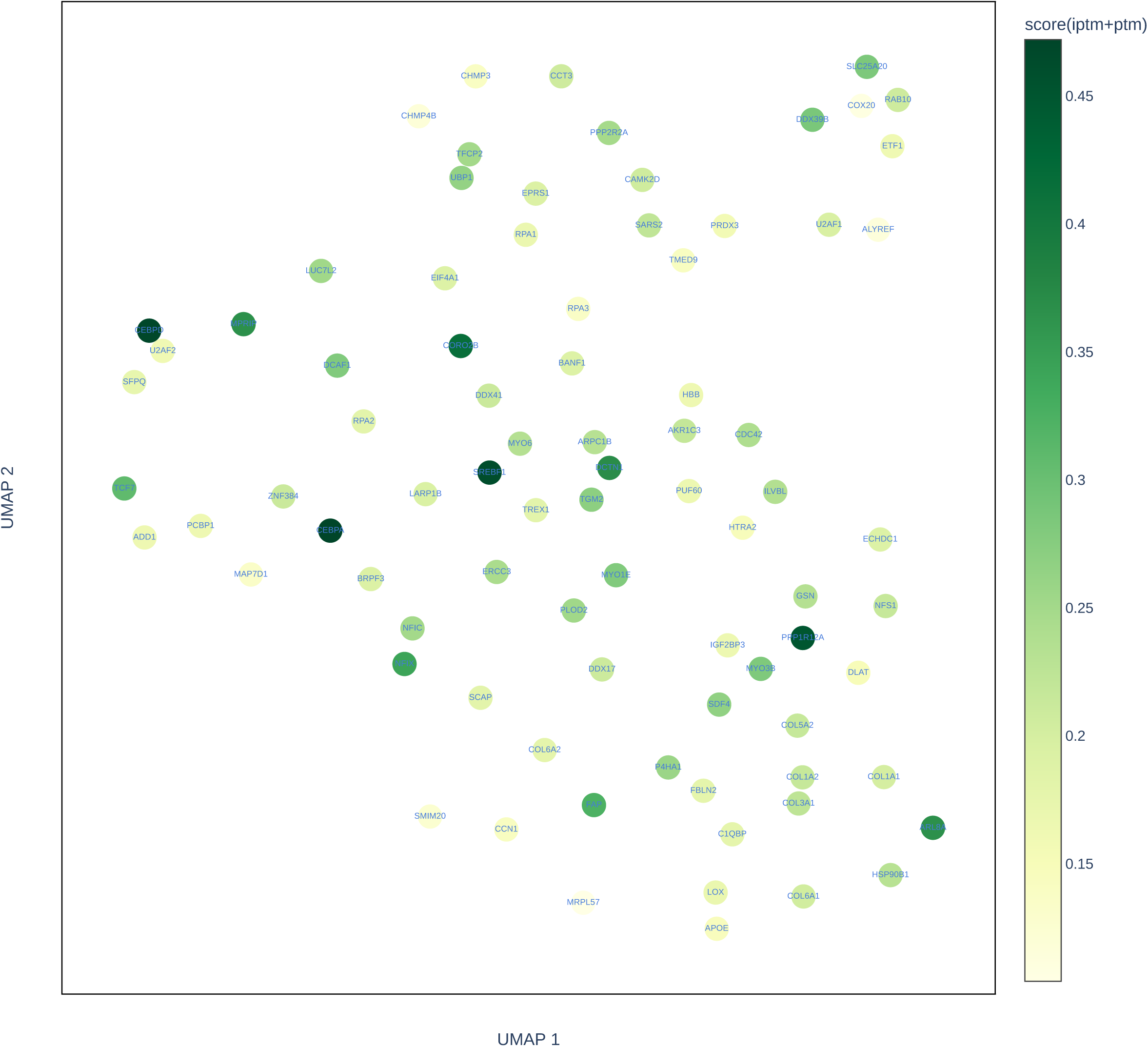
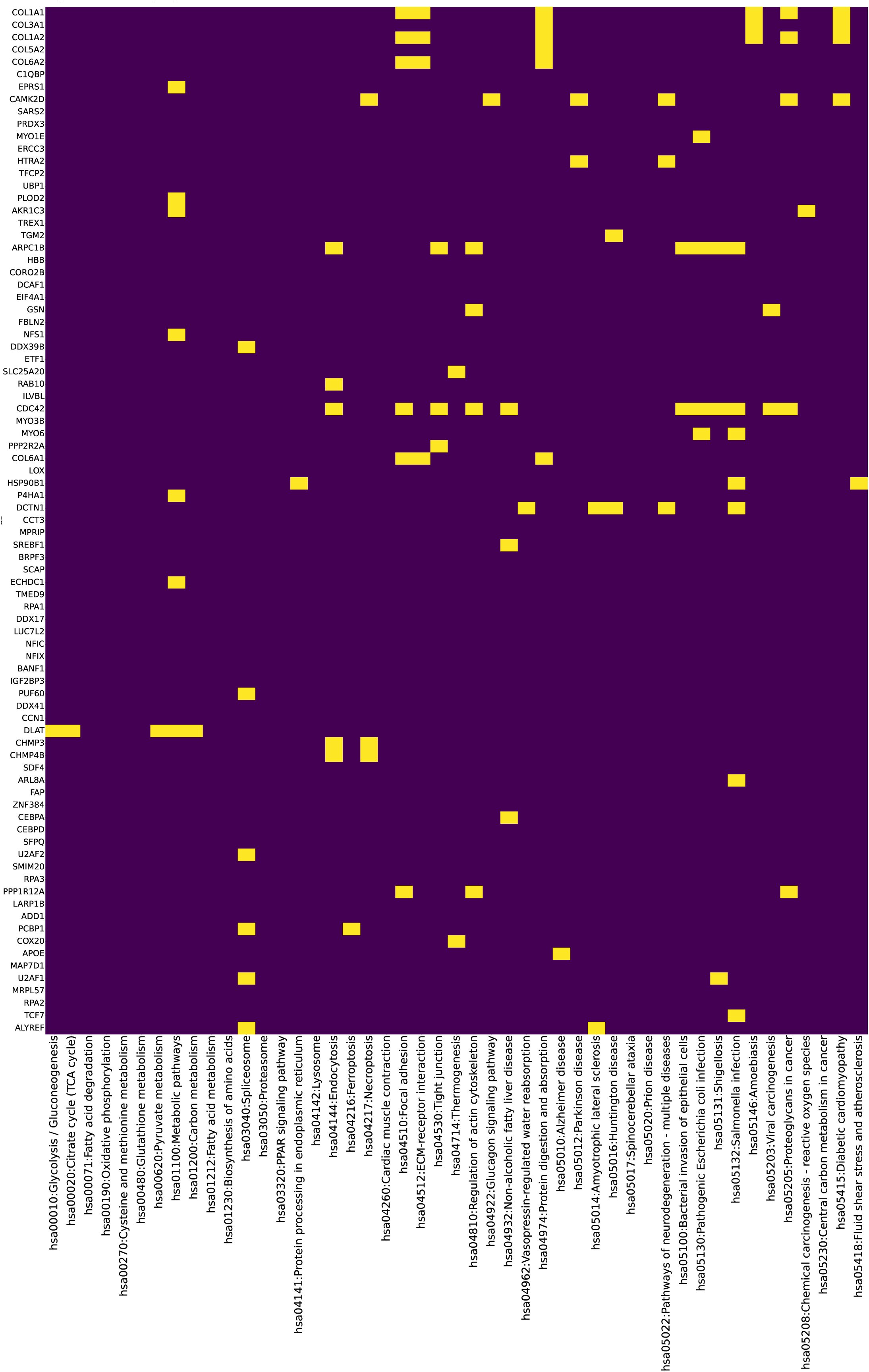
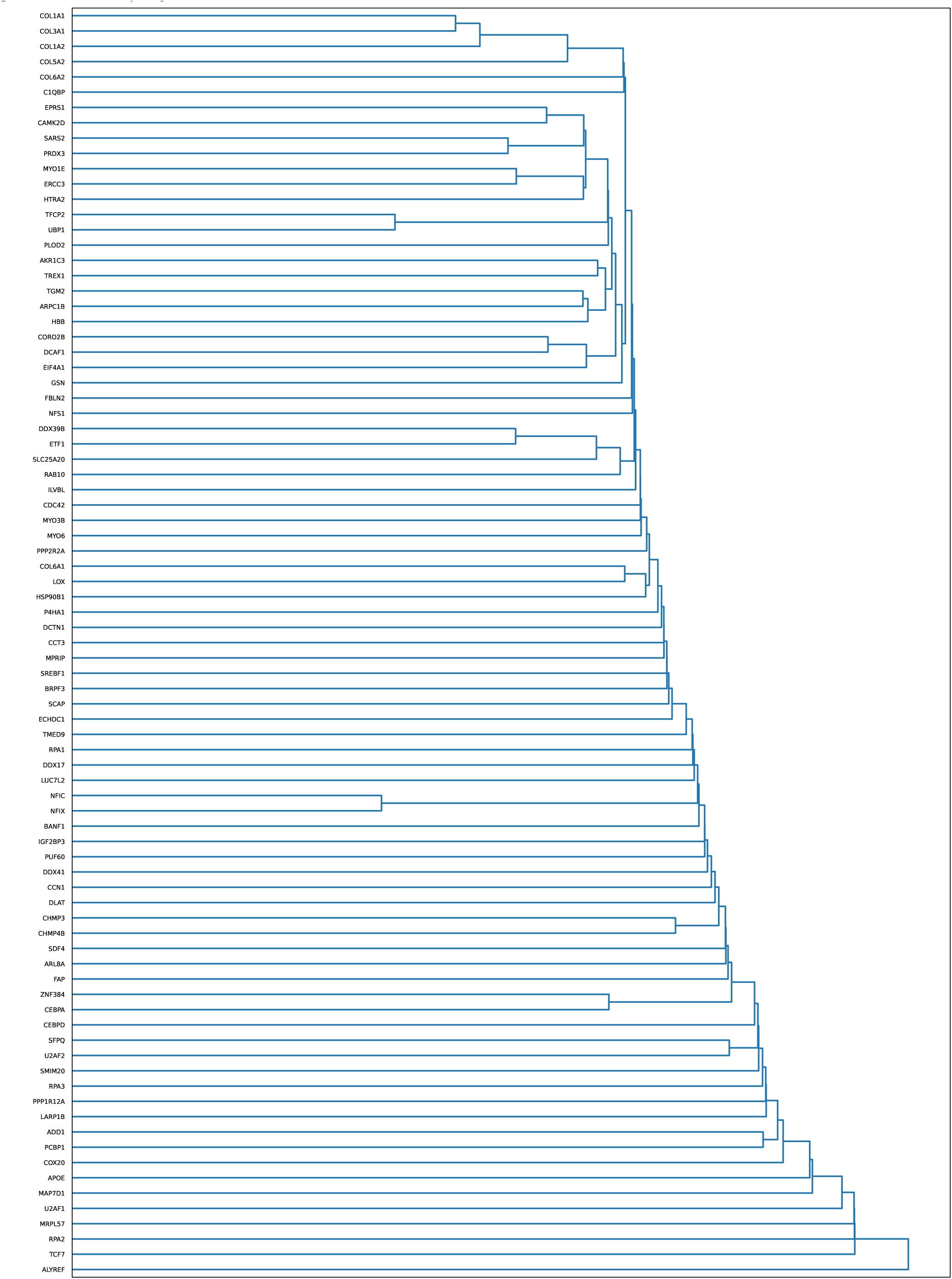
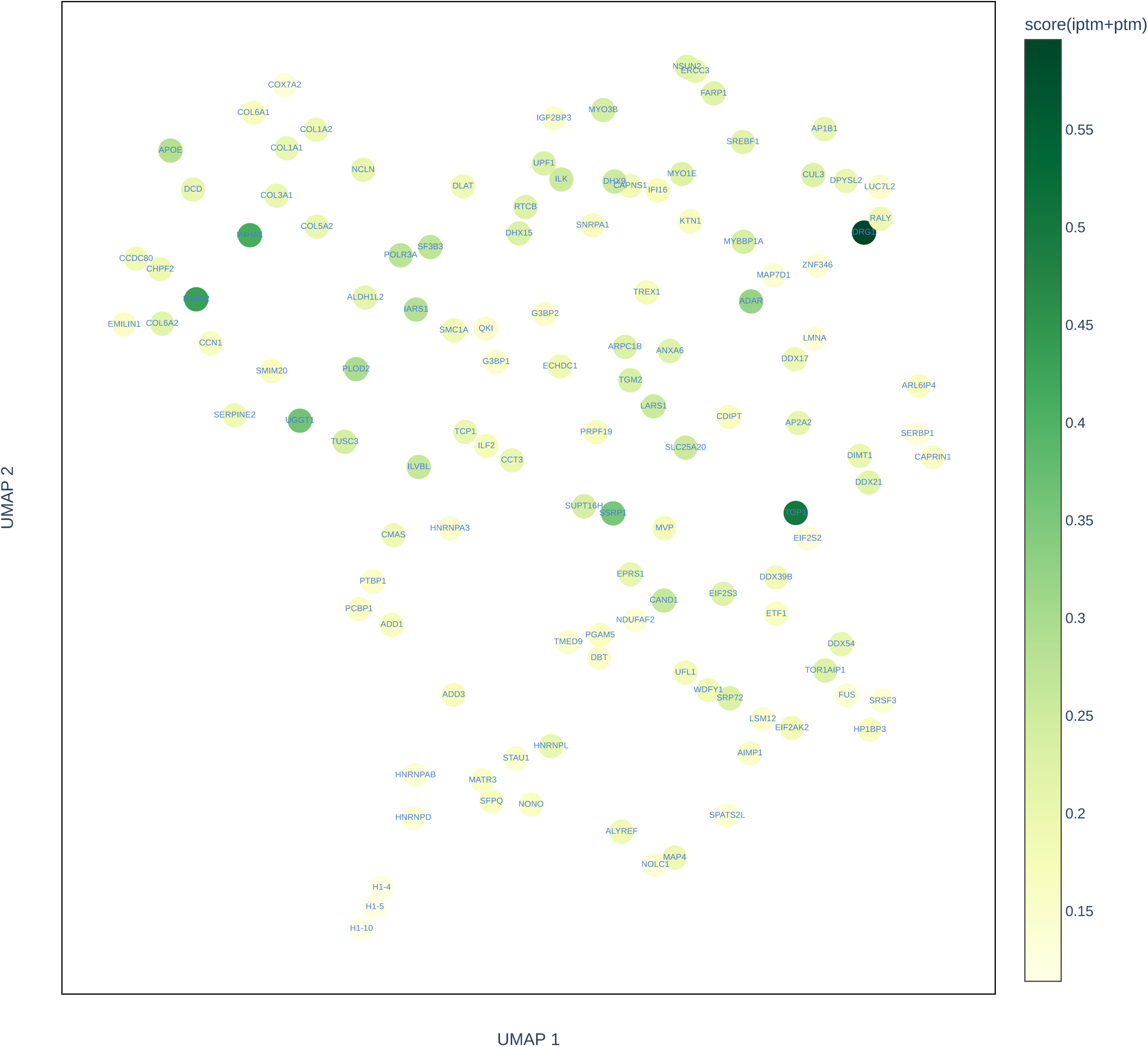
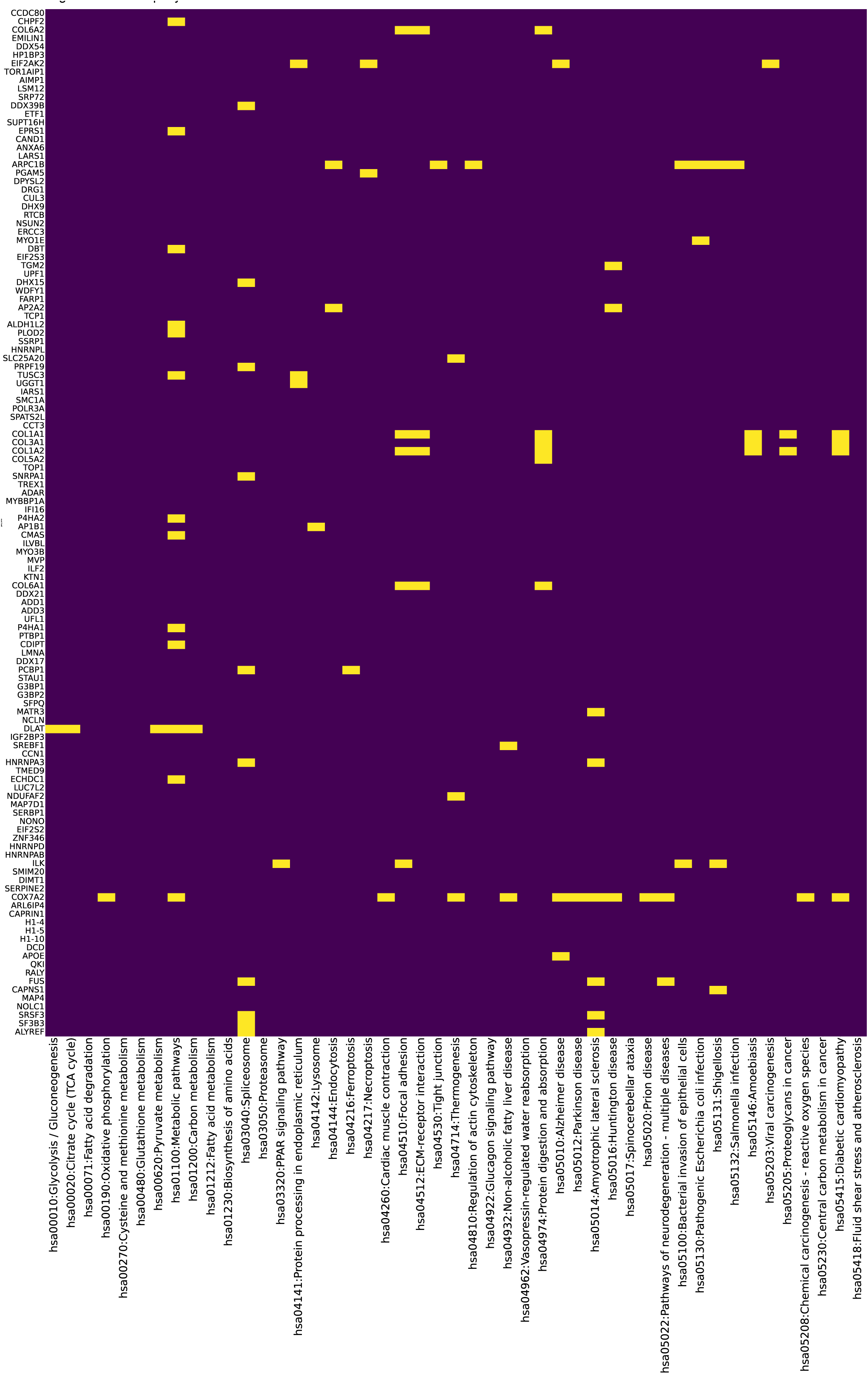
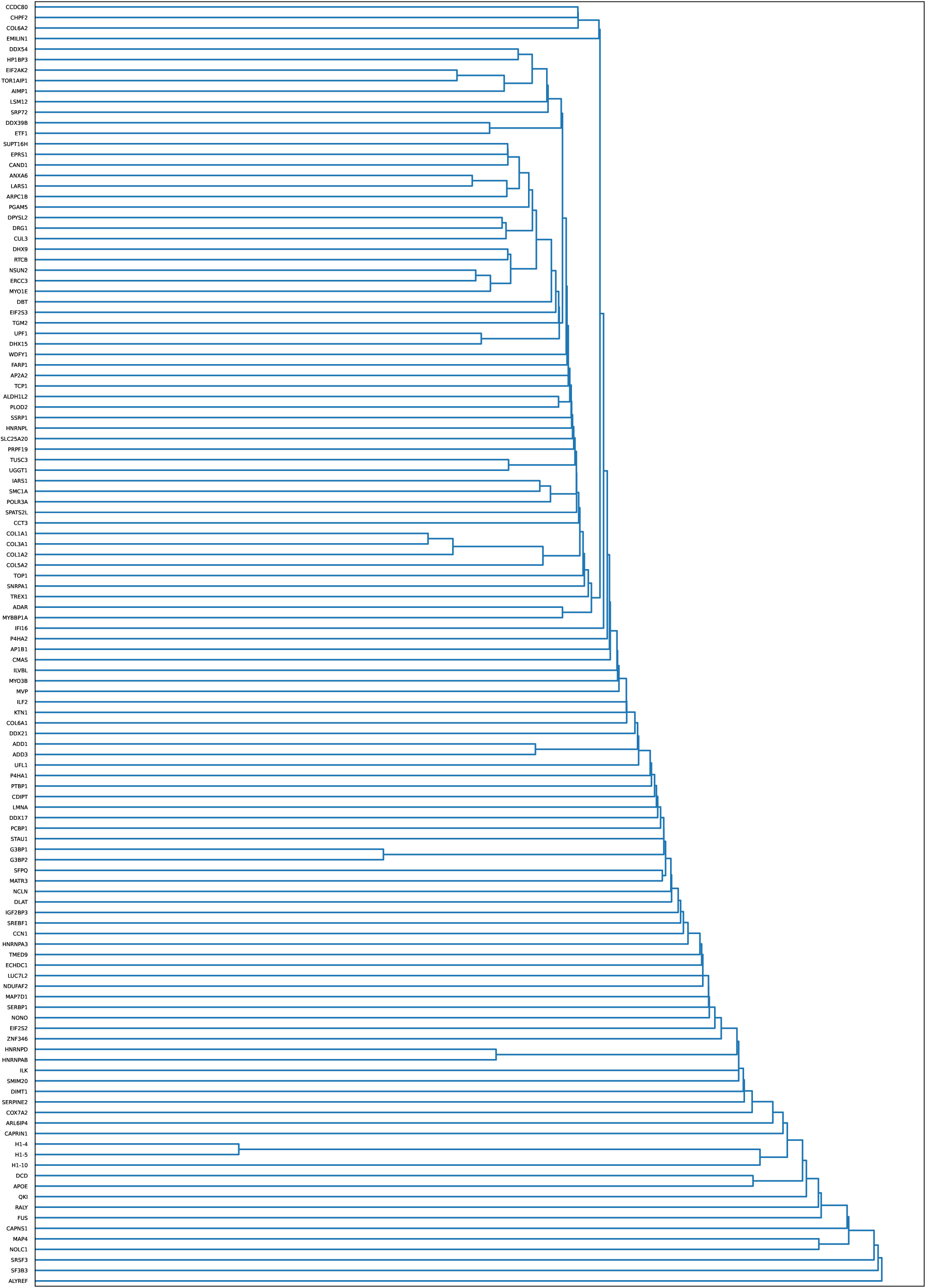
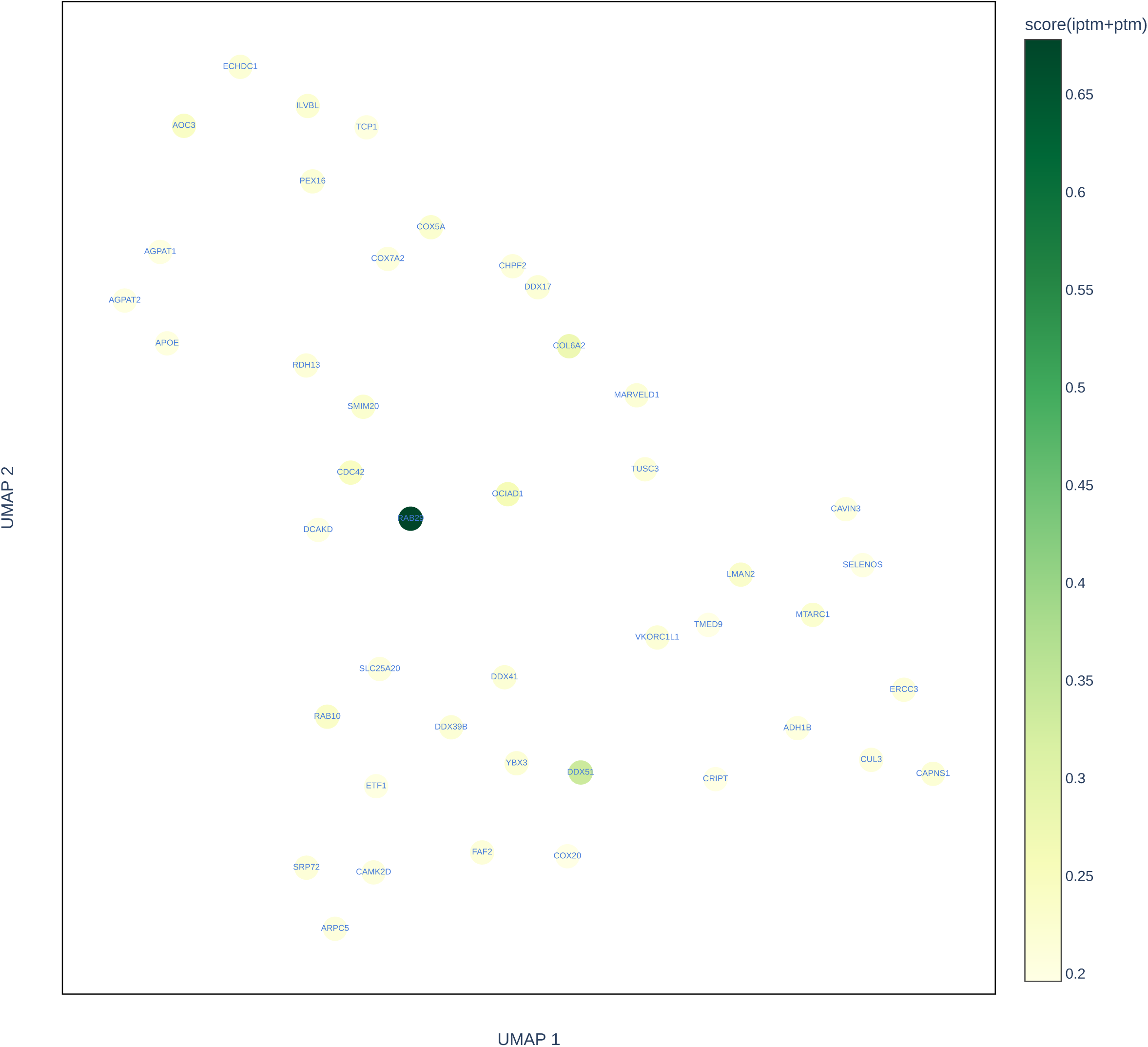
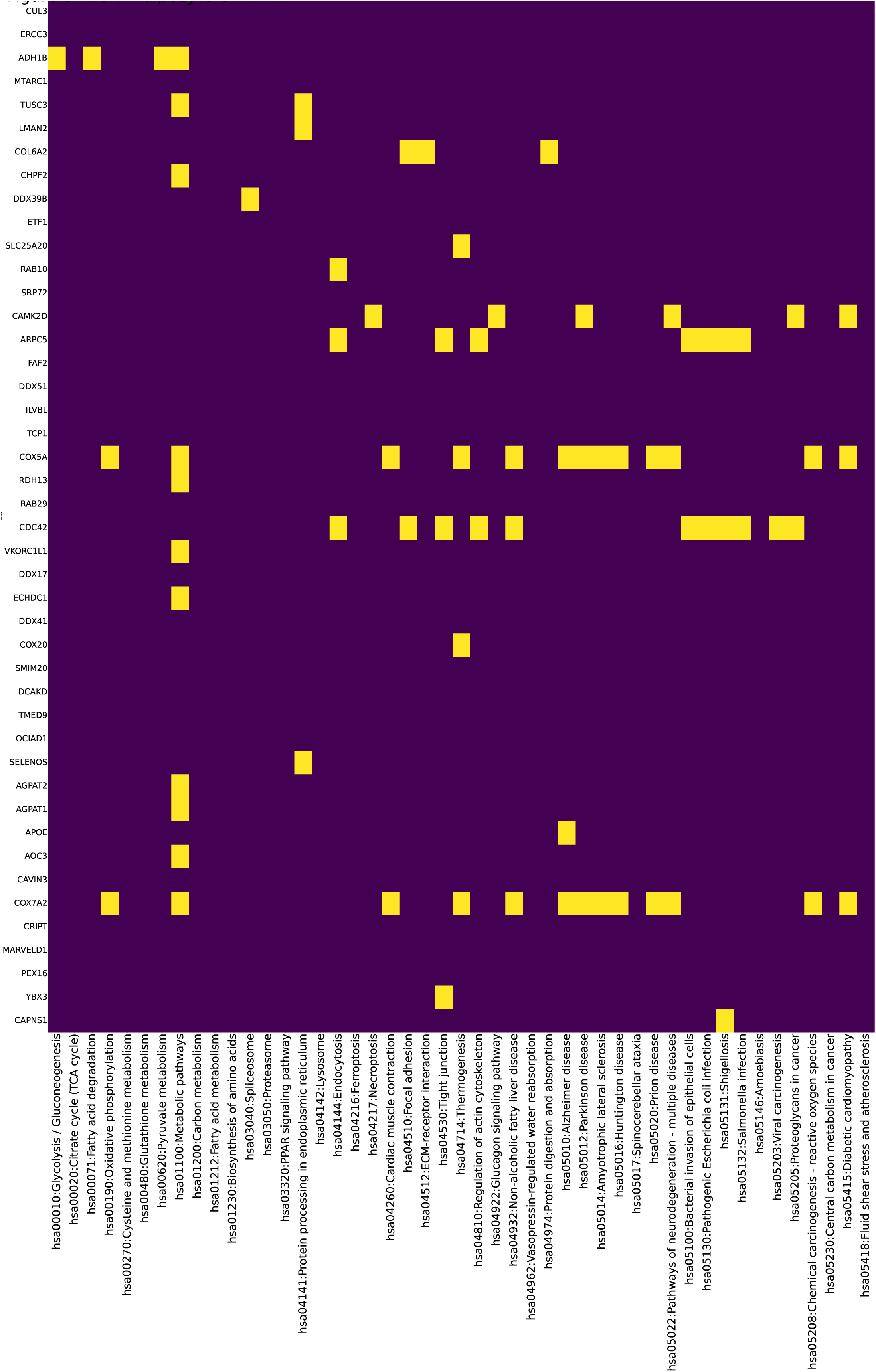
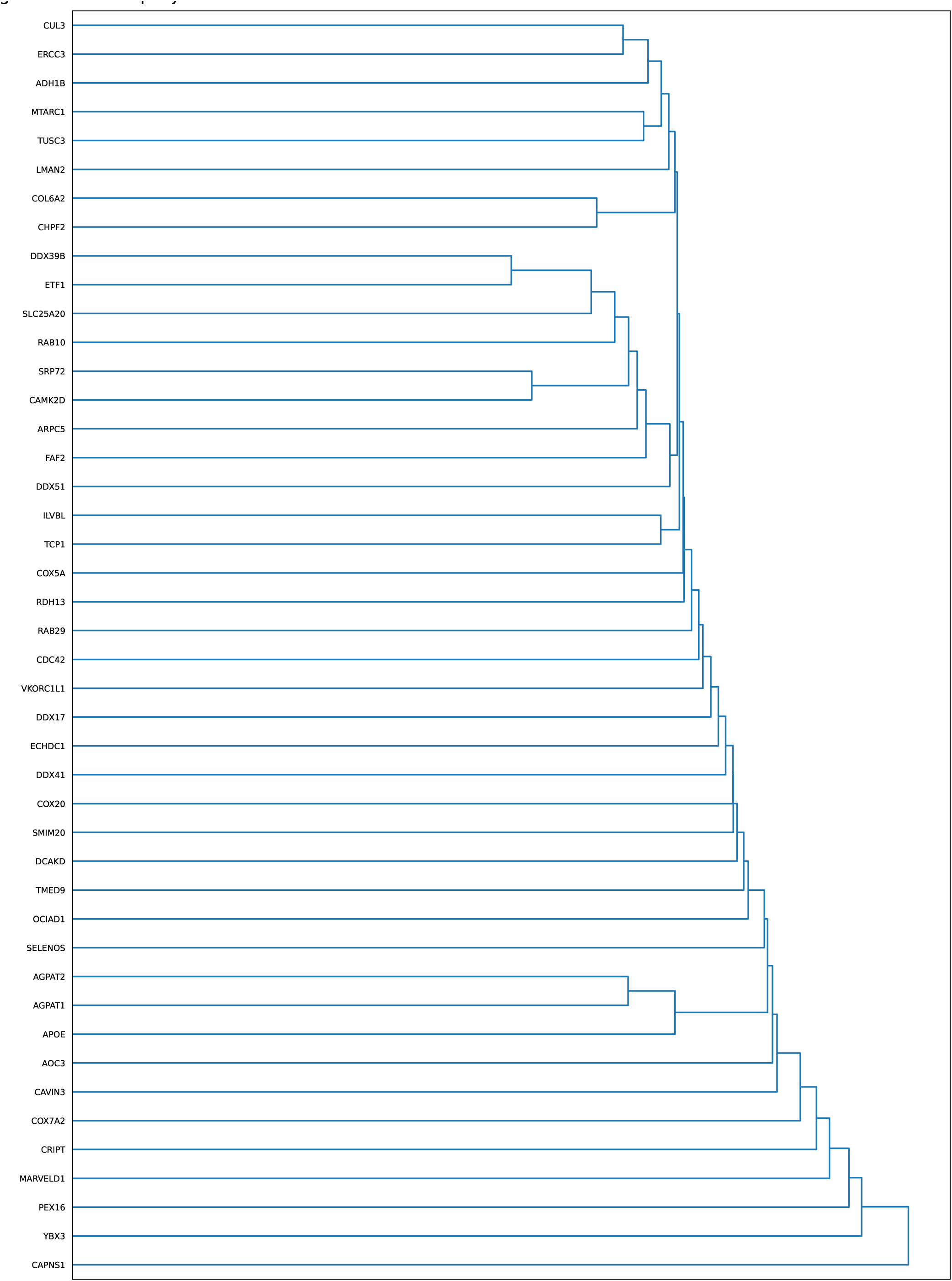
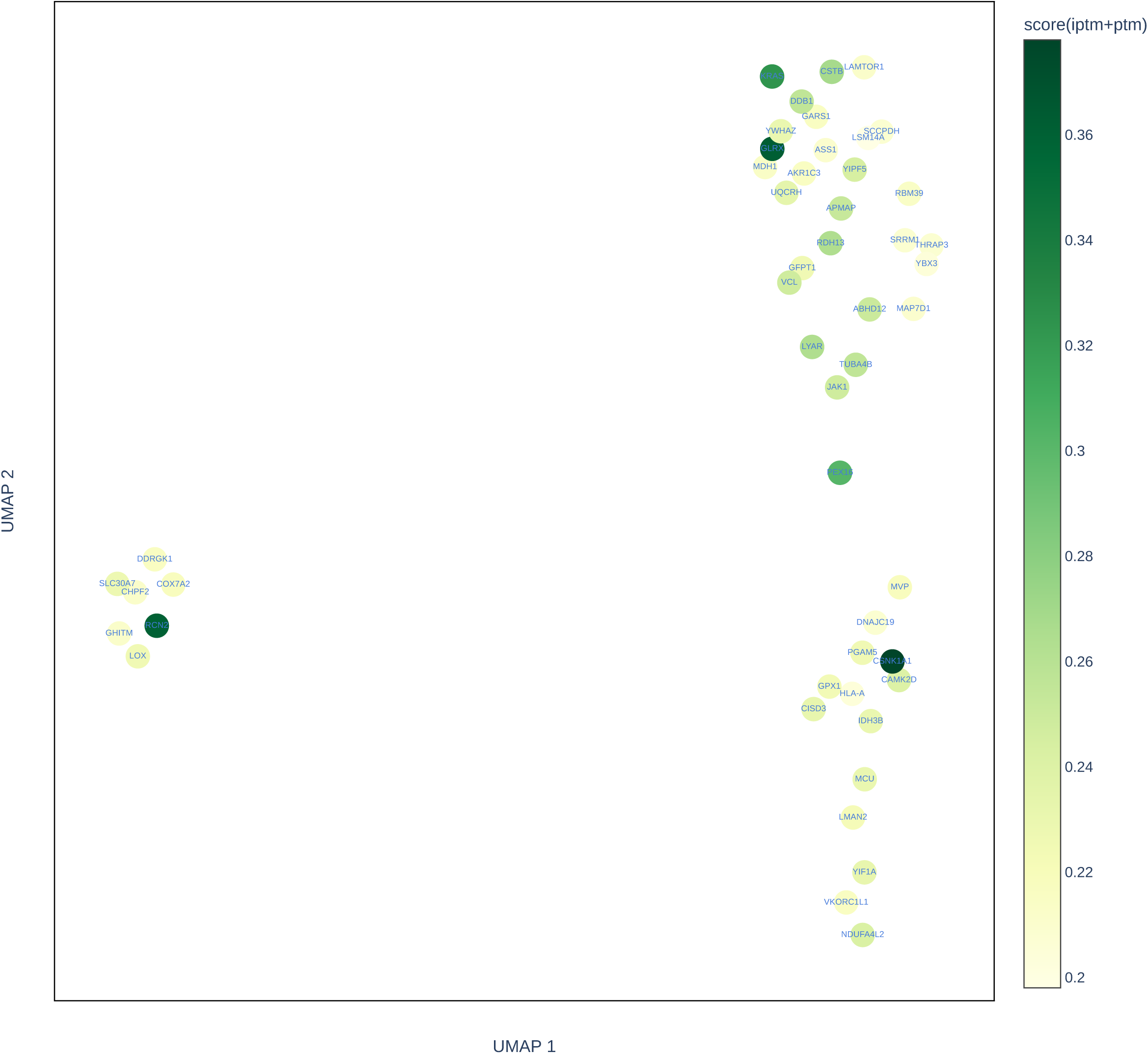
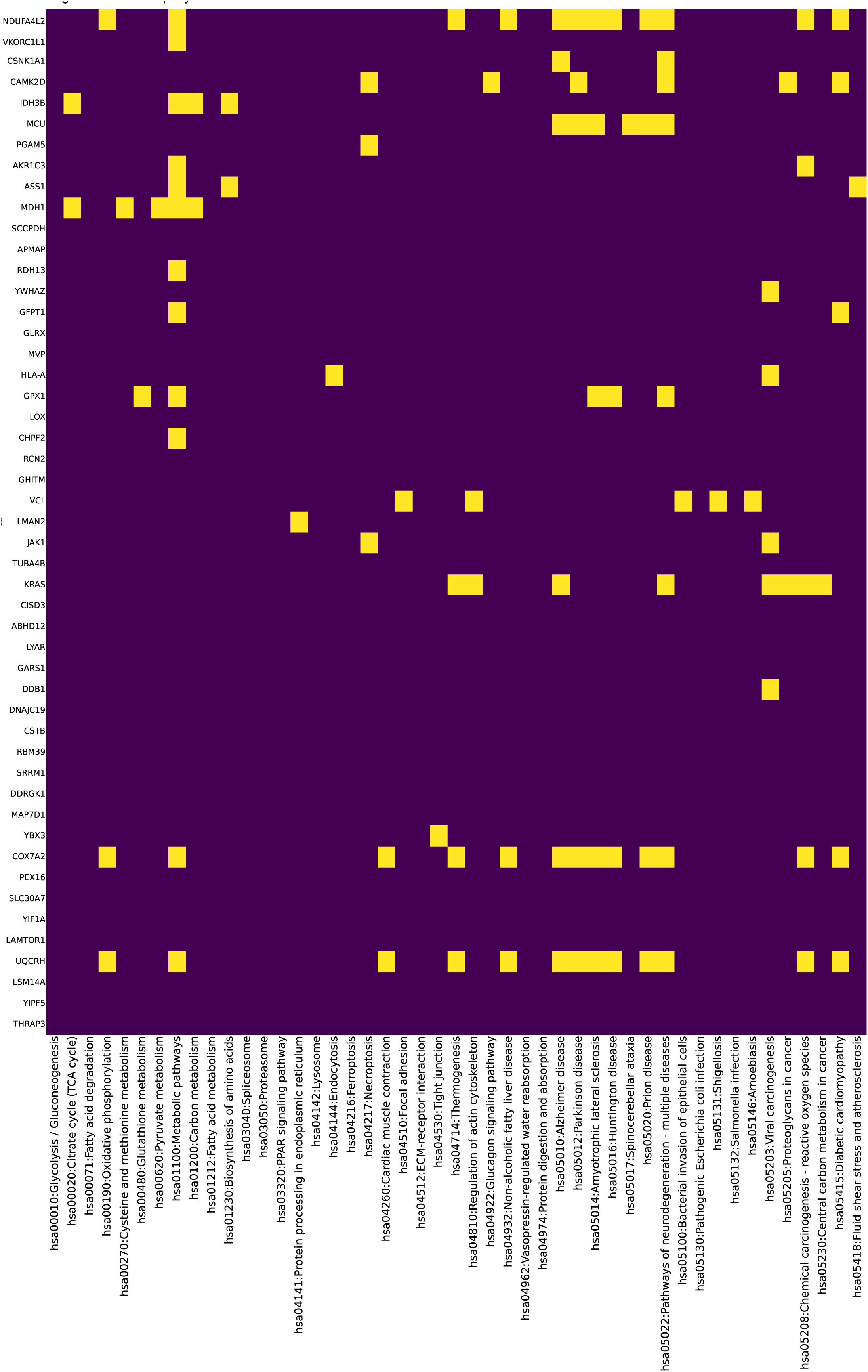
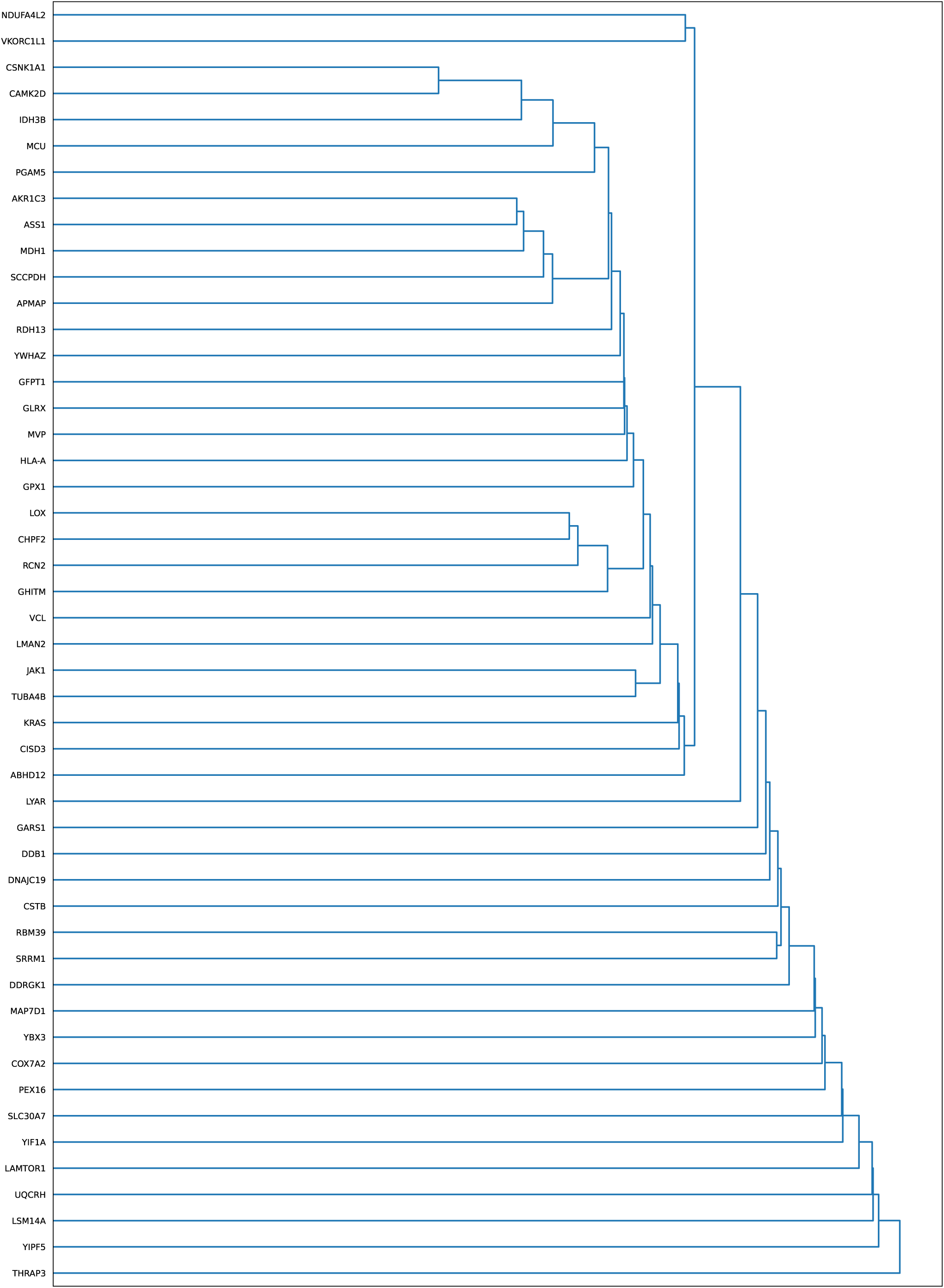
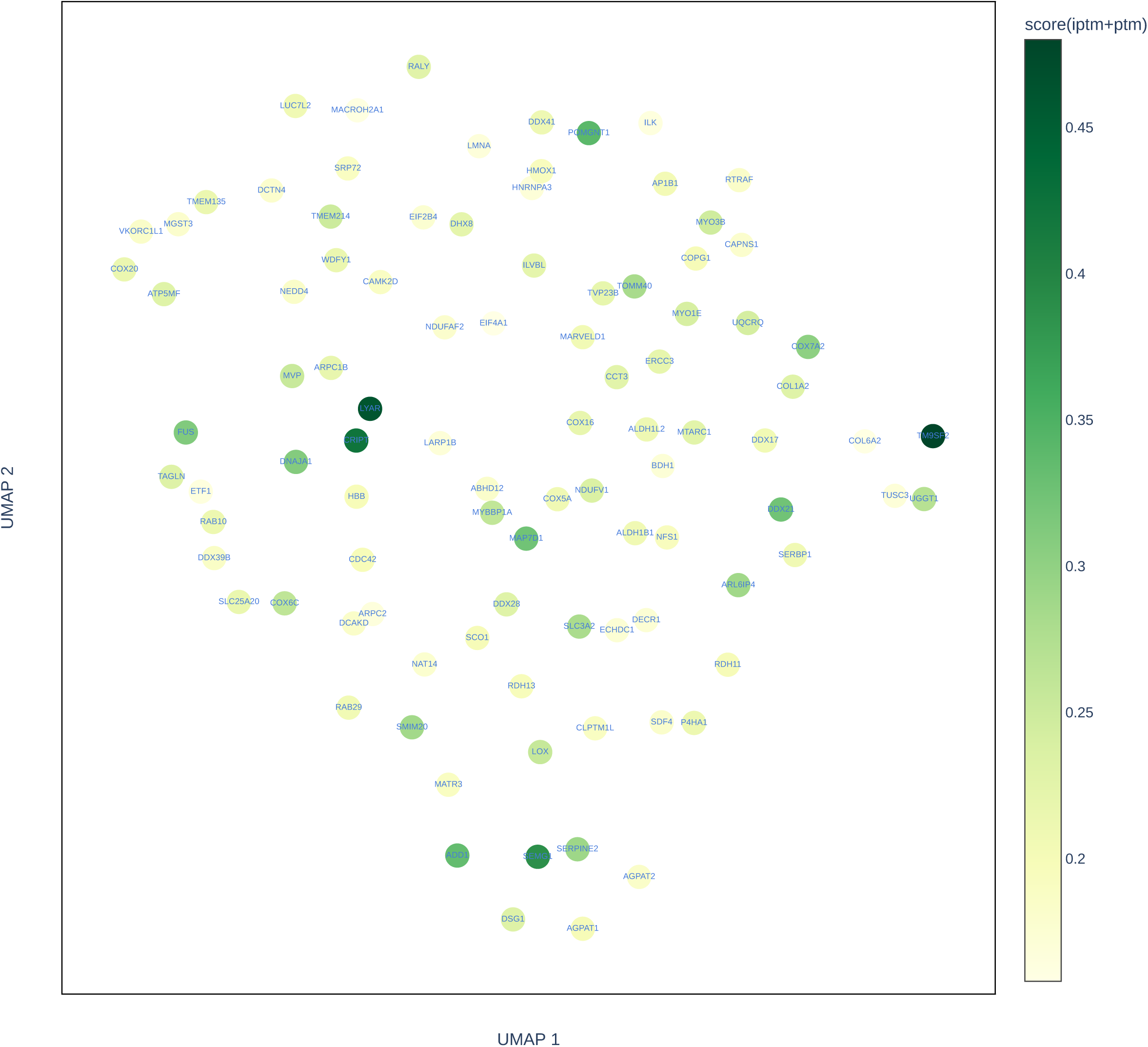
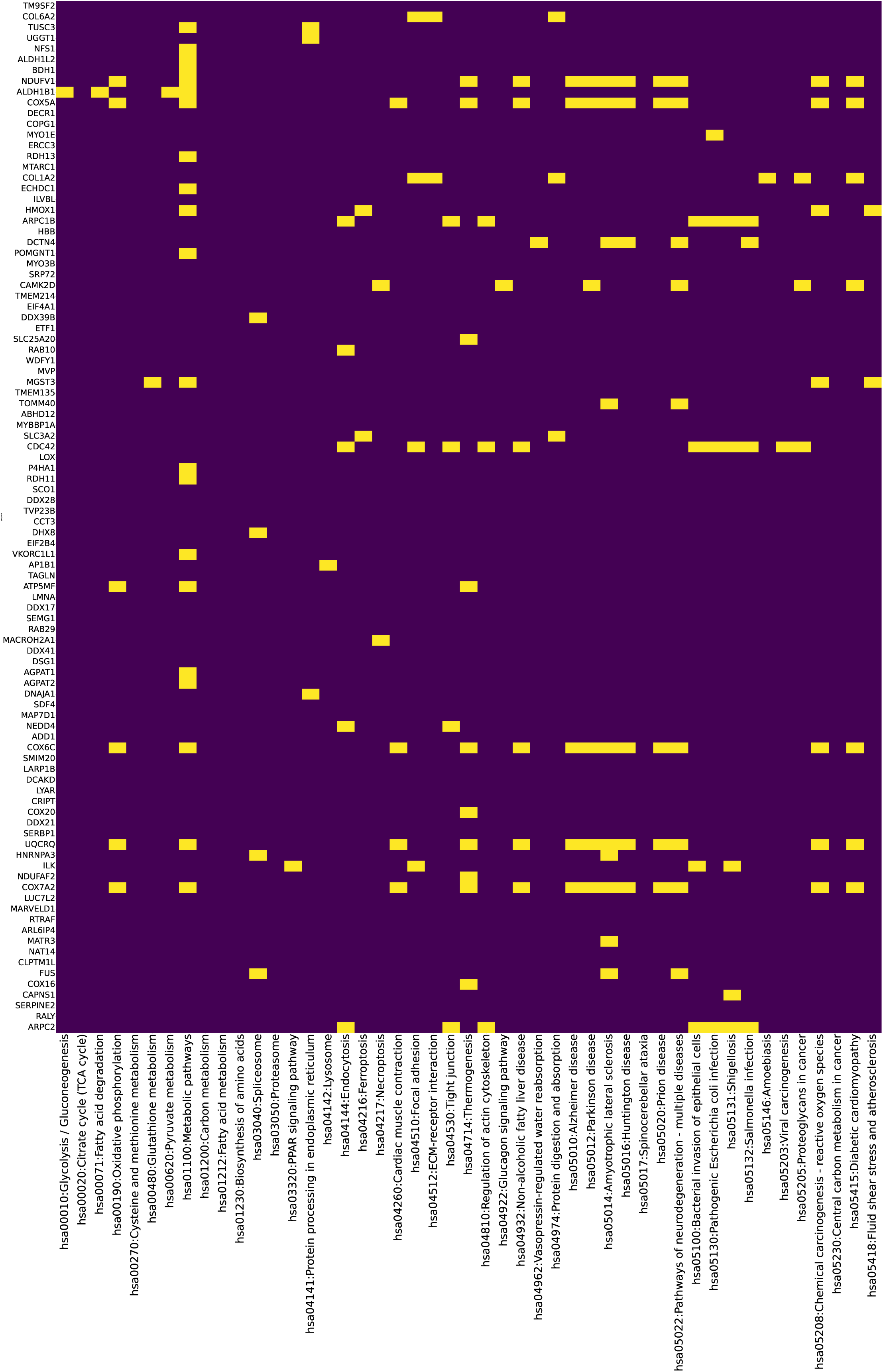
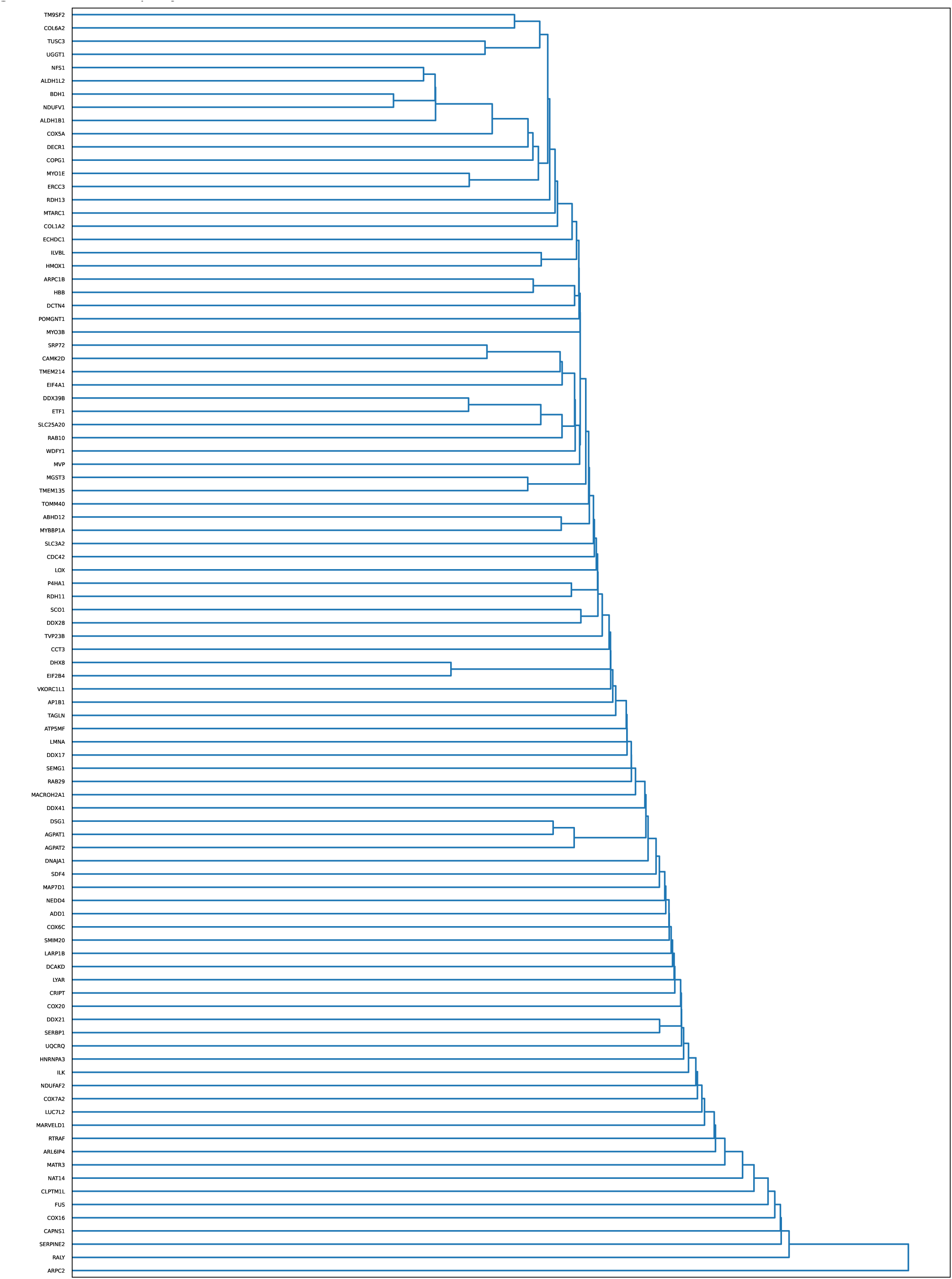
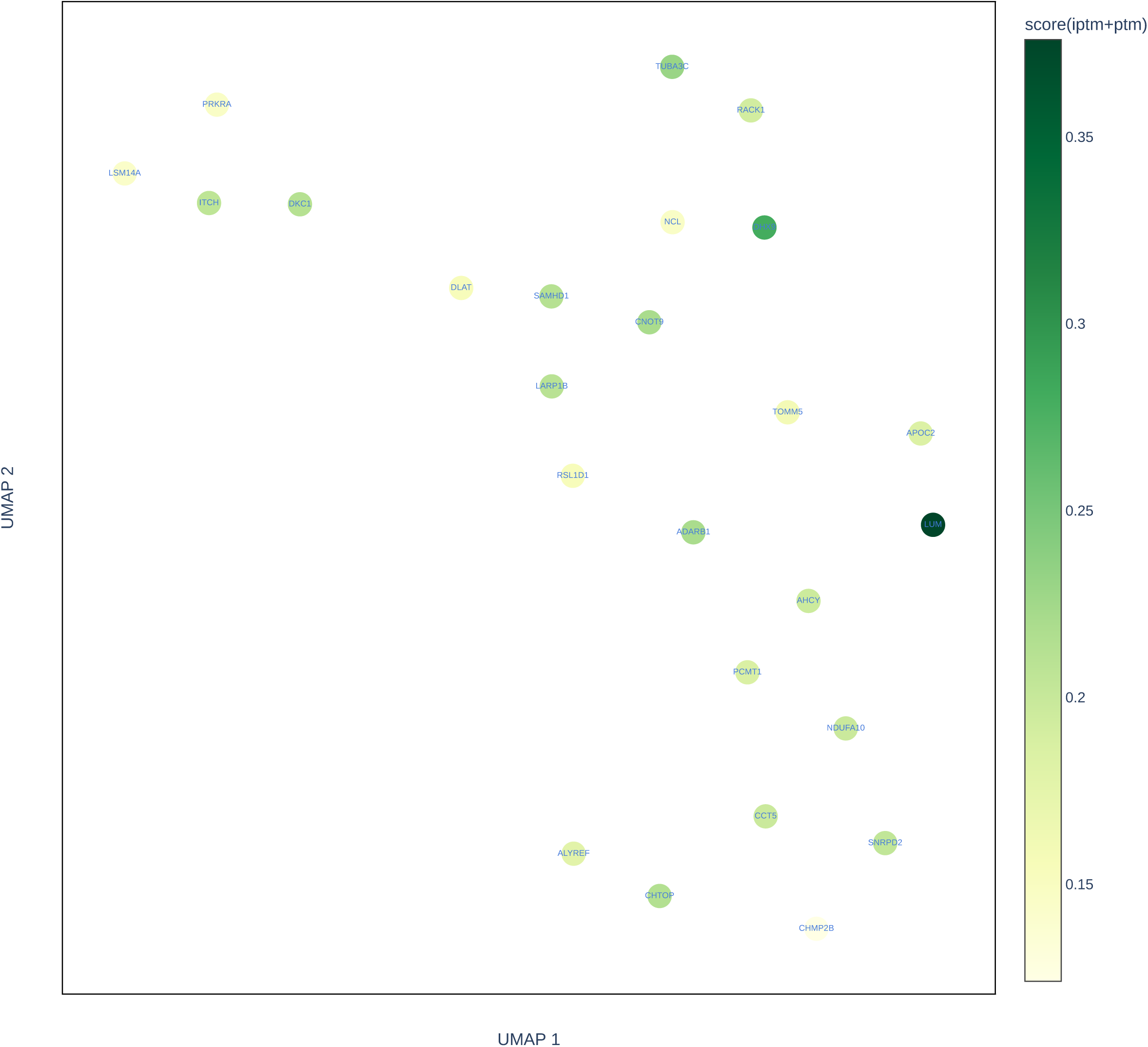
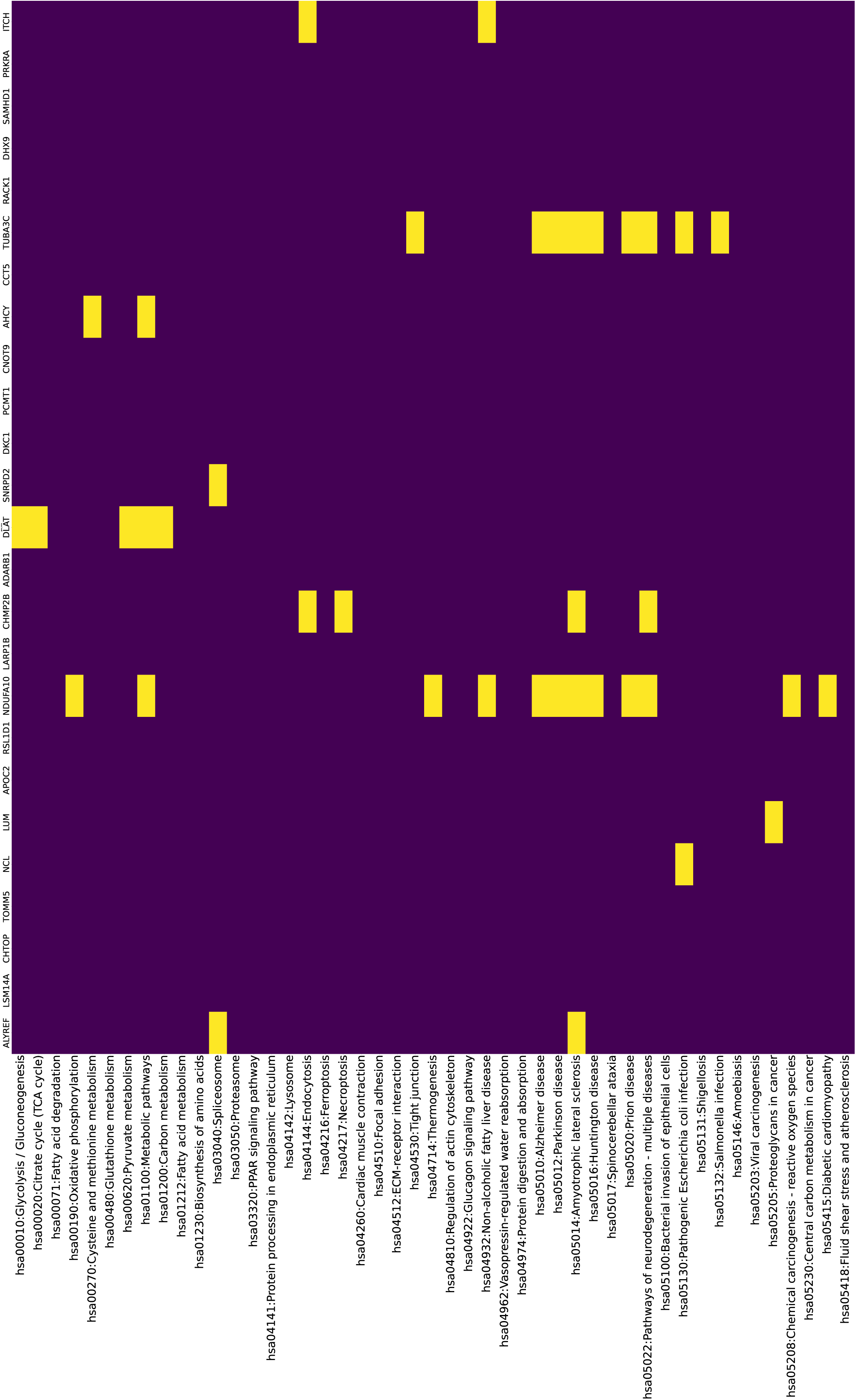
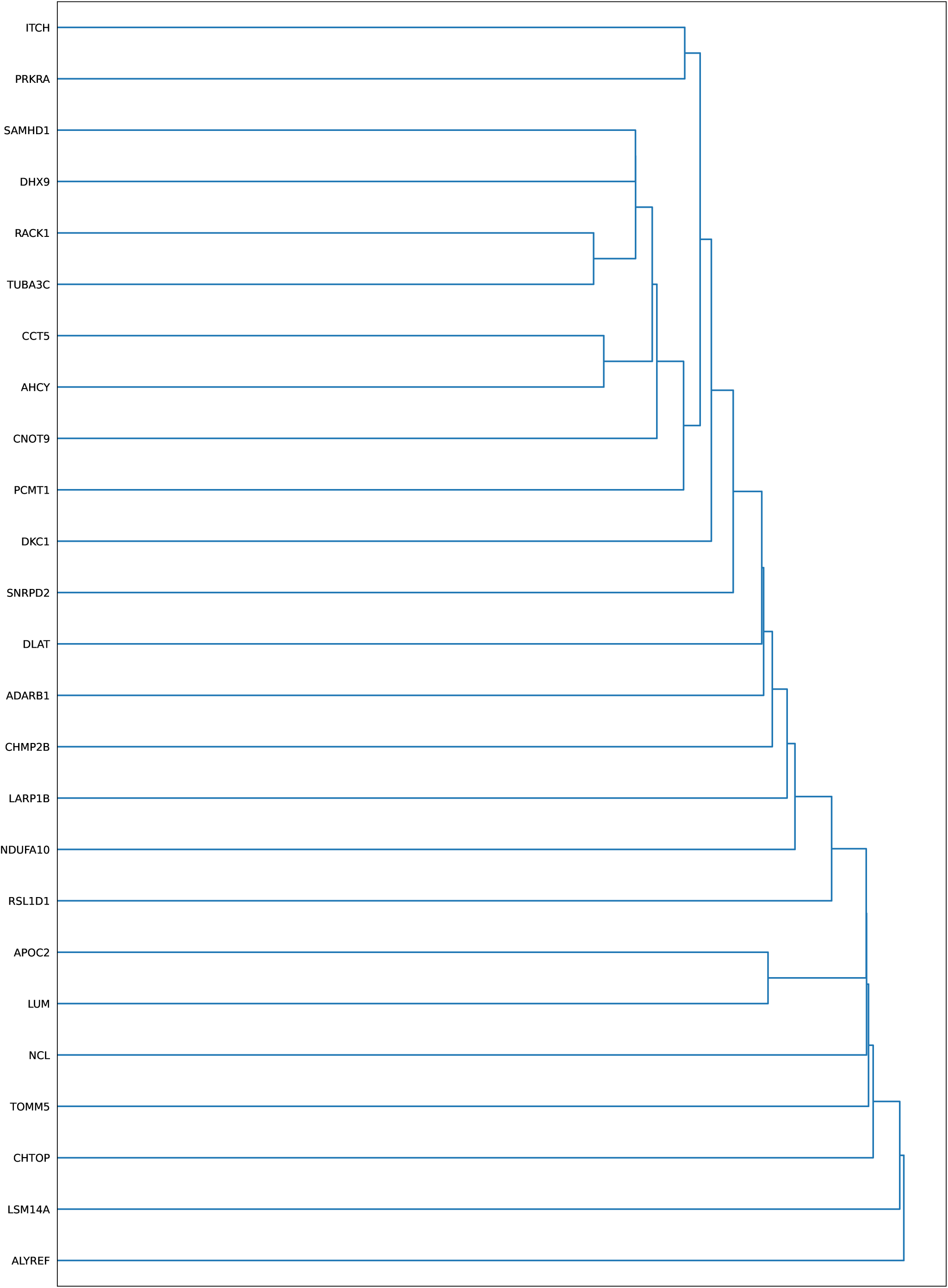
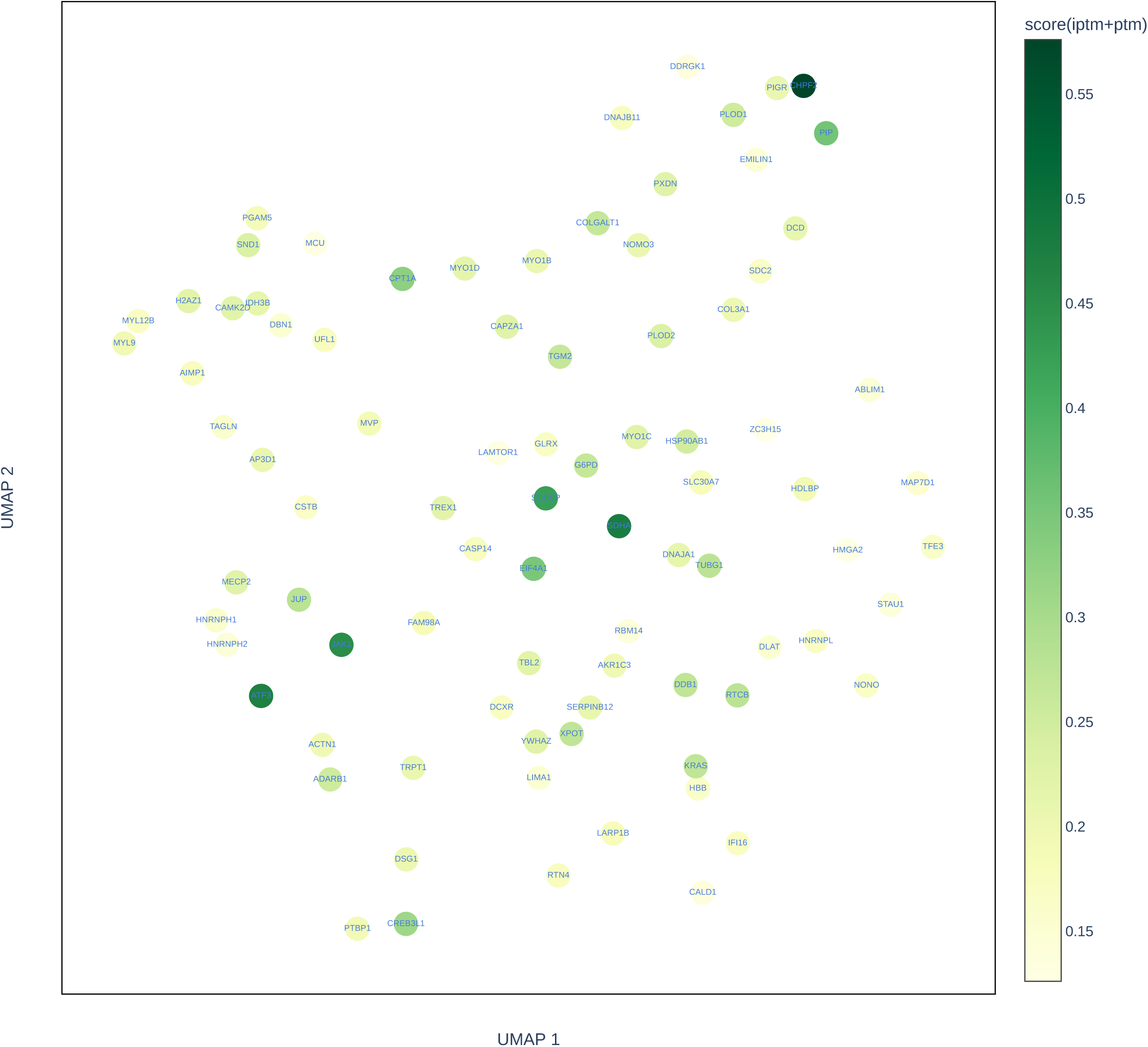
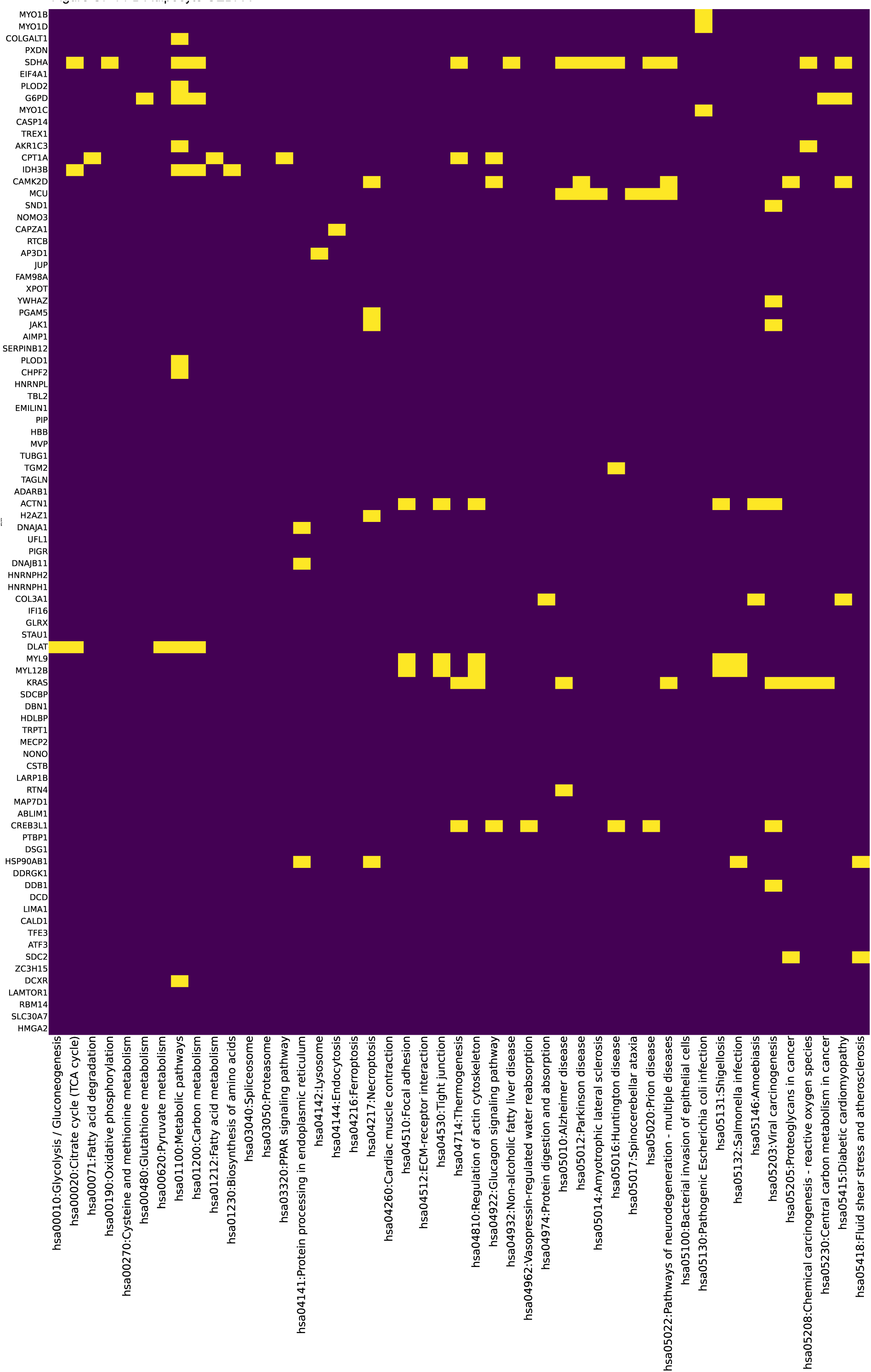
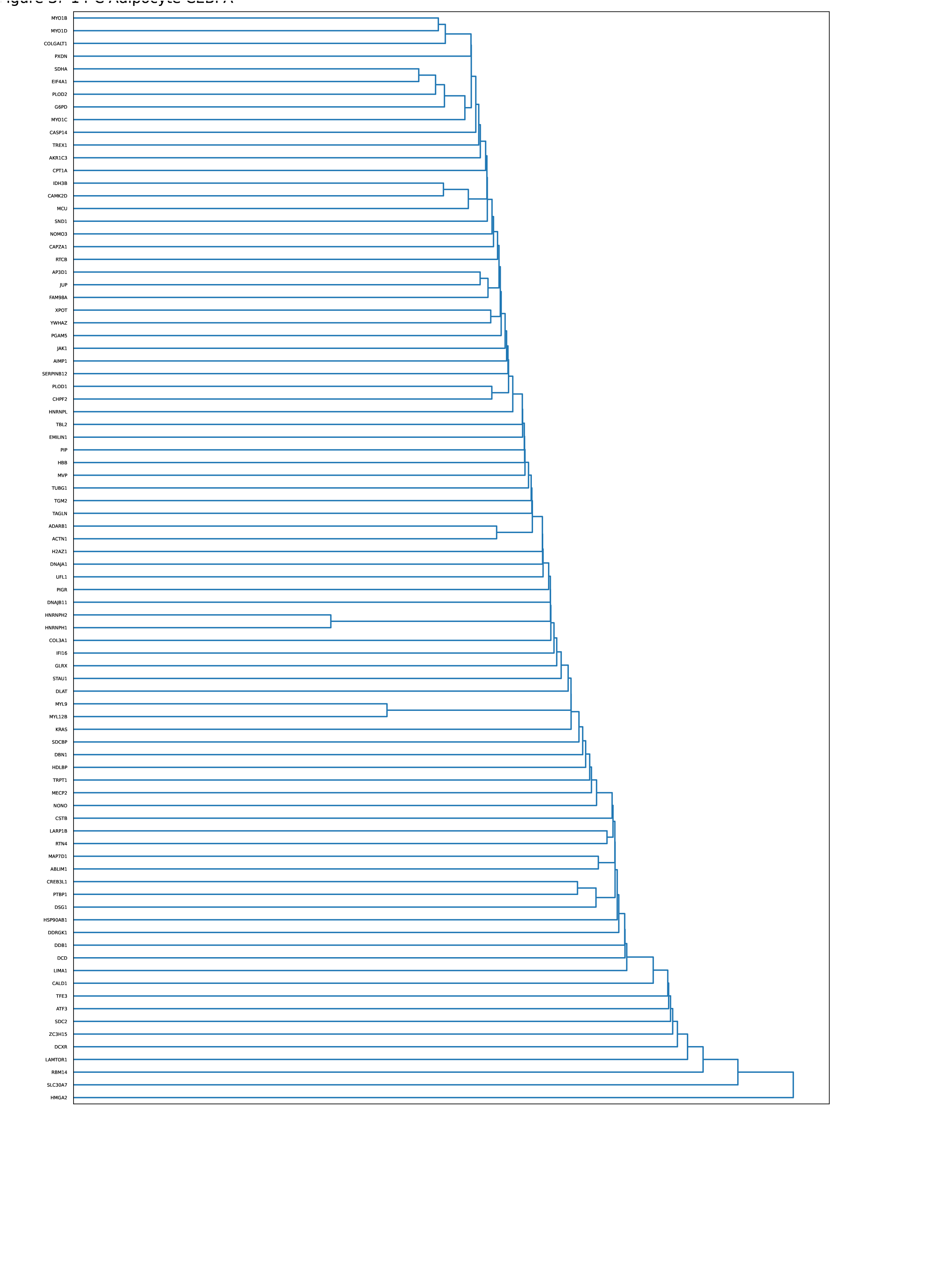
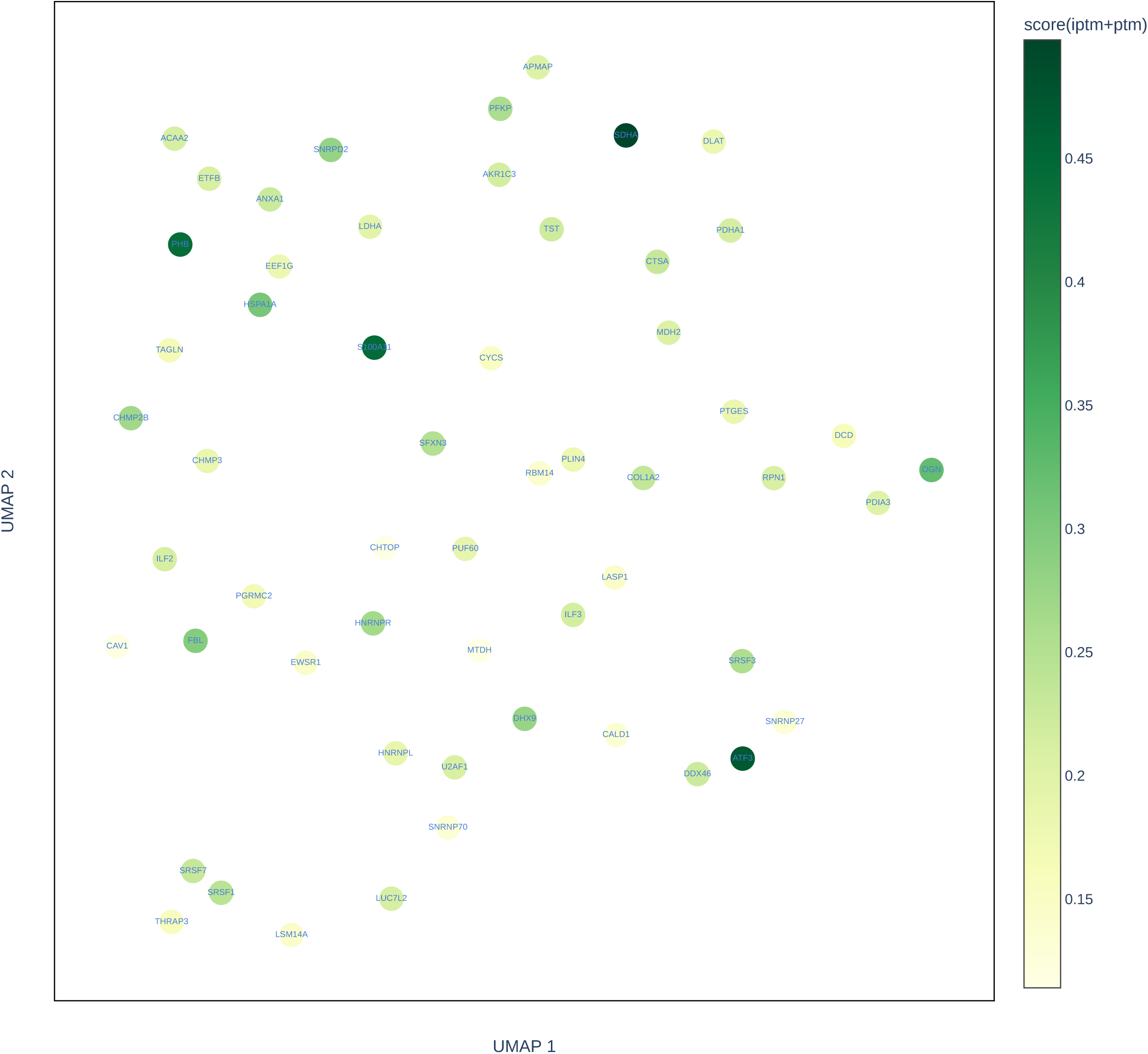
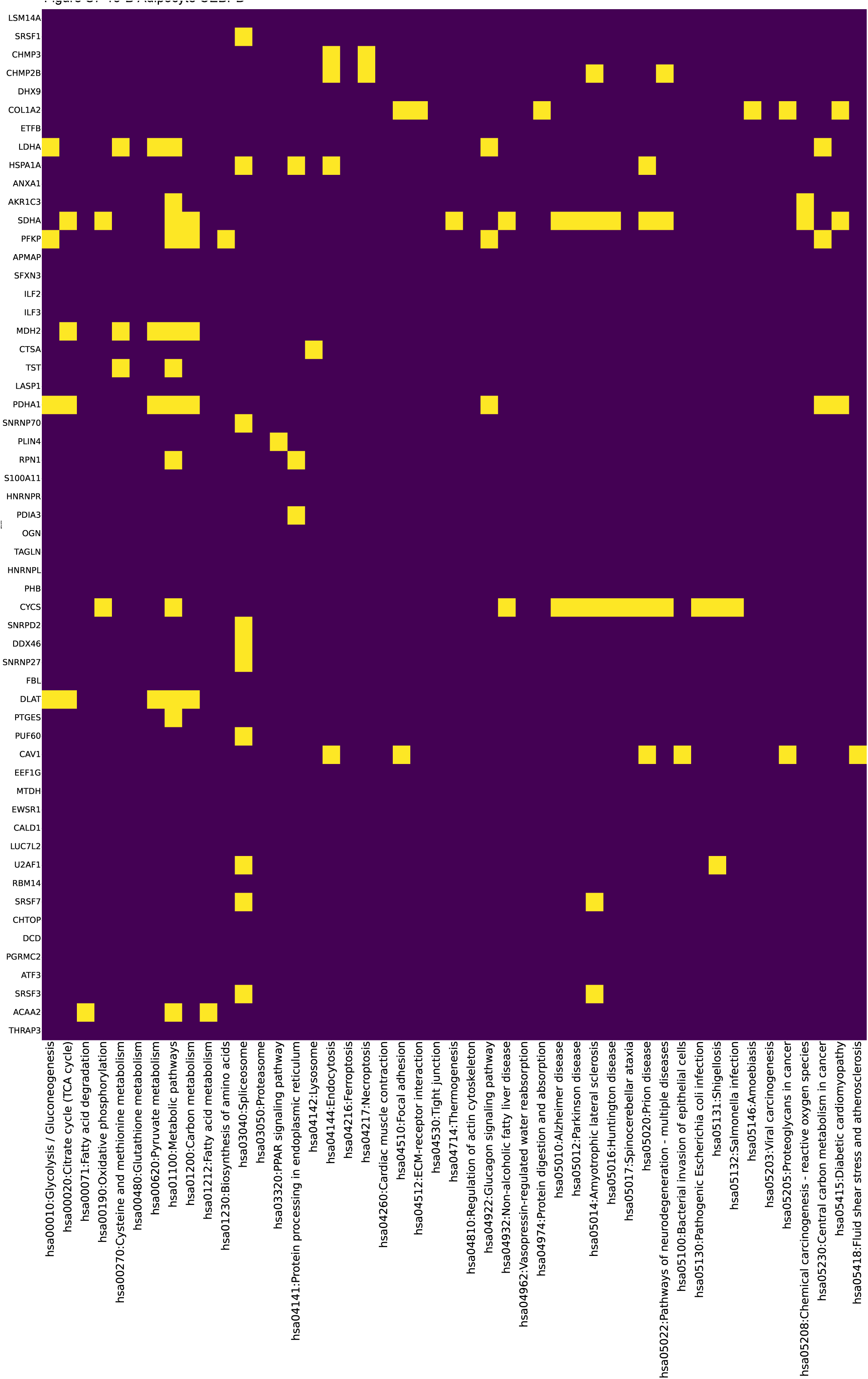
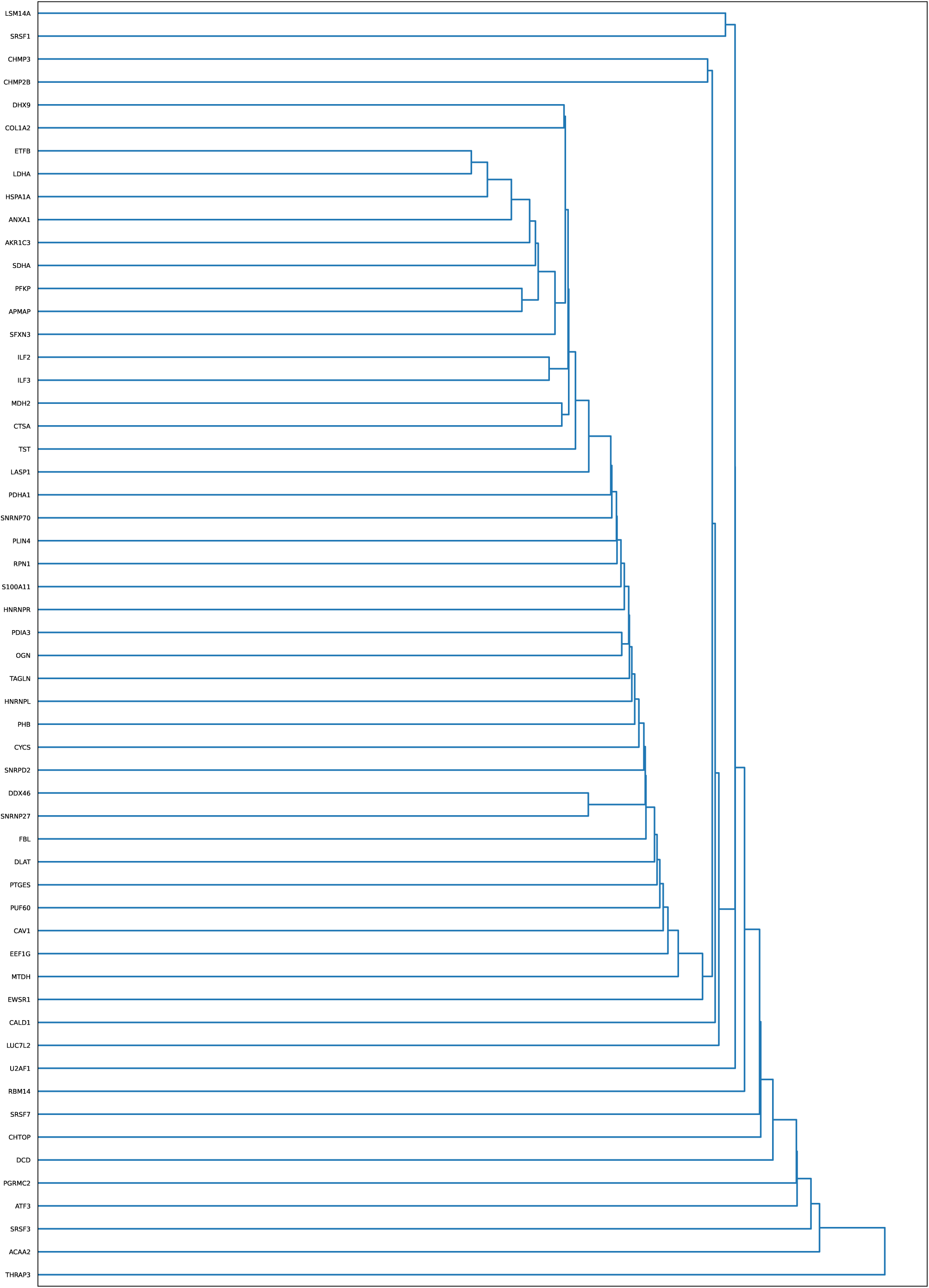
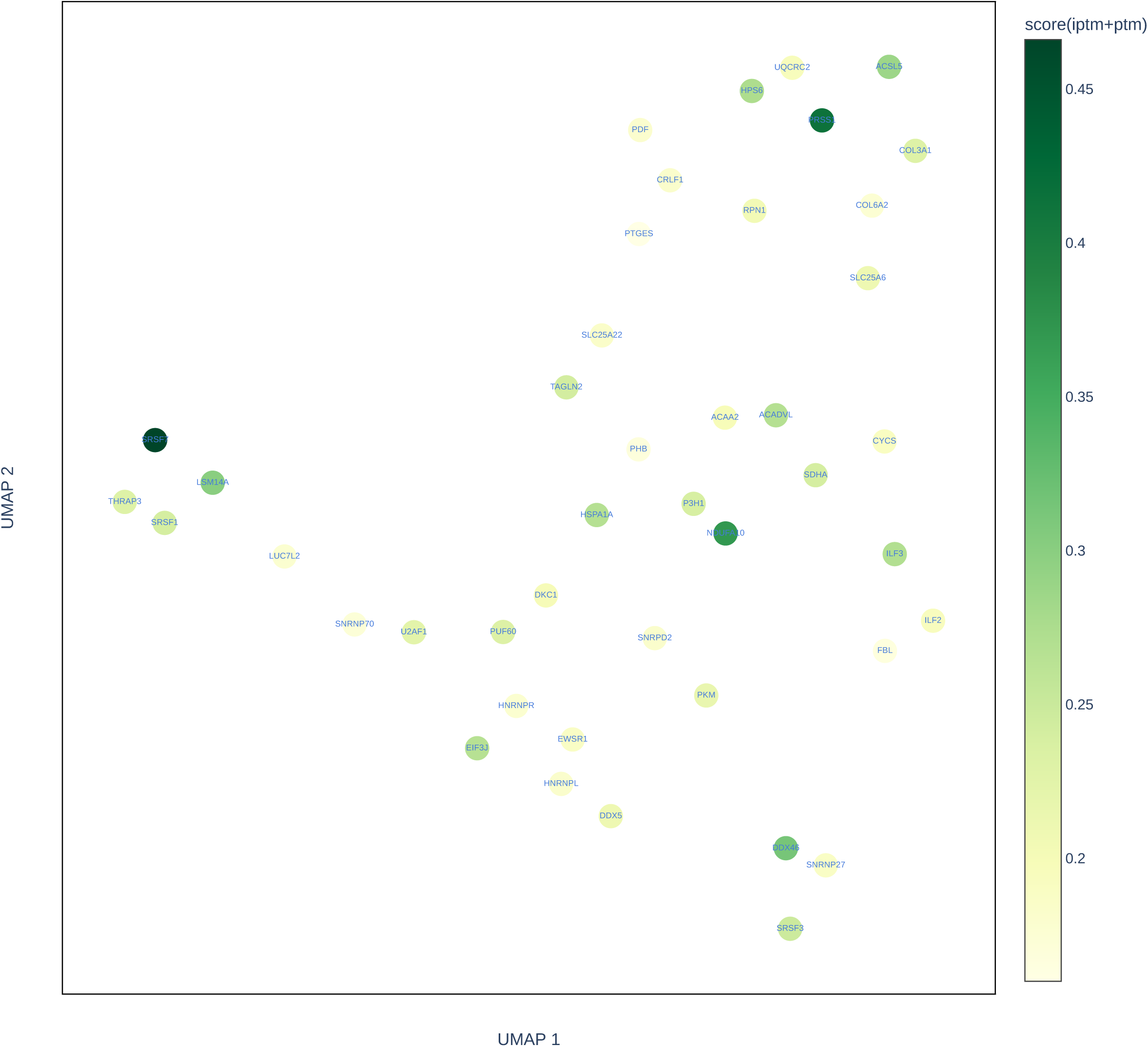
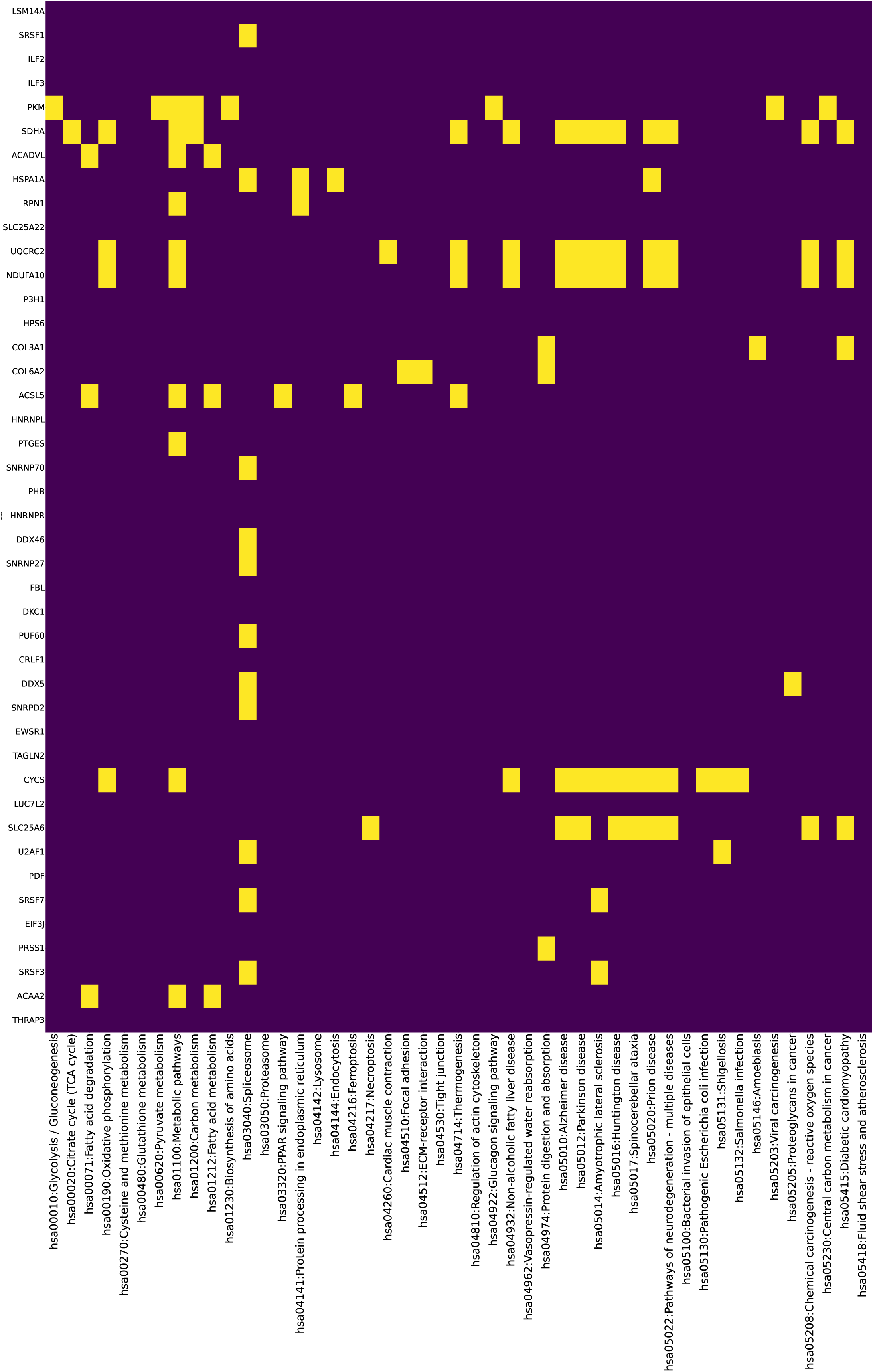
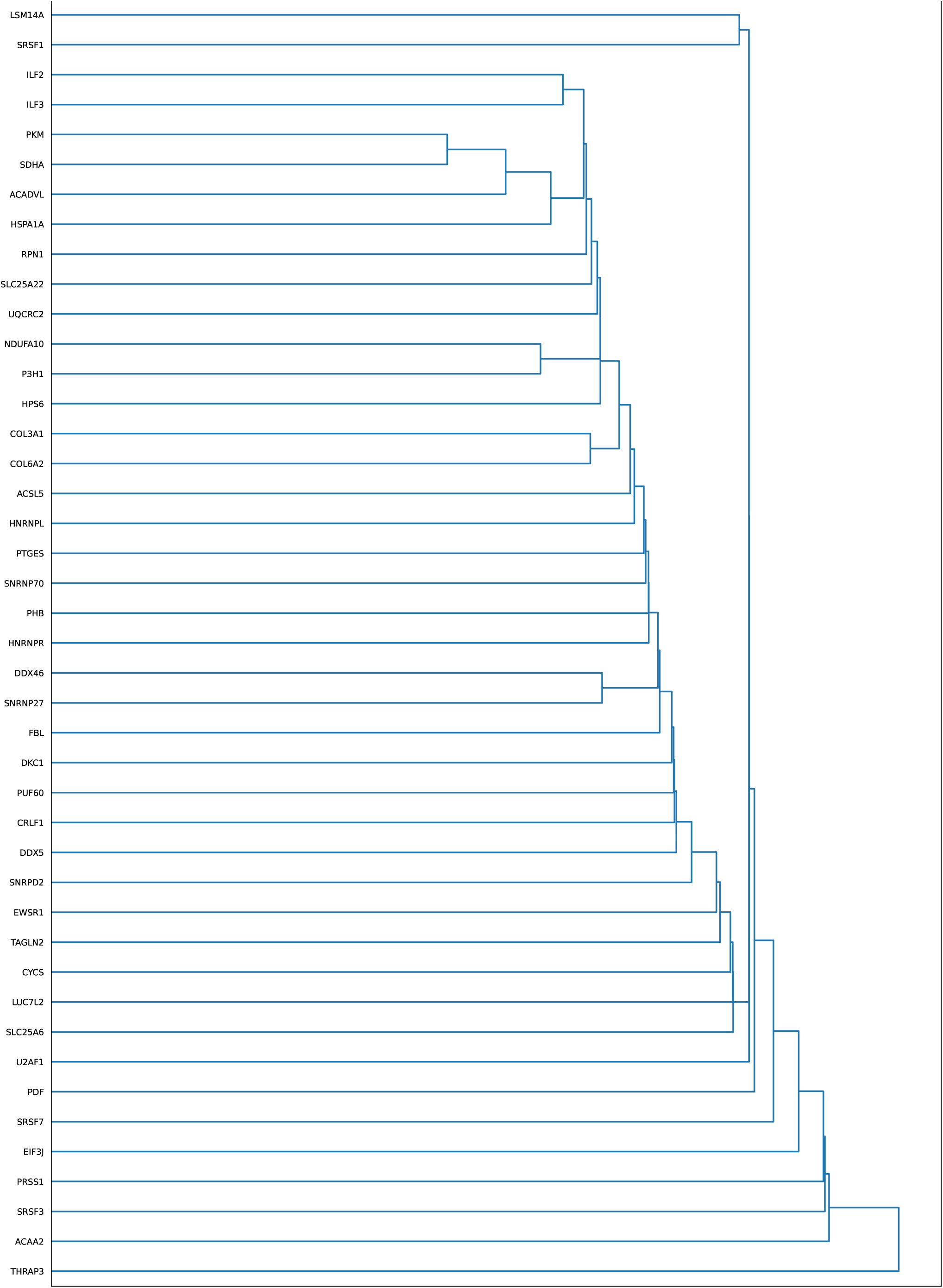
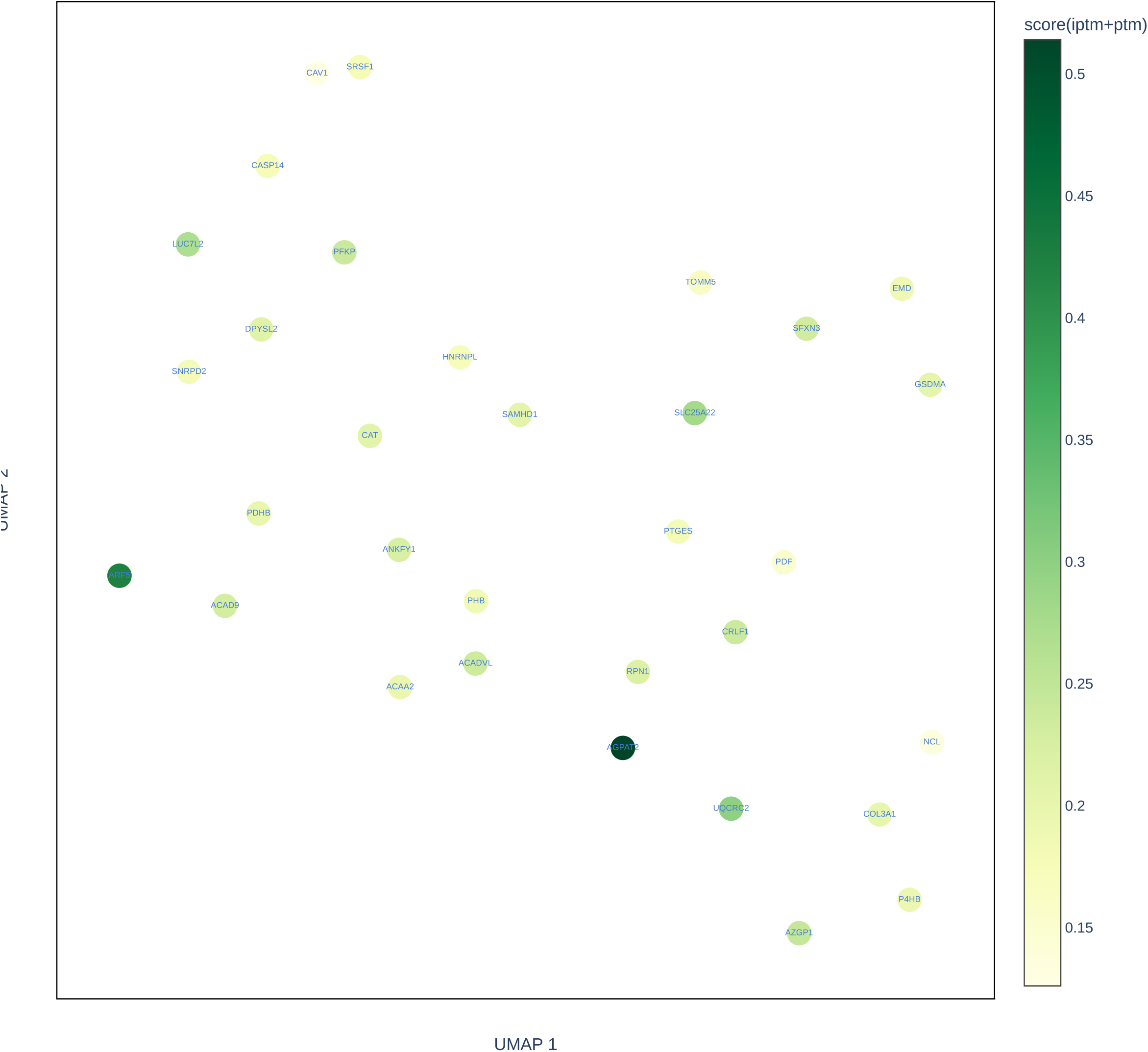
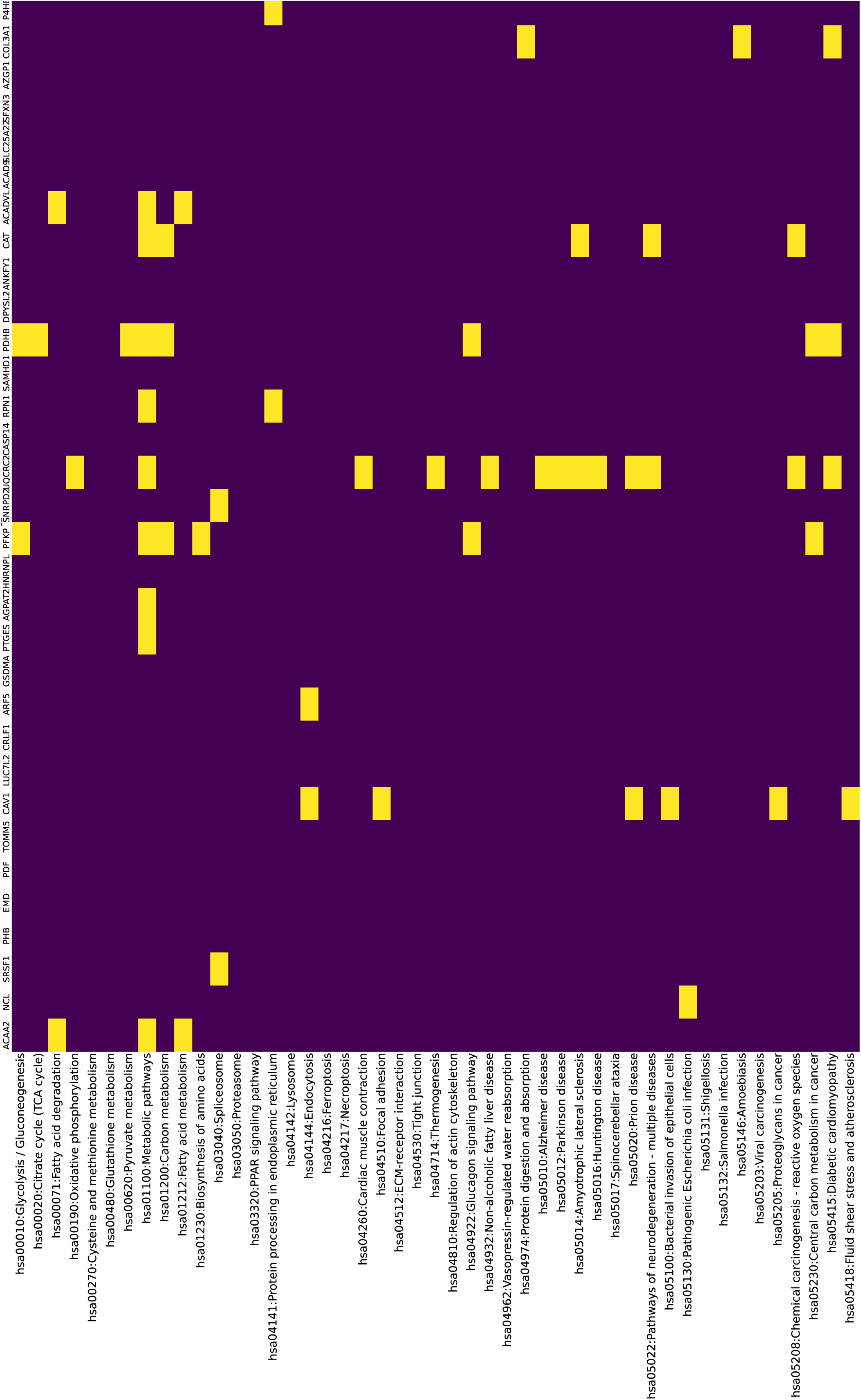
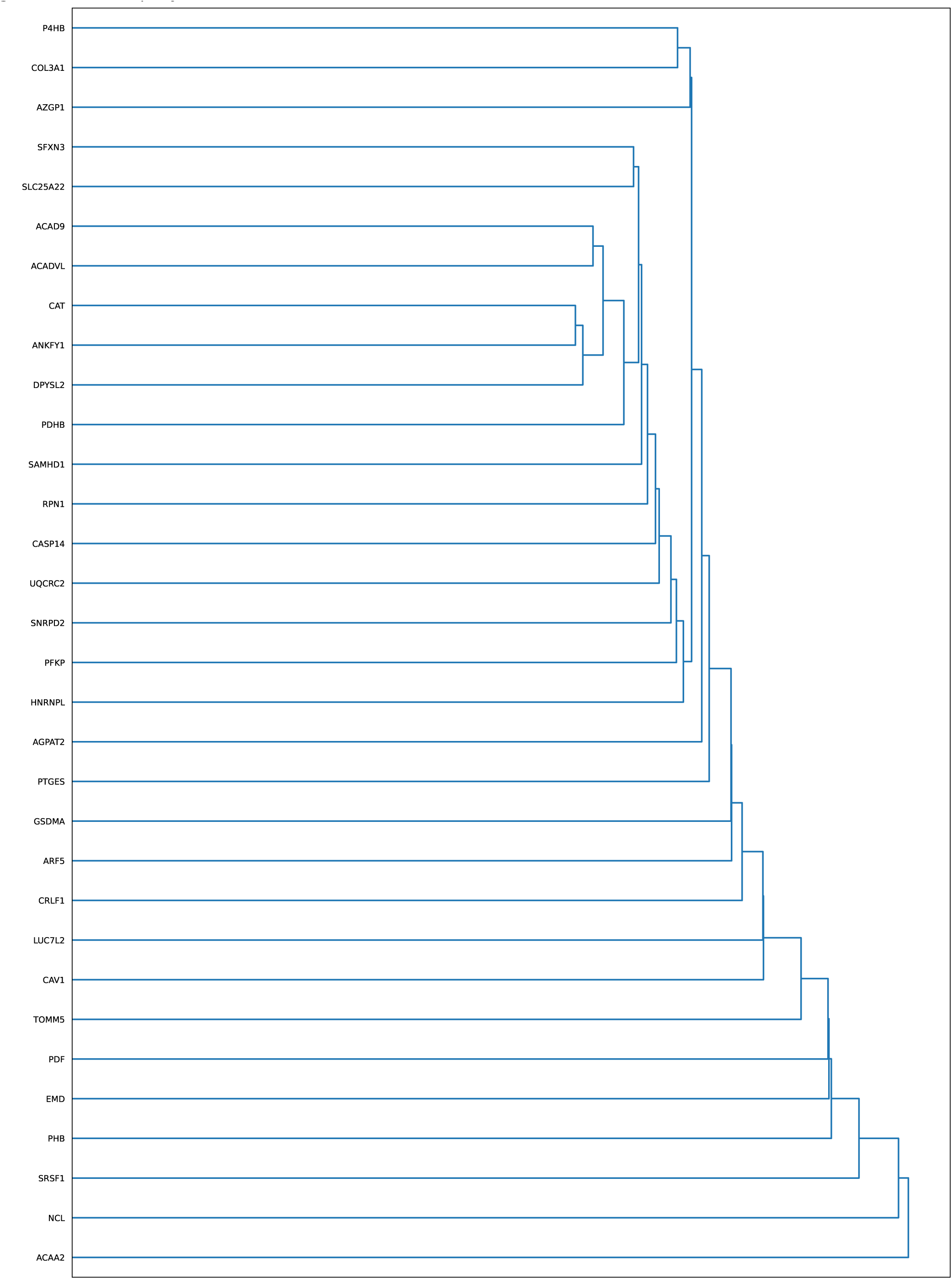
Related to Figure S8 and Supplementary Table S14. The relationship between interacting proteins in protein complexes. Each panel indicates proteins that specifically interact with bait proteins, such as FOS and GATA2. (A) 2D visualization using UMAP of proteins that specifically interact with bait proteins in adipocytes. Colors indicate CFM scores of iptm+ptm. The proteins with highly similar structures are mapped closely in UMAP at a short distance. (B, C) Hierarchical clustering of pathway terms and proteins interacting with bait proteins in an adipocyte-specific manner are detected by HaloMS. Interacting protein hits in the KEGG pathway are indicated in yellow. Hierarchical clustering distance is the similarity of latent representations of protein structure features in CFM.

**Figure S8.**
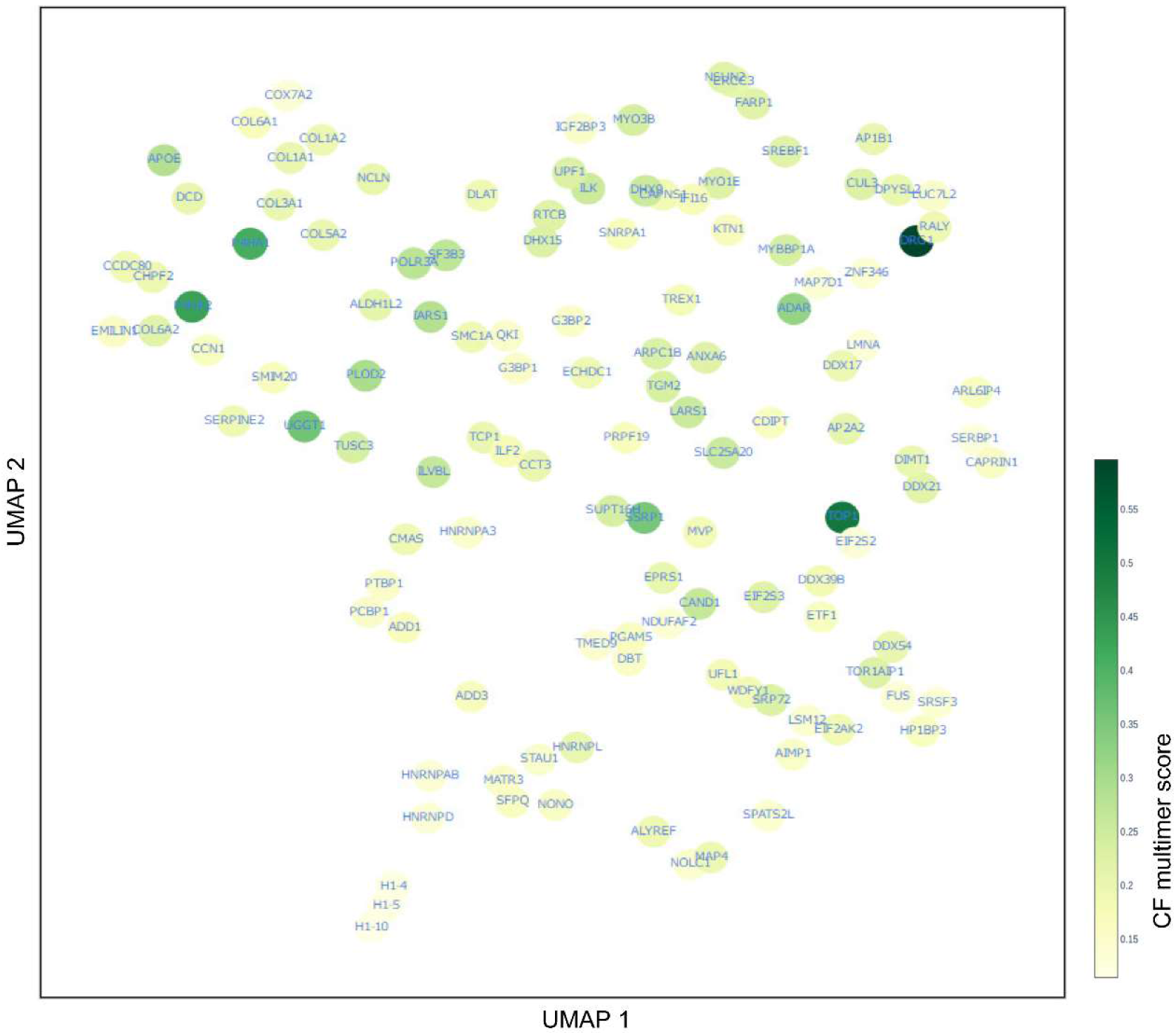
The relationship between interacting proteins in protein complexes. The 2D visualization using UMAP of proteins that specifically interact with GATA2 in adipocytes. Colors indicate CFM scores. Proteins that have closely related structures are mapped at close distances (Supplementary Table S14).

**Figure S9.**
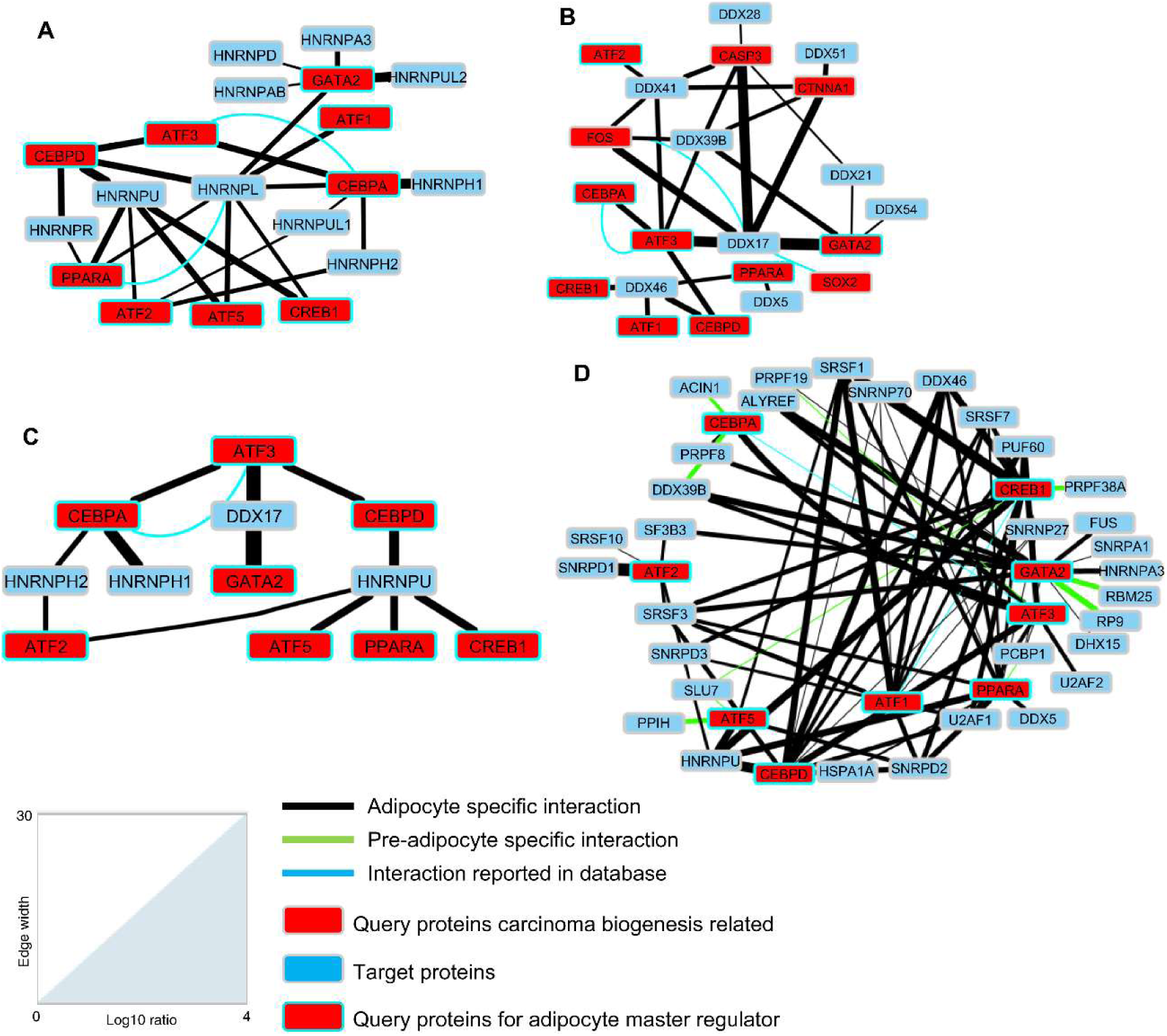
Four case studies from the TF-HaloMS dataset highlight the manner in which the human adipocyte master regulator may control and contribute to adipocyte signaling pathways. (A) Protein–protein interactions are detected in the TF-HaloMS dataset, which suggest that HNRNP proteins interact with nine master adipocyte regulators. (B) A subnetwork of ATP-dependent RNA helicases, DEAD box proteins, are involved in RNA metabolism, from transcription to degradation, based on TF-HaloMS and literature-curated interactions from BioGrid (https://thebiogrid.org/). DDX17 interacts with five bait proteins, including two adipocyte master regulators, GATA2 and ATF3, which act as connectors to other DDXs and adipocyte master regulators. (C) TF-HaloMS suggests cooperation between RNA processing and adipocyte signaling pathway, pointing to a possible connection between DDX17, HNRNPH1, HNRNPH2, and HNRNPU, and key transcription factors of adipocyte biogenesis (CEBPA, CEBPD, and ATF3). These proteins may comprise the core network of adipocyte differentiation cooperating with lnRNA. (D) Network communities in the spliceosome pathway in adipocyte and pre-adipocyte. TF-HaloMS suggests an interplay between the spliceosome-associated proteins and the adipocyte master regulator. The thickness of the edge indicates the log_10_ ratio in the indicator (Supplementary Tables S28–31).

**Figure S10.**
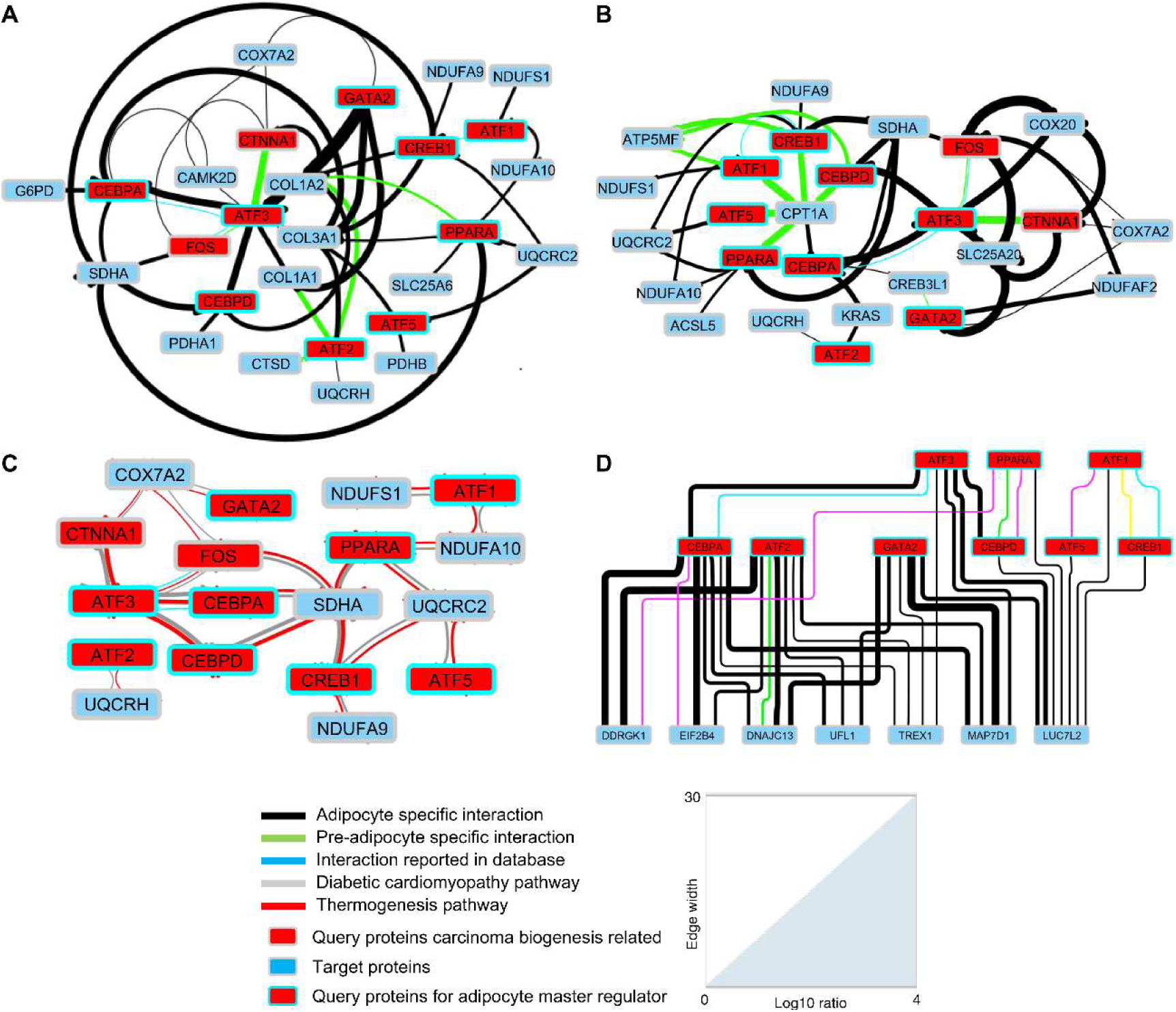
Related to Figure S4. (A) Diabetic cardiomyopathy and thermogenesis subnetworks within a dataset of adipocyte- and pre-adipocyte-specific interactions. The HaloMS interactions expand the known diabetic cardiomyopathy network of protein interactions. COL3A1 interacts with seven adipocyte regulatory molecules (ATF2, ATF3, ATF5, CEBPA, CREB1, GATA2, and PPARA). The protein pairs of the pathway are listed in Supplementary Table S32. (B) Network communities in the thermogenesis pathway. Protein pairs of the pathway are listed in Supplementary Table S32. (C) HaloMS dataset suggests an interplay between diabetic cardiomyopathy and thermogenesis pathways. Thirty percent of interaction in the diabetic cardiomyopathy subnetwork in Figure S7A overlaps with the thermogenesis subnetwork in Figure S7B composed of mitochondrial proteins. Protein pairs of the pathway are listed in Supplementary Table S32. (D) The link between ER-phagy, neurodegenerative disease, and adipocyte. The subnetwork that is not mapped by the pathway analysis is shown in Figure S4. The network includes proteins involved in ER-phagy, neurodegenerative disease, and myelodysplastic and master regulators in adipocytes. Protein pairs of the pathway are listed in Supplementary Table S33. The thickness of the edge indicates the log_10_ ratio in the indicator.

Supplementary Tables S1 to S33, doi: 10.17632/9wsrx97mg8.1 (link to the preview; https://data.mendeley.com/datasets/9wsrx97mg8/draft?a=c3f1d528-df61-4505-b6b9-c0b5e952f638)

Supplementary Table S1. Related to Figure 1. Query proteins are expressed from five ORF clones of *Arabidopsis* and consistently captured.

The protein identification threshold is set at both peptide and protein false-discovery rates of less than 1%. Intensity and quantitative protein values are calculated from MS data.

Supplementary Table S2. Related to Figure 1. Proteins from cell lysate of *Arabidopsis* leaves (9999 proteins).

Number, continuation number of the identified protein list. Accession number, the TAIR accession number of The *Arabidopsis* Information Resource (https://www.arabidopsis.org/). Protein name, the name of protein. # of grouped proteins: 1, a unique protein; 2 or more grouped proteins. Molecular weight, protein molecular weight. Peptide count, the number of peptide fragments used for protein identification, including the number of peptides shared with other proteins. Unique peptide count, the number of peptide fragments with unique peptide sequences in a protein (protein group). Protein group score, the maximum of the peptide probabilities of the peptides which are exclusive to the protein group. Intensity, quantitative protein value calculated from MS data. AtID, locus name of the protein in the *Arabidopsis* Information Resource.

Supplementary Table S3. Related to Figure 1. Protein–protein interaction network containing 740 interactions and 306 proteins is obtained after probing *Arabidopsis* leaf tissue lysate with five transcription factors.

OL, the overlap between HaloMS and protein array (reference *15*) or BioGrid database (https://thebiogrid.org/, ORGANISM-Arabidopsis_thaliana_Columbia-4.4.211.tab3).

Supplementary Table S4. Related to Figure 1. Gene ontology term enrichment of the HaloMS dataset from *Arabidopsis*.

Gene ontology annotation of protein interactions from *Arabidopsis* is performed at the TAIR site powered by PANTHER. This tool can be used to identify Gene Ontology terms that are over- or under-represented in a gene set. Data are then sent to the PANTHER classification system, which contains the latest GO annotation data for *Arabidopsis* and other plant species. The fold-enrichment and enrichment *P*-values are shown.

Supplementary Table S5. Related to Figure 1. Gene ontology term enrichment of protein microarray dataset from *Arabidopsis* (reference *15*).

Gene ontology annotation of protein interactions from *Arabidopsis* is performed at the TAIR site powered by PANTHER. This tool can be used to identify Gene Ontology terms that are over- or under-represented in a gene set. Data are then sent to the PANTHER classification system, which contains the latest GO annotation data for *Arabidopsis* and other plant species. The columns contain the fold-enrichment and the P-value for enrichment.

Supplementary Table S6. Related to Figure 1. Proteins from Hela and HEK293F cell lysates (7130 proteins).

Number, the sequence number of the identified protein list. Uniprot Ac, the Uniprot accession number (https://www.uniprot.org/). Protein name, the name of protein. # of grouped proteins: 1, a unique protein; and 2 or more, grouped proteins. Species, the species of origin. Molecular weight, the molecular weight of proteins. Identified peptide count, the number of peptide fragments used for protein identification, including the number of peptides shared with other proteins. Total unique peptide count, the number of peptide fragments with unique peptide sequences in a protein (protein group). Protein group score, the score for the accuracy of protein identification. Intensity, quantitative protein values calculated from MS data.

Supplementary Table S7. Related to Figure 2. Protein–protein interaction network containing 3302 interactions and 913 proteins, is revealed by HaloMS after probing a lysate of human cultured cells (HeLa and HEK293F cells) with 16 human proteins.

BioGrid OL, the overlap between HaloMS and BioGrid database data (https://thebiogrid.org/BIOGRID-ORGANISM-Homo_sapiens-4.4.211.tab3.txt).

Supplementary Table S8. Related to Figure 2A, 2B. Evaluation of the quality of the TF-HaloMS dataset by *in vitro* pull-down assay.

Gels, gel ID from an SDS-PAGE/western blot. Proteins, the Halo-tagged and/or 3×HA-tagged proteins are detected using a HaloTag ligand-TMR (odd-numbered gels) or an anti-HA antibody (even-numbered gels) in the “Detection” column. Classes, input, supernatant, or immunoprecipitation (IP) proteins.

Supplementary Table S9. Related to Figure 2C, 2D. Prediction of protein–protein interactions in the TF-HaloMS dataset by AlphaFold-Multimer.

Class, the types of protein–protein interactions in afmPRS, afmRRS, and HaloMS. Best score (iptm+ptm), the highest prediction score among the five scores (models 1 to 5) are obtained by AFM. Mean, the average of the five scores for model confidence. SD, the standard deviation for the five scores (models 1 to 5).

Supplementary Table S10. Related to Figure 2E. Digital pull-down assay of protein–protein interactions in the TF-HaloMS dataset is predicted using AlphaFold-Multimer.

The average of five scores for model confidence obtained by AFM is used to determine the significance of differences between each protein pair and the HaloTag-only control pairs, as in the pull-down assay. Classes, the types of protein–protein interactions in afmPRS, afmRRS, and HaloMS. Best score (iptm+ptm), the highest prediction score among the five scores (models 1–5) are obtained by AFM. Mean, the average of the five scores for model confidence. SD, the standard deviation for the five scores (models 1 to 5). *P*-value and Significance, the significance of differences between each protein pair and the HaloTag-only control pairs.

Supplementary Table S11. Related to Figure 3. Proteins from lysates of four different human cell lines.

Of 9405 proteins detected, 7258 are found in adipocytes, 7552 in HEK293F cells, 8149 in HeLa cells, and 8225 in pre-adipocytes. Each cell-line-specific column indicates quantitative protein values (log_10_) calculated from MS data.

Supplementary Table S12. Related to Figure 3. The 96 query proteins are expressed and consistently captured from 17 human ORF clones.

The protein identification threshold is set at both peptide and protein false-discovery rates of less than 1%. Query protein name, the query protein name is indicated along the x-axis in Figure S8. Cell type, the type of cell (adipocyte, pre-adipocyte, HeLa, and HEK293F) is probed with each query protein using HaloMS. Intensity (log_10_), quantitative protein values of HaloTag are calculated from MS data. Plate, type of plates that are used for HaloMS screening.

Supplementary Table S13. Related to Figure 3A. TF-HaloMS dataset containing 6512 interactions and 1690 proteins, is generated by HaloMS screening of four human cells using 17 human transcription factors.

Database, the overlap between HaloMS and literature-curated interactions in BioGrid database (https://thebiogrid.org/).

Supplementary Table S14. Related to Figure 3C, 3D, Supplementary Figures S7 and S8. Prediction of protein–protein interactions for adipocyte- and pre-adipocyte-specific interactions from HaloMS data set by ColabFold.

Class, the type of protein–protein interactions in adipocyte, pre-adipocyte, cfPRS, and cfPRS-Halo. ColabFold score, prediction score by artificial intelligence. Database, the overlap between HaloMS dataset and literature-curated interactions in BioGrid database (https://thebiogrid.org/).

Supplementary Table S15. Related to Supplementary Figure S4. Every cell-specific interaction protein is detected, enriched, and classified by pathway analysis.

DAVID and Heatmapper are used for the analysis (https://david.ncifcrf.gov/, http://www.heatmapper.ca/expression/). A z-score in the Supplementary Table is obtained from the *P*-values of the enriched classes. Higher z-scores indicate the presence of cell-specific subnetworks. Samples that are not enriched in a pathway and for which no *P*-value is obtained have an assigned *P*-value (–log_10_) of 0. The list of cell-specific interacting proteins used in this analysis is derived from 1428 (HeLa), 1201 (adipocyte), 1357 (HEK), and 1110 (pre-adipocyte) in Supplementary Figure S4.

Supplementary Table S16. Related to Supplementary Figure S4. Pathway-analyzed cell-specific interacting proteins.

DAVID is used for the analysis (https://david.ncifcrf.gov/).

Supplementary Table S17. Related to Supplementary Figures S4 and S5A. Protein–protein interaction subnetwork of mitochondria-associated neurodegenerative disease pathways.

This list includes amyotrophic lateral sclerosis, Alzheimer’s, Parkinson’s, Huntington’s, and prion disease pathway genes from Supplementary Figure S5A. Ratio (log_10_), quantitative protein values of HaloTag are calculated from MS data. An interaction is scored as positive when the average ratio from multiple datasets is more than 1 SD above the median of the ratio obtained only using the 33-kDa HaloTag protein as the negative control. Pairs with a negative control of 0, no ratio available, and literature-curated interaction in the BioGrid database (thebiogrid.org) are assigned a ratio of 0.1.

Supplementary Table S18. Related to Supplementary Figures S4 and S5B.

Protein–protein interaction subnetwork of PPAR signaling pathway is shown in Supplementary Figure S5B. Ratio (log_10_), quantitative protein values of HaloTag are calculated from MS data. An interaction is scored as positive when the average ratio from multiple datasets is more than 1 SD above the median of the ratio obtained only using the 33-kDa HaloTag protein as the negative control. Pairs with a negative control of 0 and no ratio available, and literature-curated interaction in the BioGrid database (thebiogrid.org) are assigned a ratio of 0.1.

Supplementary Table S19. Related to Supplementary Figures S4 and S5C. Protein–protein interaction subnetwork of the non-alcoholic fatty liver disease pathway is shown in Supplementary Figure S5C. Ratio (log_10_), quantitative protein values of HaloTag are calculated from MS data.

An interaction is scored as positive when the average ratio from multiple datasets is more than 1 SD above the median of the ratio obtained only using the 33-kDa HaloTag protein as the negative control. Pairs with a negative control of 0 and no ratio available, and literature-curated interaction in the BioGrid database (thebiogrid.org) are assigned a ratio of 0.1.

Supplementary Table S20. Related to Supplementary Figures S4 and S5D. Protein–protein interaction subnetwork of fluid shear stress and the atherosclerosis pathway is shown in Supplementary Figure S5D.

Ratio (log_10_), quantitative protein values of HaloTag are calculated from MS data. An interaction is scored as positive when the average ratio from multiple datasets is more than 1 SD above the median of the ratio obtained only using the 33-kDa HaloTag protein as the negative control. Pairs with a negative control of 0 and no ratio available, and literature-curated interaction in the BioGrid database (thebiogrid.org) are assigned a ratio of 0.1.

Supplementary Table S21. Related to Supplementary Figures S4 and S6A. Protein–protein interaction subnetwork of the adrenergic signaling in cardiomyocytes pathway is shown in Supplementary Figure S6A. Ratio (log_10_), quantitative protein values of HaloTag are calculated from MS data.

An interaction is scored as positive when the average ratio from multiple datasets is more than 1 SD above the median of the ratio obtained only using the 33-kDa HaloTag protein as the negative control. Pairs with a negative control of 0 and no ratio available, and literature-curated interaction in the BioGrid database (thebiogrid.org) are assigned a ratio of 0.1.

Supplementary Table S22. Related to Supplementary Figures S4 and S6B. Protein–protein interaction subnetwork of the thyroid hormone synthesis pathway is shown in Supplementary Figure S6B.

Ratio (log_10_), quantitative protein values of HaloTag are calculated from MS data. An interaction is scored as positive when the average ratio from multiple datasets is more than 1 SD above the median of the ratio obtained only using the 33-kDa HaloTag protein as the negative control. Pairs with a negative control of 0 and no ratio available, and literature-curated interaction in the BioGrid database (thebiogrid.org) are assigned a ratio of 0.1.

Supplementary Table S23. Related to Supplementary Figures S4 and S6C. Protein–protein interaction subnetwork of the AMPK signaling pathway is shown in Supplementary Figure S6C. Ratio (log_10_), quantitative protein values of HaloTag are calculated from MS data. An interaction is scored as positive when the average ratio from multiple datasets is more than 1 SD above the median of the ratio obtained only using the 33-kDa HaloTag protein as the negative control. Pairs with a negative control of 0 and no ratio available, and literature-curated interaction in the BioGrid database (thebiogrid.org) are assigned a ratio of 0.1.

Supplementary Table S24. Related to Supplementary Figures S4 and S6D. Protein–protein interaction subnetwork of the platelet activation pathway is shown in Supplementary Figure S6D. Ratio (log_10_), quantitative protein values of HaloTag are calculated from MS data. An interaction is scored as positive when the average ratio from multiple datasets is more than 1 SD above the median of the ratio obtained only using the 33-kDa HaloTag protein as the negative control. Pairs with a negative control of 0 and no ratio available, and literature-curated interaction in the BioGrid database (thebiogrid.org) are assigned a ratio of 0.1.

Supplementary Table S25. Related to Figure S4 and Figure S6E. Protein–protein interaction subnetwork of the peroxisome pathway is shown in Figure S6E. Ratio (log_10_), quantitative protein values of HaloTag are calculated from MS data. An interaction is scored as positive when the average ratio from multiple datasets is more than 1 SD above the median of the ratio obtained only using the 33-kDa HaloTag protein as the negative control. Pairs with a negative control of 0 and no ratio available, and literature-curated interaction in the BioGrid database (thebiogrid.org) are assigned a ratio of 0.1.

Supplementary Table S26. Related to Supplementary Figures S4 and S6F. Protein–protein interaction subnetwork of N-glycan biosynthesis pathway is shown in Supplementary Figure S6F.

Ratio (log_10_), quantitative protein values of HaloTag are calculated from MS data. An interaction is scored as positive when the average ratio from multiple datasets is more than 1 SD above the median of the ratio obtained only using the 33-kDa HaloTag protein as the negative control. Pairs with a negative control of 0 and no ratio available, and literature-curated interaction in the BioGrid database (thebiogrid.org) are assigned a ratio of 0.1.

Supplementary Table S27. Related to Supplementary Figures S7–9BC and S8 and Supplementary Table S14. Protein list from hierarchical clustering between pathway terms and proteins interacting with GATA2 in an adipocyte-specific manner is detected by HaloMS.

Supplementary Table S28. Related to Supplementary Figure S9. Protein–protein interaction subnetwork of heterogeneous nuclear ribonucleoprotein in adipocyte is shown in Supplementary Figure S9A.

Ratio (log_10_), quantitative protein values of HaloTag are calculated from MS data. An interaction is scored as positive when the average ratio from multiple datasets is more than 1 SD above the median of the ratio obtained only using the 33-kDa HaloTag protein as the negative control. Pairs with a negative control of 0 and no ratio available, and literature-curated interaction in the BioGrid database (thebiogrid.org) are assigned a ratio of 0.1.

Supplementary Table S29. Related to Supplementary Figure S9. Protein–protein interaction subnetwork of ATP-dependent RNA helicases and DEAD box proteins involved RNA metabolism from transcription to degradation in adipocyte is shown in Supplementary Figure S9B.

Ratio (log_10_), quantitative protein values of HaloTag are calculated from MS data. An interaction is scored as positive when the average ratio from multiple datasets is more than 1 SD above the median of the ratio obtained only using the 33-kDa HaloTag protein as the negative control. Pairs with a negative control of 0 and no ratio available, and literature-curated interaction in the BioGrid database (thebiogrid.org) are assigned a ratio of 0.1.

Supplementary Table S30. Related to Supplementary Figure S9. Interactions among DDX17, HNRNPs (H1/H2/U), and key transcription factors of adipocyte biogenesis are shown in Supplementary Figure S9C. Ratio (log_10_), quantitative protein values of HaloTag are calculated from MS data.

An interaction is scored as positive when the average ratio from multiple datasets is more than 1 SD above the median of the ratio obtained only using the 33-kDa HaloTag protein as the negative control. Pairs with a negative control of 0 and no ratio available, and literature-curated interaction in the BioGrid database (thebiogrid.org) are assigned a ratio of 0.1.

Supplementary Table S31. Related to Supplementary Figure S9. Protein–protein interaction subnetwork of the spliceosome in adipocyte and pre-adipocyte is shown in Supplementary Figure S9D.

Ratio (log_10_), quantitative protein values of HaloTag are calculated from MS data. An interaction is scored as positive when the average ratio from multiple datasets is more than 1 SD above the median of the ratio obtained only using the 33-kDa HaloTag protein as the negative control. Pairs with a negative control of 0 and no ratio available, and literature-curated interaction in the BioGrid database (thebiogrid.org) are assigned a ratio of 0.1.

Supplementary Table S32. Related to Supplementary Figures S4 and S10. Protein–protein interaction subnetwork of diabetic cardiomyopathy (Supplementary Figure S10A) and thermogenesis (Supplementary Figure S10B and S10C) in adipocyte and pre-adipocyte. Ratio (log_10_), quantitative protein values of HaloTag are calculated from MS data.

An interaction is scored as positive when the average ratio from multiple datasets is more than 1 SD above the median of the ratio obtained only using the 33-kDa HaloTag protein as the negative control. Pairs with a negative control of 0 and no ratio available, and literature-curated interaction in the BioGrid database (thebiogrid.org) are assigned a ratio of 0.1.

Supplementary Table S33. Related to Supplementary Figures S4 and S10. Protein–protein interaction subnetwork of ER-phagy, neurodegenerative disease, and myelodysplastic pathway is shown in Supplementary Figure S10D. Ratio (log_10_), quantitative protein values of HaloTag are calculated from MS data.

An interaction is scored as positive when the average ratio from multiple datasets is more than 1 SD above the median of the ratio obtained only using the 33-kDa HaloTag protein as the negative control. Pairs with a negative control of 0 and no ratio available, and literature-curated interaction in the BioGrid database (thebiogrid.org) are assigned a ratio of 0.1.

